# Short RNA chaperones promote aggregation-resistant TDP-43 conformers to mitigate neurodegeneration

**DOI:** 10.1101/2024.12.14.628507

**Authors:** Katie E. Copley, Jocelyn C. Mauna, Helen Danielson, Marilyn Ngo, Longxin Xie, Ashleigh Smirnov, Matt Davis, Leland Mayne, Miriam Linsenmeier, Jack D. Rubien, Bede Portz, Bo Lim Lee, Hana M. Odeh, Martina Hallegger, Jernej Ule, Piera Pasinelli, Yan Poon, Nicolas L. Fawzi, Ben E. Black, Christopher J. Donnelly, Brigid K. Jensen, James Shorter

## Abstract

Aberrant aggregation of the prion-like, RNA-binding protein TDP-43 underlies several debilitating neurodegenerative proteinopathies, including amyotrophic lateral sclerosis (ALS). Here, we define how short, specific RNAs antagonize TDP-43 aggregation. Short, specific RNAs engage and stabilize the TDP-43 RNA-recognition motifs, which allosterically destabilizes a conserved helical region in the prion-like domain, thereby promoting aggregation-resistant conformers. By mining sequence space, we uncover short RNAs with enhanced activity against TDP-43 and diverse disease-linked variants. The solubilizing activity of enhanced short RNA chaperones corrects aberrant TDP-43 phenotypes in optogenetic models and ALS patient-derived neurons. Remarkably, an enhanced short RNA chaperone mitigates TDP-43 proteinopathy and neurodegeneration in mice. Our studies reveal mechanisms of short RNA chaperones and pave the way for the development of short RNA therapeutics for fatal TDP-43 proteinopathies.

## Main Text

There are no effective therapeutics for several fatal TDP-43 proteinopathies, including amyotrophic lateral sclerosis (ALS), frontotemporal dementia (FTD), limbic-predominant age-related TDP-43 encephalopathy (LATE), Alzheimer’s disease (AD), and chronic traumatic encephalopathy (CTE) (*1–8*). A unifying pathological feature of degenerating neurons in ∼97% of ALS cases (*3*), ∼45% of FTD cases (*3*), all LATE cases (*2*), ∼57% of AD cases (*9, 10*), and ∼85% of CTE cases (*7, 8*) is the aberrant cytoplasmic mislocalization and aggregation of TDP-43 (*11–18*). TDP-43 is an essential and predominantly nuclear RNA-binding protein with a prion-like domain (PrLD), which plays critical roles in numerous RNA processing modalities, including regulation of splicing, polyadenylation, and transcript stability (*19–25*).

Accumulating evidence suggests that an aberrant phase transition of TDP-43 in the cytoplasm is a key pathological event that is difficult for neurons to reverse (*16, 17, 26–32*). Deliverable agents that prevent and reverse the aberrant phase transitions of TDP-43 and restore functional TDP-43 to the nucleus in the degenerating neurons of ALS/FTD, LATE, AD, and CTE patients could provide an important therapeutic solution (*26, 31*). Indeed, such an agent would simultaneously eliminate any toxic gain-of-function of aberrant TDP-43 conformers in the cytoplasm and any toxic loss-of-function caused by depletion of TDP-43 from the nucleus (*26, 31*).

TDP-43 contains two RNA recognition motifs (RRMs), RRM1 and RRM2, which engage RNA and confer a preference for UG-rich binding motifs (Fig. 1A) (*24, 25, 33–36*). TDP-43 also contains an intrinsically disordered PrLD (Fig. 1A), which includes a short, conserved region (CR) with transient α-helical structure (*14, 37, 38*). The CR plays a pivotal role in TDP-43 phase separation and aggregation (*16, 17, 37–41*). Typically, wild-type (WT) TDP-43 aggregates in disease, but rare forms of disease are connected with TDP-43 missense variants (*42*). The PrLD harbors the vast majority of disease-linked mutations (*42*). Aberrant post-translational modifications (PTMs) of TDP-43 are prevalent in disease, including hyperphosphorylation and lysine acetylation (*15, 16, 43–47*). A therapeutic strategy that combats the broad spectrum of TDP-43 proteinopathies should be effective against each of these different forms of disease-relevant TDP-43.

**Fig. 1.**
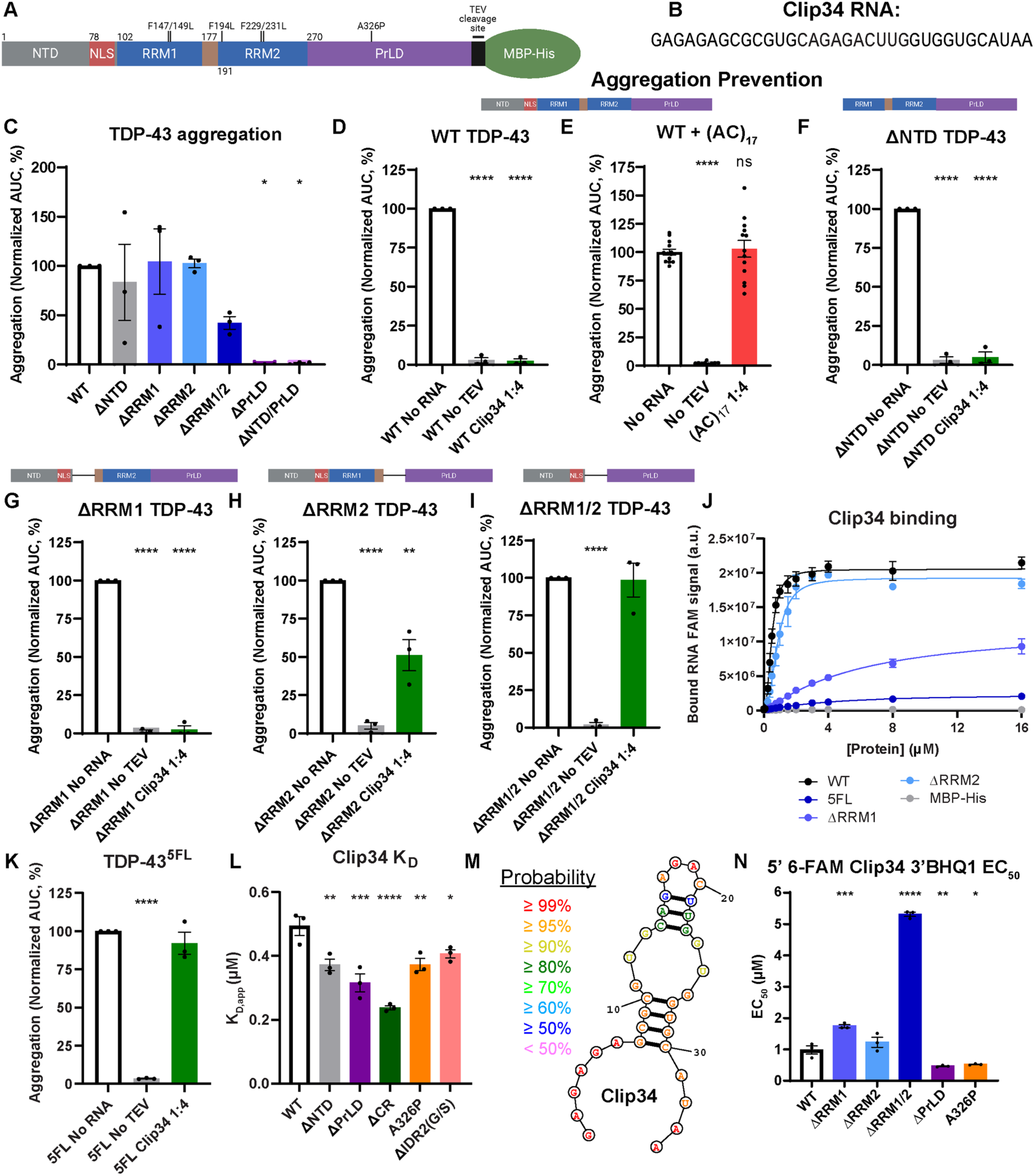
Clip34 is an allosteric antagonist of TDP-43 aggregation. (**A)** Domain map of TDP-43 indicating the locations of five phenylalanine-to-leucine mutations within RRM1 and RRM2, and the A326P mutation in the PrLD. **(B)** RNA sequence of Clip34. Clip34 is a 34nt RNA derived from the 3’ UTR of *TARDBP* RNA, to which TDP-43 binds to autoregulate its expression (*19, 33*). **(C)** Area under the curve (AUC) of standardized aggregation turbidity data for each TDP-43 deletion construct, normalized to WT TDP-43. Data are mean ± SEM (n=3; one-way ANOVA with Dunnett’s correction comparing to WT; *p ≤ 0.05). **(D-I)** AUC of turbidity data for each variant normalized to its respective No RNA control. The domain map for the construct used in each assay is shown above the graph. For (D) and (F to I), No RNA conditions are based on the same data as shown in (C). Data are mean ± SEM (n=3; n=13 for (E); one-way ANOVA with Dunnett’s correction comparing to No RNA; **p ≤ 0.01, ****p ≤ 0.0001). **(J)** Bound 5’ 6-FAM Clip34 signal for EMSAs performed with indicated TDP-43-MBP-His variants or MBP-His. Data are mean ± SEM (n=3 for TDP-43 variants; n=2 for MBP-His; shown is the nonlinear regression: [agonist] vs response with variable slope, of the combined replicates). **(K)** AUC of TDP-43^5FL^ turbidity data normalized to the No RNA control. Data are mean ± SEM (n=3; one-way ANOVA with Dunnett’s correction comparing to No RNA; ****p ≤ 0.0001). **(L)** Apparent K_D_ values calculated from the bound signal from individual replicates of EMSAs performed with 5’ 6-FAM Clip34 and WT TDP-43-MBP-His or the indicated TDP-43 variant. WT data shown is the same data as in (J). Data are mean ± SEM (n=3; one-way ANOVA with Dunnett’s correction comparing to WT; *p≤ 0.05; **p≤ 0.01; ***p ≤ 0.001; ****p ≤ 0.0001). **(M)** Secondary structure of Clip34 as predicted by RNAstructure. Text color indicates the probability for each nucleotide, as in the legend. **(N)** Half maximal effective concentration (EC_50_) values calculated from individual replicates of relative fluorescence intensity for 5’ 6-FAM Clip34 3’ BHQ1 with indicated TDP-43-MBP-His variants. Data are mean ± SEM (n=3; 100 nM RNA; one-way ANOVA with Dunnett’s correction comparing to WT; *p ≤ 0.05, **p ≤ 0.01, ***p ≤ 0.001, ****p ≤ 0.0001).

RNA has recently emerged as a potent solubilizing agent for TDP-43 *in vitro* (*28, 29, 48–52*). Indeed, one short 34-nucleotide (nt) RNA derived from the 3’UTR of the *TARDBP* mRNA, termed Clip34 (Table S1 and Fig. 1B), which TDP-43 binds to regulate its own expression, can prevent and reverse WT TDP-43 phase separation and aggregation *in vitro* and in engineered human cell lines (*28, 29, 50*). There has been great success in utilizing short oligonucleotides (e.g., antisense oligonucleotides [ASOs]) as therapeutics for patients with various diseases, including cases where the oligonucleotide must exert its effect in the brain (*53–57*). Thus, Clip34 and potentially other short RNA chaperones are strong candidates as therapeutic agents to treat TDP-43 proteinopathies.

Several critical barriers, however, limit our understanding of short RNA chaperones like Clip34, which restricts their development. First, it is not clear how short RNAs must engage TDP-43 to antagonize aberrant assembly. Second, it remains uncertain how short RNAs might alter TDP-43 structure to prevent aberrant TDP-43 assembly. Third, it is unclear whether short RNAs can prevent aggregation of diverse disease-linked TDP-43 variants, including missense mutants and TDP-43 bearing disease-linked PTMs. Fourth, it is not clear whether RNA sequences exist that have enhanced chaperone activity against TDP-43 beyond Clip34. Finally, it is not known whether short RNAs can mitigate aberrant TDP-43 phenotypes in patient-derived neurons or mouse models of disease.

Here, we address these pressing issues. We elucidate the mechanism by which specific, short RNAs must engage TDP-43 and alter its structure to antagonize aggregation. By exploring sequence space, we uncover short RNAs with enhanced chaperone activity against TDP-43 and diverse disease-linked variants. Importantly, enhanced short RNA chaperones correct aberrant TDP-43 phenotypes in optogenetic human cell models, ALS patient-derived neurons, and mice. Our studies reveal mechanistic aspects of short RNA chaperone activity and open a window for the development of short RNA therapeutics for several fatal TDP-43 proteinopathies.

### Clip34 is an allosteric antagonist of TDP-43 aggregation

First, we mapped the domains of TDP-43 that enable the chaperone activity of Clip34. Thus, we purified full-length TDP-43 and TDP-43 constructs lacking the N-terminal domain (TDP-43^ΔNTD^), RRM1 (TDP-43^ΔRRM1^), RRM2 (TDP-43^ΔRRM2^), RRM1 and RRM2 (TDP-43^ΔRRM1/2^), the PrLD (TDP-43^ΔPrLD^), or the NTD and PrLD (TDP-43^ΔNTD/PrLD^) with a C-terminal MBP tag (Fig. 1A and fig. S1, A and B). Specific removal of the MBP tag with TEV protease elicits rapid TDP-43 aggregation, whereas TDP-43 remains soluble if the tag is not removed (fig. S1C and Fig. 1, C to I) (*28, 58, 59*). Under our assembly conditions, upon removal of the MBP tag, TDP-43 aggregated robustly, as did TDP-43^ΔNTD^, TDP-43^ΔRRM1^, and TDP-43^ΔRRM2^ (Fig. 1C). As expected, TDP-43 proteins lacking the PrLD did not aggregate (Fig. 1C) (*41*). Unexpectedly, TDP-43^ΔRRM1/2^ exhibited reduced aggregation (Fig. 1C). This finding indicates that the RRMs contribute to the aggregation propensity of full-length TDP-43.

We next assessed the ability of Clip34 to antagonize the aggregation of these TDP-43 deletion constructs. At a 1:4 ratio of RNA to TDP-43, Clip34 abolished TDP-43 aggregation (Fig. 1D), indicating that substoichiometric levels of Clip34 suffice to potently inhibit TDP-43 aggregation (*28*). This effect was specific, as Clip34 was unable to reduce phase separation of FUS (fig. S1, D and E), another RNA-binding protein with a PrLD connected to ALS/FTD (*31*). Moreover, the UG-deficient RNA (AC)_17_, which does not bind TDP-43, was unable to inhibit TDP-43 aggregation (Fig. 1E). Thus, not any short RNA can antagonize TDP-43 aggregation, indicating that specific sequence features are required for chaperone activity.

Clip34 potently inhibited the aggregation of TDP-43^ΔNTD^ (Fig. 1F), indicating that Clip34 binding to the NTD is not required for inhibition. Clip34 also effectively inhibited TDP-43^ΔRRM1^ aggregation (Fig. 1G). By contrast, Clip34 exhibited reduced ability to antagonize aggregation of TDP-43^ΔRRM2^ (Fig. 1H). Thus, RRM2 plays an important role in enabling Clip34 to antagonize TDP-43 aggregation. Clip34 was unable to antagonize aggregation of TDP-43^ΔRRM1/2^ (Fig. 1I).

Furthermore, Clip34 was unable to antagonize the aberrant assembly of a construct containing only the TDP-43 PrLD (data not shown). Thus, Clip34 does not inhibit TDP-43 aggregation via direct interactions with the PrLD, which drives TDP-43 aggregation (*16, 17, 41*). Rather, Clip34 must engage the RRMs to antagonize TDP-43 aggregation. These results suggest that Clip34 binding to the TDP-43 RRMs elicits an allosteric effect on other domains of TDP-43, which precludes TDP-43 aggregation.

### Revealing allosteric crosstalk between the TDP-43 RRMs, PrLD, and RNA

To explore how Clip34 promotes aggregation-resistant TDP-43 conformers, we first explored how TDP-43 binds to Clip34. TDP-43 binds Clip34 cooperatively, with a hill slope (h) ∼2.4 and a *K_D_* of ∼0.49 µM (Fig. 1J and fig. S2, A to C). By contrast, Clip34 does not bind strongly to TDP-43^5FL^, which bears F147L, F149L, F194L, F229L, and F231L mutations in the RRMs that impair RNA binding (Fig. 1J and fig. S2B) (*34*). Indeed, Clip34 is unable to inhibit TDP-43^5FL^ aggregation (Fig. 1K) (*28*). Thus, Clip34 does not engage the NTD or PrLD to exert chaperone activity. Rather, Clip34 engages the RRMs to abrogate TDP-43 aggregation and must exert allosteric effects to inhibit aggregation that is driven by intermolecular contacts between PrLDs.

We next assessed the contribution of RRM1 and RRM2. TDP-43^ΔRRM1^ binding to Clip34 was reduced by ∼1.6-fold in terms of B_max_ (the maximum specific binding) compared to full-length TDP-43, whereas TDP-43^ΔRRM2^ binding was only reduced by ∼1.1-fold (Fig. 1J and fig. S2B). TDP-43^ΔRRM1^ binds Clip34 with reduced cooperativity (h∼1.2) and a *K_D_* of ∼6.4 µM, whereas TDP-43^ΔRRM2^ binds Clip34 cooperatively (h∼2.5) with a *K_D_* of ∼0.9 µM (Fig. 1J and fig. S2C). Thus, RRM1 makes a larger contribution to tight, cooperative binding of Clip34 than RRM2. Despite weaker binding, Clip34 effectively inhibited aggregation of TDP-43^ΔRRM1^, but was less effective against TDP-43^ΔRRM2^ (Fig. 1, G and H). Thus, binding affinity does not necessarily predict chaperone activity. Indeed, these findings suggest that Clip34 must engage TDP-43 in a specific manner to effectively prevent aggregation, consistent with an allosteric mode of action.

We next explored the role of the NTD in binding to Clip34. We found that the NTD negatively regulates Clip34 binding, as TDP-43^ΔNTD^ binds Clip34 with a *K_D_* of ∼0.37 µM, indicating an ∼1.3-fold increase in affinity compared to full-length TDP-43 (Fig. 1L and fig. S2D). This finding is consistent with previous work identifying an allosteric connection between the NTD and RRMs (*60*). However, while the NTD negatively regulates binding to Clip34 (Fig. 1L and fig. S2D), Clip34 effectively inhibited TDP-43^ΔNTD^ aggregation (Fig. 1F). Thus, the NTD is not required for Clip34 chaperone activity.

How does the interaction between Clip34 and the RRMs prevent intermolecular contacts between PrLDs that drive aggregation? We considered whether Clip34 binding to the RRMs might allosterically affect the PrLD. The PrLD drives TDP-43 aggregation (*16, 17, 41*), but in cells the PrLD promotes binding and regulation of a subset of RNA targets, which contain over 100nt-long binding regions composed of dispersed binding sequences, including the 3’UTR of the *TARDBP* mRNA (*59*). Most studies of TDP-43 binding to RNA at the pure protein level have employed the isolated RRMs and not the full-length protein (*35*), hence the mechanism by which the PrLD may impact RNA binding remains unclear. We thus assessed how the PrLD affects Clip34 binding by soluble TDP-43. Unexpectedly, we found that the PrLD negatively regulates Clip34 binding to the RRMs. Deletion of the PrLD enhanced binding to Clip34 (Fig. 1L and fig. S2D). TDP-43^ΔPrLD^ binds Clip34 more cooperatively (h∼2.7) with a *K_D_* of ∼0.32 µM, indicating an ∼1.5-fold increase in affinity compared to full-length TDP-43 (Fig. 1L and fig. S2D). Hence, in addition to directly establishing intermolecular contacts that drive TDP-43 aggregation (*16, 17, 41*), the PrLD also indirectly promotes TDP-43 insolubility by reducing the apparent affinity of the RRMs for RNA (*29*). Indeed, RNA-binding deficient TDP-43 is highly aggregation-prone in cells (*29, 49, 61*).

To determine whether this inhibitory effect stems from a specific portion of the PrLD, we tested TDP-43 variants with specific deletions within the PrLD (*59*). Deletion of the extreme C-terminal portion of the PrLD (TDP-43^ΔIDR2(G/S)^) slightly enhanced binding to Clip34, with an ∼1.2-fold increase in affinity compared to full-length TDP-43 (Fig. 1L and fig. S2, D and E). Strikingly, deletion of the α-helical conserved region (CR) of the PrLD strongly enhanced binding to Clip34, indicated by an ∼2.1-fold increase in affinity for TDP-43^ΔCR^ compared to full-length TDP-43 (Fig. 1L and fig. S2, D and E). A helix-breaking mutation within the CR, TDP-43^A326P^, also enhanced binding to Clip34 (Fig. 1L and fig. S2D). Thus, negative regulation of RNA binding by the PrLD appears to be mediated, at least in part, by the α-helicity of the CR, a region that is critical for TDP-43 phase separation via helix-helix interactions, and aggregation via intermolecular β-sheet interactions (*16, 17, 32, 59, 62*).

To explore whether the inhibitory effect of the PrLD on RNA binding impacts the ability of Clip34 to prevent TDP-43 aggregation, we assessed the ability of Clip34 to antagonize aggregation of TDP-43 variants with specific deletions within the PrLD: TDP-43^ΔIDR1^, TDP-43^ΔCR^, TDP-43^ΔCR/IDR2(Q/N)^, TDP-43^ΔIDR2(G/S)^, and TDP-43^ΔCR/IDR2^ (fig. S3, A to I). As anticipated, TDP-43^ΔCR^ and TDP-43^ΔCR/IDR2^ exhibited reduced aggregation, whereas TDP-43^ΔIDR1^, TDP-43^ΔCR/IDR2(Q/N)^, and TDP-43^ΔIDR2(G/S)^ aggregated to a similar extent as full-length TDP-43 (fig. S3B) (*59*). Notably, Clip34 exhibited an enhanced ability to antagonize aggregation of these partial PrLD deletion variants (fig. S3, C to I). Thus, the PrLD antagonizes the ability of Clip34 to reduce TDP-43 aggregation. Collectively, these findings reveal allosteric crosstalk between the TDP-43 RRMs, PrLD, and RNA, which regulates the balance of soluble and aggregation-prone forms of TDP-43.

### TDP-43 binding remodels Clip34 by unfolding stem-loop structure

The TDP-43 RRMs engage single-stranded RNA often found in introns (*24, 25, 35*). However, Clip34 is predicted to form a stem-loop structure (Table S1 and Fig. 1M) (*63*). To assess this prediction, we utilized Clip34 RNA with a 5’ fluorophore and a 3’ quencher, based on the structural prediction that the ends of Clip34 are likely to be in close proximity (Fig. 1M) (*63*). Very low fluorescent signal was measured for the Clip34 fluorophore-quencher RNA on its own, indicating that the 5’ and 3’ ends are in close proximity, consistent with a stem-loop structure (Fig. 1N and fig. S2F). However, TDP-43 cooperatively binds to Clip34 (Fig. 1J), which may enable unfolding of the stem-loop structure. Indeed, upon addition of TDP-43, fluorescence increased strongly (Fig. 1N and fig. S2F), indicating that TDP-43 remodels Clip34 in a manner that increases the distance between the 5’ and 3’ ends. Deletion of RRM1 slightly impaired remodeling, whereas deletion of both RRMs strongly impaired remodeling (Fig. 1N and fig. S2F). Conversely, deletion of the PrLD or the CR helix-breaking TDP-43^A326P^ mutation enhanced remodeling (Fig. 1N and fig. S2F). Thus, TDP-43 remodels the stem-loop structure of Clip34 in a manner that depends on the RRMs and is negatively regulated by the PrLD. These findings made us wonder whether the energetics of the TDP-43:Clip34 interaction might alter the structural dynamics of TDP-43.

### Clip34 binding remodels TDP-43 by stabilizing the RRMs and destabilizing the PrLD CR

We next examined how Clip34 affected TDP-43 native structure via hydrogen/deuterium-exchange mass spectrometry (HXMS). HXMS measures the exchange of backbone amide hydrogen atoms over time after dilution in D_2_O-based buffer (*64*). When backbone hydrogens make hydrogen bonds, they exchange more slowly with deuterium (*64*). As backbone hydrogens make hydrogen bonds involved in protein secondary and tertiary structure, the kinetics by which the hydrogens exchange reports on the stability of structure in that region of the protein (*64*). In this way, we can establish how Clip34 might alter TDP-43 structural dynamics to preclude aggregation.

We performed HXMS across a wide timescale (1 s-14.5 h) with TDP-43 (with a C-terminal MBP tag to ensure solubility) in the presence or absence of excess Clip34 to saturate binding (Table S2 and fig. S4, A to C). We achieved 87.9% or greater coverage of the TDP-43 sequence for all conditions (Table S2 and fig. S4, A and B).The percentage difference in exchange between Clip34-bound and free states was calculated for each peptide at each timepoint, based on which consensus values for the percentage difference in exchange were calculated for each TDP-43 amino acid (Fig. 2, A and B, and fig. S5).

**Fig. 2.**
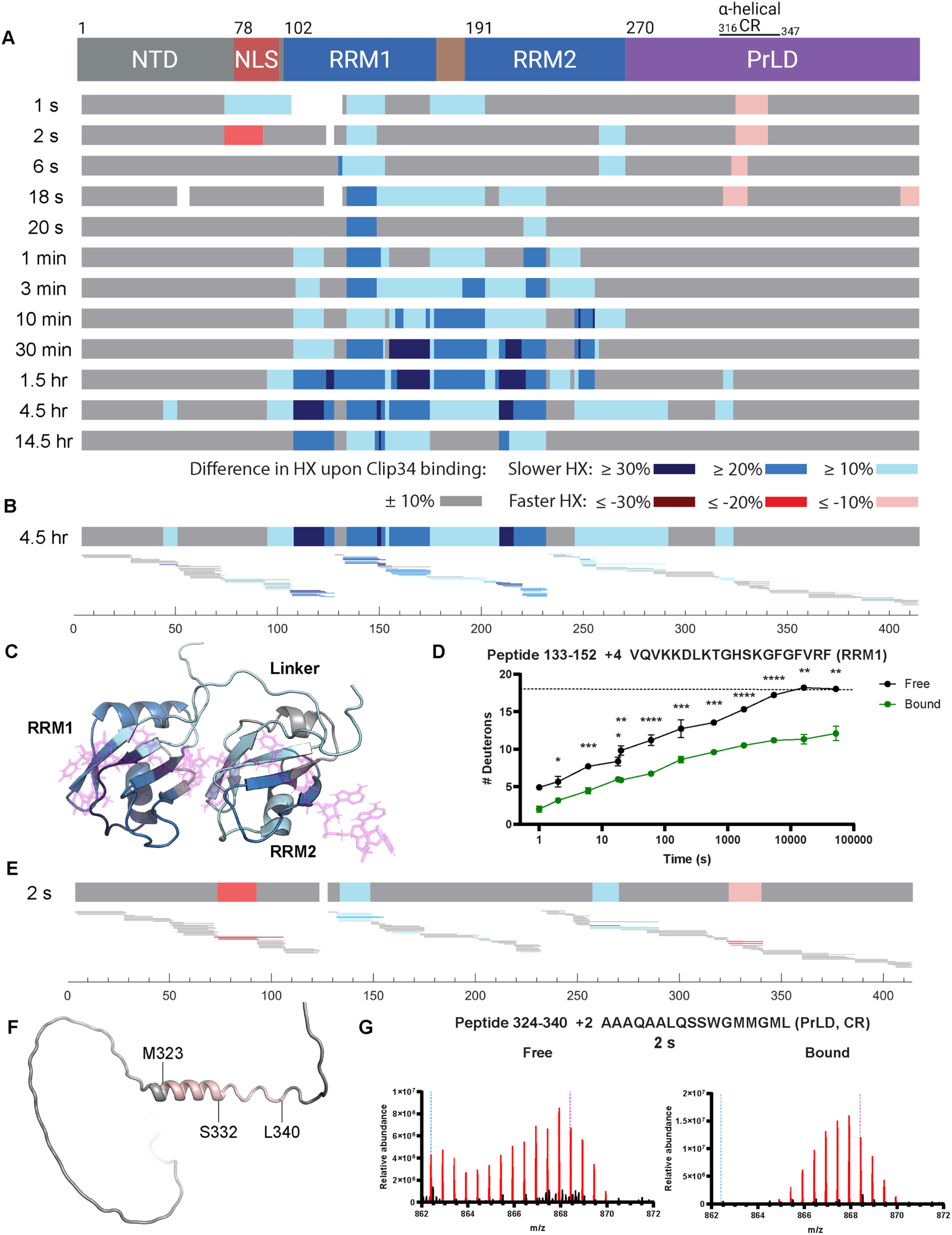
Hydrogen/deuterium-exchange reveals stabilization of the TDP-43 RRMs and destabilization of the CR within the PrLD upon Clip34 RNA binding. (A) *Top:* domain map of TDP-43, aligned with data underneath. *Bottom:* For each timepoint, residues are colored corresponding to the consensus percentage difference in exchange between the Clip34-bound (2:1::[Clip34]:[TDP-43]) and free states as indicated in the legend, as calculated by manual analysis of percentage exchange differences for all peptides including an amino acid. White spaces represent small coverage gaps. **(B)** The consensus percentage difference in exchange between Clip34-bound and free states at the 4.5 hr timepoint shown again as in (A). Aligned beneath it are peptides analyzed at the 4.5 hr timepoint, with percentage differences in exchange between Clip34-bound and free states for each peptide colored as shown in the legend for (A). Amino acid number is indicated on the axis below. **(C)** Image was generated using Pymol. Consensus percentage difference in exchange between Clip34-bound and free states at the 4.5 hr timepoint is shown on the structure of the RRMs bound to AUG12 RNA (PDB: 4BS2), colored according to the legend in (A). The RRMs are represented as a cartoon, whereas the RNA is represented as a stick. **(D)** HX for a representative RRM1 peptide. The dashed line represents the fully-deuterated condition. Data are mean ± SD (n=3-7 replicates run on mass spectrometry per timepoint; some error bars are too small to visualize; Welch’s t-test comparing bound and free at each timepoint; *p ≤ 0.05, **p ≤ 0.01, ***p ≤ 0.001, ****p ≤ 0.0001). **(E)** The consensus percentage difference in exchange between Clip34-bound and free states at the 2 s timepoint shown again as in (A). Aligned beneath it are peptides analyzed at the 2 s timepoint, displayed as in (B). **(F)** Image was generated using Pymol. Consensus percentage difference in exchange between Clip34-bound and free states at the 2 s timepoint shown on the cartoon representation of the AlphaFold structure of WT TDP-43 (Uniprot: Q13148), downloaded from the AlphaFold Protein Structure Database, colored according to the legend in (A). **(G)** Representative raw mass spectra at the 2 s timepoint for a representative peptide located in the CR of the PrLD. Signal corresponding to this peptide as determined by appropriate m/z values is colored red, whereas noise from overlapping peptide(s) is colored black. Dashed lines serve as visual guides; the blue dashed line indicates the monoisotopic peak, whereas the purple dashed line indicates the centroid value of the peptide in the fully deuterated sample.

We find that exchange is similar in the NTD in the Clip34-bound and free states, indicating that binding to Clip34 does not strongly impact NTD structure (Fig. 2A and fig. S6, A to C), which is consistent with our finding that the NTD is dispensable for Clip34 chaperone activity (Fig. 1F). By contrast, extensive decreases in exchange in the RRMs occurred in the presence of Clip34, particularly at later timepoints (Fig. 2, A to D, and fig. S6, D to I). This decreased exchange indicates that binding to Clip34 stabilizes the structure of the RRMs of TDP-43. Previous NMR studies have revealed the structure of the RRMs of TDP-43 in complex with a 12nt RNA (*35*). In our HXMS data, extensive stabilization was observed in both RRMs, including throughout the β-sheet binding surface determined in the NMR structure (Fig. 2C). In particular, sub-localizing exchange differences by comparing overlapping peptides reveals that Clip34 binding strongly stabilizes residues established to play key roles in binding, including F149 in RRM1, and F229 and F231 in RRM2 (Fig. 2B and fig. S5) (*35*). Additionally, at late timepoints Clip34 binding exerted one of its strongest stabilizing effects on the extreme C-terminal portion of RRM1 (Y155-D174) (Fig. 2A). Clip34-mediated stabilization of this C-terminal portion of RRM1 likely prevents any localized unfolding that is proposed to contribute to TDP-43 aggregation (*32*).

Exchange was rapid and unaffected by Clip34 for the majority of the PrLD, including both IDR1 and IDR2, as expected for an intrinsically disordered domain (Fig. 2A and fig. S6, J to L). There was, however, one important exception. Exchange in the CR in the PrLD increased in the presence of Clip34, particularly at the early timepoints (1-18 s) (Fig. 2A and E to G, and fig. S6, M and N, and fig. S7, A to C). This increased exchange indicates that Clip34 binding to TDP-43 destabilizes the CR in the PrLD. As displayed on the AlphaFold structure of TDP-43 (*65, 66*), the destabilization occurs from residue M323 within the predicted α-helix, to L340, at the end of the region predicted to have transient helical structure (Fig. 2F) (*37, 38*). Sub-localizing exchange differences reveals that Clip34 binding strongly destabilizes Q331 and S332 in the minor helical region (Fig. 2E and fig. S5). Thus, the destabilization of the PrLD upon Clip34 binding is specific to the CR and occurs throughout both the major and minor helical regions. Thus, these findings confirm the allosteric effect of Clip34 on the PrLD, which is expected to antagonize aggregation.

Delving deeper into this destabilization, we found that mass spectra for CR peptides exhibited bimodality at early timepoints (Fig. 2G and fig. S7, A to C). This bimodality suggests that there are two populations of TDP-43, with two different structures for the CR. Strikingly, the slow-exchanging, more stabilized, population is strongly represented in spectra at early timepoints in the free state, but poorly represented in the Clip34-bound state (Fig. 2G and fig. S7, A to C).

This finding indicates that Clip34 binding decreases the probability of a more stabilized CR structure. The CR is established to form a transient α-helical structure (*37, 38*). Hence, we suggest that in the absence of RNA, the TDP-43 CR forms a transient α-helix, whereas upon Clip34 binding, the TDP-43 CR is destabilized to disfavor α-helical structure. Given the important role of the α-helical structure of the CR in phase separation and aggregation (fig. S3B) (*32, 37, 38, 59*), this observation helps explain how Clip34 prevents TDP-43 aggregation. Specifically, Clip34 binding induces an aggregation-resistant form of TDP-43 with stabilized RRMs and a destabilized CR in the PrLD.

### Enhancing Clip34 activity against diverse disease-linked TDP-43 variants

Ideally, for maximal deployability, short RNA chaperones would mitigate aggregation of diverse disease-linked TDP-43 variants, including missense variants that cause disease, as well as forms of TDP-43 bearing pathological PTMs. Thus, we surveyed the chaperone capability of Clip34 against a suite of ALS/FTD-linked missense variants in RRM1 (P112H), the linker between RRM1 and RRM2 (K181E), or the PrLD (G295R, G298S, A321V, Q331K, M337V, and A382T), as well as PTM mimetic variants, including the pathological phosphorylation mimetics (S292E, S409/410E, and S292/409/410E), the pathological lysine acetylation mimetic (K145/192Q), and a physiological arginine methylation mimetic (R293F) (Fig. 3, A and B, and fig. S8, A and B) (*42, 46, 67–70*). These TDP-43 variants all aggregated to a similar extent over the time period studied (fig. S8, C and D). Strikingly, Clip34 prevented aggregation of all disease-linked TDP-43 variants, with half-maximal inhibitor concentration (IC_50_) values ranging from ∼0.12 µM-0.69 µM (Fig. 3, C and F, and fig. S8E). Notably, for a subset of TDP-43 variants Clip34 prevented aggregation more effectively than for WT TDP-43 (IC_50_∼0.5 µM), including TDP-43^P112H^ (IC_50_∼0.28 µM) located in RRM1 and TDP-43^K181E^ (IC_50_∼0.12 µM) located in the linker between RRMs (Fig. 3C). This result is intriguing as it has been suggested that these mutations may reduce RNA binding (*68–70*), indicating that Clip34 can overcome this deficit and still engage and chaperone effectively. Clip34 also exhibited a significantly lower IC_50_ against the phosphomimetic variants TDP-43^S409/410E^ (IC_50_∼0.29 µM) and TDP-43^S292/409/410E^ (IC_50_∼ 0.19 µM; Fig. 3C). Aberrant phosphoforms of TDP-43 accumulate in pathological inclusions (*45, 67*), but our findings indicate that Clip34 can antagonize aggregation of these species. Collectively our findings suggest that Clip34 has broad chaperone activity against different forms of TDP-43 connected to disease.

**Fig. 3.**
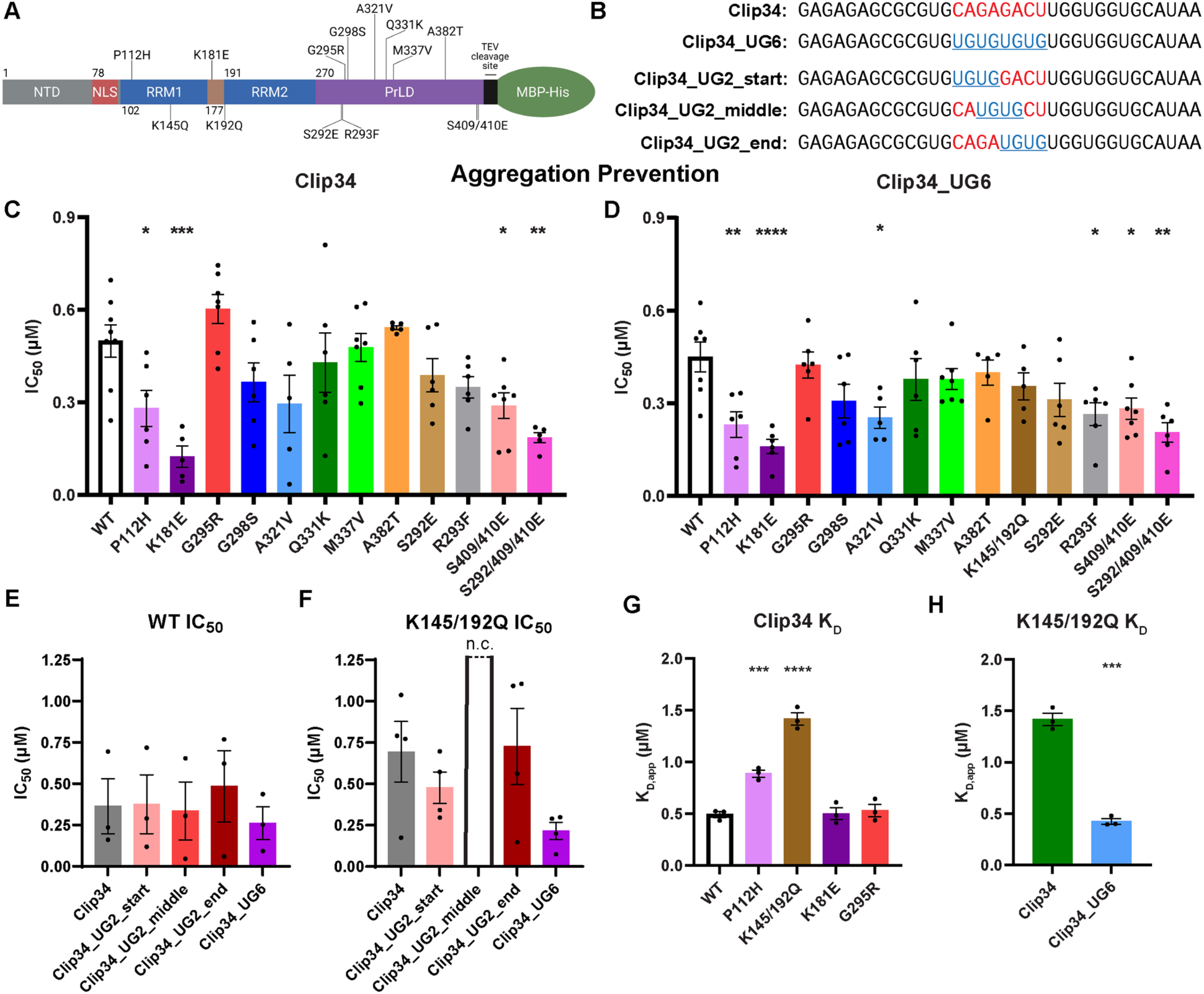
Clip34 and Clip34_UG6 effectively prevent aggregation of diverse disease-linked TDP-43 variants. **(A)** Domain map of TDP-43 to scale (excluding MBP-His solubility tags). Missense mutants (top) and post-translational modification mimetics (bottom) investigated in this study are indicated. **(B)** RNA sequences of Clip34 and its variants. Red text represents nucleotides in Clip34, whereas blue text represents nucleotides in Clip34_UG6. **(C, D)** TDP-43-MBP-His (5 µM) was incubated with TEV protease in the presence or absence of Clip34 (C) or Clip34_UG6 (D) for 16 hr, measuring turbidity every minute in a plate reader. Standardized turbidity data was normalized to the No RNA condition for that replicate; the AUC of this data was utilized to calculate an IC_50_ value (nonlinear regression: [inhibitor] vs normalized response with variable slope). Data are mean ± SEM (n=5-8; one-way ANOVA with Dunnett’s correction comparing to WT; *p ≤ 0.05, **p ≤ 0.01, ***p ≤ 0.001, ****p ≤ 0.0001). **(E, F)** IC_50_ values for Clip34 and Clip34 variants with WT TDP-43 (E) or TDP-43^K145/192Q^ (F). Data are mean ± SEM (n=3-4; n.c. indicates that an IC_50_ value could not be accurately calculated). **(G, H)** Apparent K_D_ values calculated from the bound signal from individual replicates of EMSAs performed with 5’ 6-FAM Clip34 and the indicated TDP-43 protein (G), or TDP-43^K145/192Q^ with 5’ 6-FAM Clip34 or Clip34_UG6 (H). Data shown for TDP-43^K145/192Q^ with Clip34 is the same in both figure parts, and WT TDP-43 with Clip34 as in Fig. 1, J and L. Data are mean ± SEM (n=3; one-way ANOVA with Dunnett’s correction comparing to WT (G), or unpaired t-test (H); ***p≤ 0.001, ****p ≤ 0.0001).

By contrast, Clip34 was less effective at antagonizing aggregation of the lysine acetylation mimetic TDP-43^K145/192Q^ (IC_50_∼0.69 µM) compared to WT TDP-43 (Fig. 3F and fig. S8, E to G). The K145Q:K192Q mutations reduce RNA binding (*46, 47, 61, 71*), which likely reduces Clip34 efficacy. This issue of reduced efficacy is problematic for Clip34, as TDP-43 acetylated at K145 accumulates in pathological inclusions in ALS (*46*).

To overcome this deficit of Clip34, we next defined an enhanced Clip34 variant with increased activity against TDP-43^K145/192Q^. This Clip34 variant, Clip34_UG6, harbors (UG)_4_ in place of CAGAGACU in the middle of the sequence (Fig. 3B). Clip34_UG6 binds to the isolated TDP-43 RRMs with higher affinity than Clip34 (*33*). Remarkably, Clip34_UG6 prevented aggregation of diverse disease-linked TDP-43 variants with IC_50_ values ranging from ∼0.16 µM-0.45 µM (Fig. 3D). Like Clip34, Clip34_UG6 was significantly more effective at inhibiting aggregation of the RRM1 variant, TDP-43^P112H^ (IC_50_∼0.23 µM), RRM1-RRM2 linker variant TDP-43^K181E^ (IC_50_∼0.16 µM), and the phosphomimetic variants TDP-43^S409/410E^ (IC_50_∼0.28 µM) and TDP-43^S292/409/410E^ (IC_50_∼0.21 µM) than WT TDP-43 (IC_50_∼0.45 µM; Fig. 3D and fig. S8F). Unlike Clip34, Clip34_UG6 was significantly more effective at inhibiting aggregation of the PrLD variant TDP-43^A321V^ (IC_50_∼0.25 µM) and the arginine methylation mimetic TDP-43^R293F^ (IC_50_∼0.26 µM) than WT TDP-43 (IC_50_∼0.45 µM; Fig. 3D). Importantly, Clip34_UG6 prevented aggregation of the double acetylation mimetic TDP-43^K145/192Q^ with a similar efficacy (IC_50_∼0.35 µM) as for WT TDP-43 (IC_50_∼0.45 µM; Fig. 3D). The enhanced activity of Clip34_UG6 against TDP-43^K145/192Q^ could not be recapitulated by introducing (UG)_2_ at different positions in the central portion of Clip34 (Fig. 3B). Although Clip34_UG2_start, Clip34_UG2_middle, and Clip34_UG2_end all displayed robust chaperone activity against WT TDP-43 (Fig. 3E), they were not as effective as Clip34_UG6 against TDP-43^K145/192Q^ (Fig. 3F and fig. S8G). Indeed, Clip34_UG2_middle was ineffective against TDP-43^K145/192Q^ (Fig. 3F and fig. S8G). Overall, these data suggest that Clip34_UG6 chaperone activity has broader applicability than Clip34 and is likely less affected by pathological K145/K192 acetylation.

To gain mechanistic insight into the differences in short RNA chaperone activity, we assessed whether differing IC_50_ values could be due to alterations in binding affinity. Thus, we determined the *K_D_* of select TDP-43 variants for Clip34. In some cases, *K_D_* tracked closely with IC_50_ (Fig. 3, G and H, and fig. S8, H and I). For example, TDP-43^G295R^ bound to Clip34 with a similar affinity as WT TDP-43, in accordance with the similar IC_50_ values of TDP-43^G295R^ and WT TDP-43 with Clip34 (Fig. 3, C and G, and fig. S8H). Moreover, TDP-43^K145/192Q^ exhibited impaired binding to Clip34 compared to Clip34_UG6, with binding affinity to Clip34 reduced by ∼2.9-fold compared to WT TDP-43 (Fig. 3, G and H, and fig. S8, H and I). This finding helps explain why Clip34 is less effective against TDP-43^K145/192Q^. In other cases, the *K_D_* does not precisely track with IC_50_.

For example, despite exhibiting lower IC_50_ values with Clip34 than WT TDP-43, TDP-43^P112H^ and TDP-43^K181E^ did not show increased binding affinity to Clip34 (Fig. 3, C and G, and fig. S8H). In fact, TDP-43^P112H^ displayed impaired binding to Clip34 compared to WT TDP-43 (Fig. 3G and fig. S8H). We suggest that while a certain threshold of binding affinity (*K_D_* < 1.4 µM) is critical for a short RNA to effectively chaperone TDP-43, other components of the interaction beyond simple binding affinity must contribute to chaperone activity.

### Defining additional potent short RNA chaperones against diverse disease-linked TDP-43 variants

To expand our arsenal of short RNA chaperones, we next defined additional RNAs that prevent aggregation of the broad spectrum of TDP-43 disease-relevant variants (Fig. 4A and fig. S9A). We first assessed the synthetic RNA, (UG)_17_, which is an extremely potent RNA chaperone (IC_50_∼0.2 µM) for WT TDP-43 (fig. S9B). (UG)_17_ also potently chaperoned all tested disease-relevant TDP-43 variants, including missense mutants in both the RRMs (TDP-43^P112H^ and TDP-43^K181E^) and the PrLD (TDP-43^G295R^ and TDP-43^Q331K^) (fig. S9B). Thus, simple repetitive UG sequences can effectively chaperone TDP-43.

**Fig. 4.**
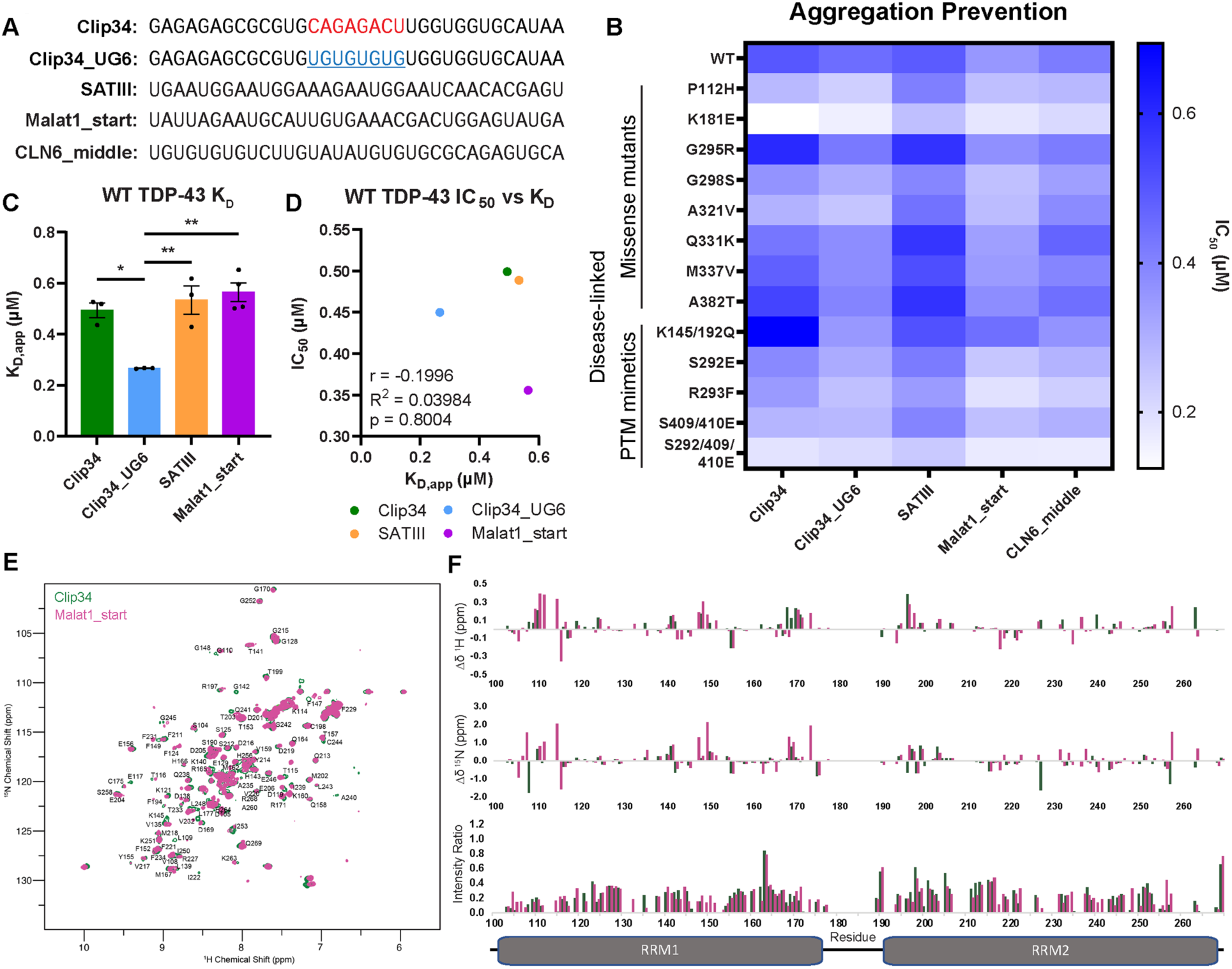
Malat1_start RNA displays enhanced chaperone activity against diverse disease-linked TDP-43 variants. **(A)** RNA sequences of tested RNAs. **(B)** Heatmap displaying mean values of the individual IC_50_ data shown in Fig. 3 (C, D, F) and in fig. S9 (C-E). **(C)** Apparent K_D_ values calculated from bound 5’ 6-FAM signal of the indicated RNAs, from individual replicates of EMSAs performed with WT TDP-43. Clip34 data is the same as shown in Fig. 3G. Data are mean ± SEM (n=3-4; one-way ANOVA with Tukey’s correction; *p ≤ 0.05; **p ≤ 0.01). **(D)** The K_D,app_ for WT TDP-43 with each indicated RNA, as shown in (C), is plotted against the IC_50_ for that RNA with WT TDP-43, as shown in (B) (Pearson correlation; not significant). **(E)** Overlay of ^1^H-^15^N heteronuclear single quantum coherence (HSQC) spectra of TDP-43 RRMs with Clip34 RNA (green, 2:1::[Clip34]:[TDP-43]) and Malat1_start RNA (magenta, 2:1::[Malat1_start]:[TDP-43]). **(F)** ^1^H and ^15^N chemical shift perturbations (Δδ^1^H (*top*) and Δδ^15^N (*middle*), respectively), and intensity ratios (*bottom*) of TDP-43 RRMs upon binding of Clip34 (green) and Malat1_start (magenta) RNA. Domain map of TDP-43 RRMs shown at the bottom, aligned to x-axes of graphs.

We next identified potent short RNA chaperones based on natural RNA sequences known to interact with TDP-43 (Fig. 4A). These include SATIII, derived from the pericentromeric satellite III repeat RNA; Malat1_start, from the *MALAT1* long non-coding RNA; and CLN6_middle, from the *CLN6* protein-coding transcript (*25, 59, 72–75*). We found that each of these short RNAs effectively chaperoned TDP-43 and disease-linked variants (Fig. 4B and fig. S9, C to H). The response of TDP-43 variants to these short RNAs closely resembled the pattern observed for Clip34 and Clip34_UG6 (Fig. 4B). However, compared to Clip34 and Clip34_UG6, there were fewer significant differences in the SATIII, Malat1_start, and CLN6_middle IC_50_ values for WT TDP-43 and the disease-associated TDP-43 variants (Fig. 3, C and D, and fig. S9, C to E). This finding suggests that the nature of how SATIII, Malat1_start, and CLN6_middle engage TDP-43 is more similar across TDP-43 variants than it is for Clip34 and Clip34_UG6. Overall, Malat1_start emerged as the most potent chaperone based on natural RNA sequences.

Malat1_start had the lowest IC_50_ values against WT TDP-43 (IC_50_∼0.36 µM) and disease-relevant variants (IC_50_ ranges from ∼0.17 µM-0.44 µM) (Fig. 4B and fig. S10, A and B). These findings support the development of Malat1_start as a short therapeutic RNA. Another feature that emerged across the tested short RNA chaperones and TDP-43 variants was a positive correlation between the IC_50_ value and the steepness of the hill slope for inhibition (fig. S10C). In general, RNAs inhibited TDP-43 aggregation in a cooperative manner with h ranging from −1.2 to −20 (fig. S10C). However, increased cooperativity correlated with decreased efficacy of the RNA at preventing TDP-43 aggregation.

In addition to Malat1_start and CLN6_middle, we tested two additional RNAs derived from both the *MALAT1* and *CLN6* RNAs: Malat1_middle, Malat1_end, CLN6_start, and CLN6_end (fig. S10A). These RNAs were effective at preventing WT TDP-43 aggregation, although less potently than Malat1_start or CLN6_middle, respectively, based on IC_50_ values (fig. S10, D to G). Each of these four RNAs also prevented aggregation of the TDP-43 RRM1 missense variant TDP-43^P112H^ and the RRM1-RRM2 linker variant TDP-43^K181E^ (fig. S10, D to G). Out of all tested natural RNAs, CLN6_end had the lowest potency against WT TDP-43 (IC_50_∼0.67 µM; Fig. 3C and fig. S9, C to E, and fig. S10, D to G). This trend was also captured by sedimentation analysis of the end timepoint of aggregation assays, where CLN6_end was less effective at maintaining WT TDP-43 in the soluble fraction (fig. S10, H and I).

To identify additional natural RNAs that effectively chaperone TDP-43, we also considered findings that G-quadruplex DNAs and RNAs can serve as general protein chaperones (*76, 77*). Indeed, a 28nt G-quadruplex-forming DNA sequence, *LTR-III*, can effectively chaperone denatured TagRFP675 protein (*76, 78*). As TDP-43 binds to HIV-1 LTR DNA (*36*), and the *LTR-III* RNA derived from this HIV-1 LTR sequence contains UG dinucleotides, we tested the ability of this RNA to prevent TDP-43 aggregation (fig. S10, A and J). We found that this short G-quadruplex-forming RNA effectively prevents aggregation of WT TDP-43 (fig. S10, J and K). Thus, short G-quadruplex RNAs emerge as potent chaperones for TDP-43.

### Effective RNA chaperones can engage the TDP-43 RRMs differently

Having assembled an arsenal of effective short RNA chaperones for TDP-43, we next set out to understand the basis of the differences in chaperone efficacy between RNAs. The binding affinity for WT TDP-43 or TDP-43^P112H^ for each RNA did not correlate with the IC_50_ values of these RNAs with the respective TDP-43 variant (Fig. 4, C and D, and fig. S11, A to C). This observation confirms that binding affinity is not the sole determinant of the ability of a short RNA to prevent TDP-43 aggregation. Another factor that may influence chaperone activity is exactly how each RNA engages the TDP-43 RRMs. To understand whether different short RNAs engage the TDP-43 RRMs in the same way, we conducted NMR experiments on the isolated RRMs of TDP-43 in solution with two-fold excess Clip34, Clip34_UG6, and Malat1_start to saturate binding. We observed overall similarities in resonance shifts and broadening, which report on molecular structure and conformational exchange as these three RNAs engage the RRMs, particularly in RRM1 (residues 138-142 and 160-172), indicating that these regions are important for binding independent of the details of the RNA sequence (Fig. 4, E and F, and fig. S11, D and E). However, there were focused regions of discernable differences between the three different RNAs, indicative of changes in interaction and conformational exchange and motions dependent upon the RNA sequence (Fig. 4, E and F, and fig. S11, D and E). When in complex with Clip34 compared to with Malat1_start, the TDP-43 RRMs showed more broadening and unique shifts in the region of residues 145-151 in the third beta-strand of RRM1 (Fig. 4, E and F). This region harbors K145 that can be acetylated to disrupt RNA binding (*46*), and F147 and F149 that stack to interact with a U or G base, respectively, and are essential for RNA binding in RRM1 (*35*). Additionally, broadening and unique shifts were also observed in the region of residues 104-106 on the adjacent first beta-strand of RRM1 (Fig. 4, E and F). Therefore, unique interactions or motions in these regions are present for Clip34 compared to Malat1_start.

Additionally, Clip34_UG6 also showed distinct perturbations compared to Clip34 in the vicinity of K145, as well as large chemical shift perturbations at three positions between 130 and 140, distinguishing these two similar sequences (fig. S11, D and E). These differences may help explain why Clip34 is less effective in chaperoning the pathological acetylation mimic, TDP-43^K145/192Q^ (*46*), in comparison to Clip34_UG6 and Malat1_start, which are more effective (Fig. 4B). We conclude that while RNAs exhibit partial similarity in their interactions with the TDP-43 RRMs, there are also noticeable differences in specific regions, which likely translate into differences in chaperone activity.

### Short RNAs reduce cytoplasmic TDP-43 aggregation in an optogenetic human cell model

We next assessed whether short RNAs combat TDP-43 aggregation in human cells. We utilized an optogenetic model of TDP-43 proteinopathy, in which TDP-43 is fused to Cry2olig, a domain derived from the cryptochrome 2 protein that undergoes homo-oligomerization upon exposure to blue light (*79*). This homo-oligomerization results in the formation of cytoplasmic TDP-43 puncta that display typical hallmarks of disease, including colocalization with p62 and pTDP-43(pS409/410) signal (*29*). Using this optogenetic model in human (HEK293) cells, we tested the effect of a subset of short RNAs with a range of activities *in vitro*: Malat1_start (IC_50_∼0.36 µM), (UG)_17_ (IC_50_∼0.2 µM), and CLN6_middle (IC_50_∼0.42 µM). Compared to cells treated with a control (CTR) RNA not expected to bind TDP-43, Malat1_start and (UG)_17_ significantly reduced the average area of cytoplasmic TDP-43 inclusions per cell, whereas CLN6_middle did not (Fig. 5, A and B, and fig. S12, A to C). Malat1_start and (UG)_17_ were the two most effective RNAs at reducing cytoplasmic TDP-43 puncta area in human cells and preventing TDP-43 aggregation *in vitro*. Thus, our *in vitro* aggregation assays provide a powerful platform for identifying RNAs with chaperone activity against TDP-43 in human cells.

**Fig. 5.**
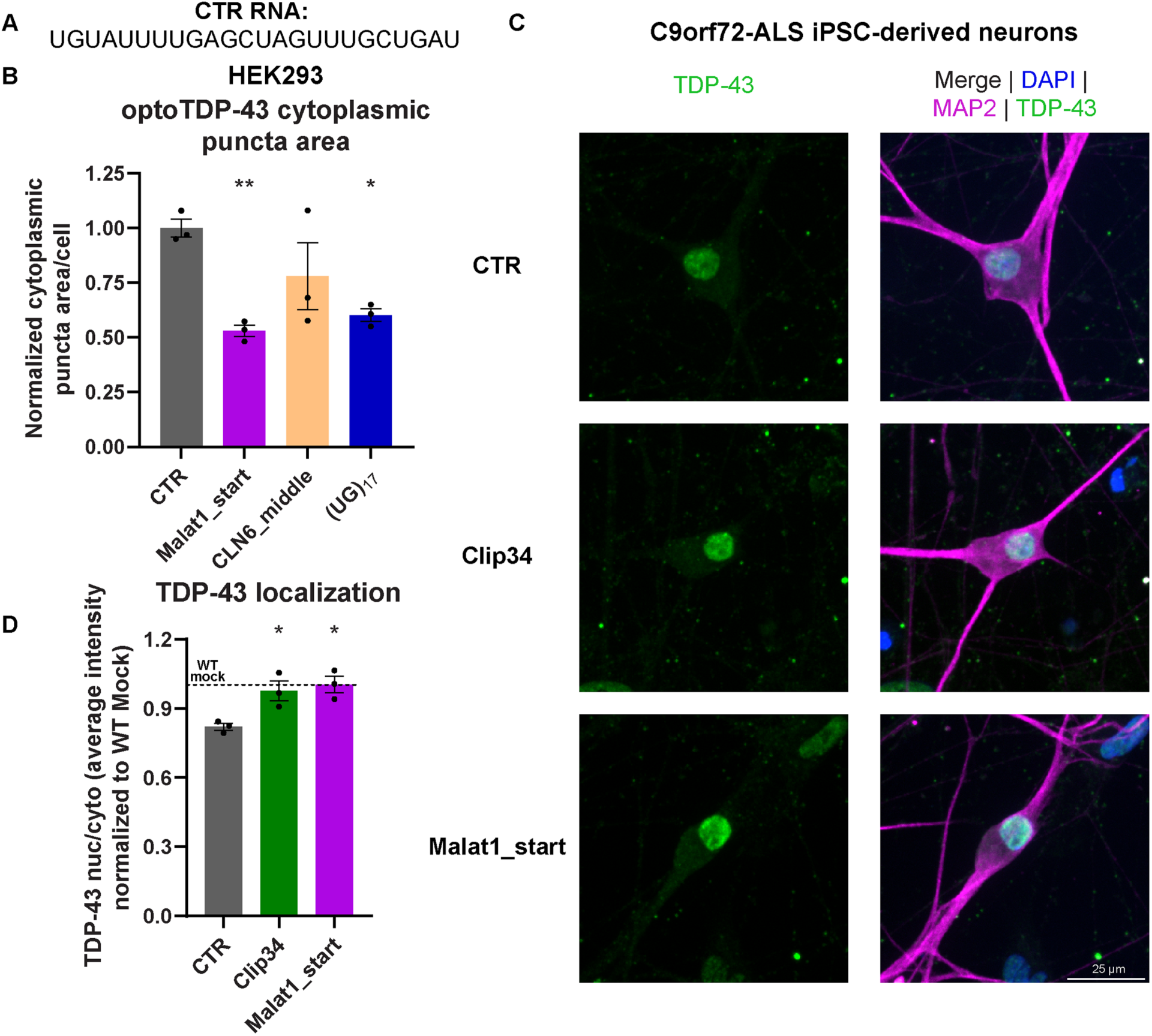
Malat1_start RNA mitigates aberrant TDP-43 phenotypes in optogenetic human cell models and patient-derived neurons. **(A)** Sequence of the control (CTR) RNA. **(B)** OptoTDP-43 stable HEK293 cells were treated with the indicated RNA, followed by blue light exposure to induce Cry2olig oligomerization, and imaged after fixation. The average area of cytoplasmic puncta per cell, normalized to the average of the CTR-treated condition. Data are mean ± SEM (n=3 biological replicates; one-way ANOVA with Dunnett’s correction comparing to CTR; *p ≤ 0.05; **p≤ 0.01). **(C)** Representative images of C9orf72-ALS patient iPSC-derived neurons treated with the indicated RNAs, stained with DAPI and for TDP-43 and MAP2. Scale bar indicates 25 μm. **(D)** The average ratio of TDP-43 nuclear to cytoplasmic signal, normalized to healthy control iPSC-derived neurons without RNA treatment. Data are mean ± SEM (n=3 biological replicates, represented as the average of n=2 technical replicates each; one-way ANOVA with Dunnett’s correction comparing to CTR; *p ≤ 0.05).

### Clip34 and Malat1_start RNAs do not cause TDP-43 loss of function

A possible concern with employing short RNAs in this way is that they might remain too stably bound to TDP-43 and interfere with essential RNA processing reactions. However, when RNA-binding proteins are engaged by nuclear-import receptors in the cytoplasm for transport to the nucleus, bound RNA is ejected, such that the short RNA would be recycled for further rounds of chaperone activity (*80, 81*). Clip34 is not toxic to human cells and does not affect nuclear localization of endogenous TDP-43 (*28*). Moreover, Clip34 does not inhibit TDP-43 function in pre-mRNA splicing reactions (*28*). To assess whether Malat1_start, Clip34, and (UG)_17_ might interfere with TDP-43 function, we employed the CUTS (CFTR UNC13A TDP-43 Loss-of-Function) biosensor, a cryptic exon RNA biosensor enabling real-time detection of TDP-43 loss of splicing function (*82*). CUTS can detect even an ∼10% decrease in TDP-43 functionality (*82*). Clip34 and Malat1_start did not interfere with TDP-43 function (fig. S12D). By contrast, (UG)_17_ interfered with TDP-43 function, and this effect was larger than a siRNA positive control that reduces TDP-43 expression by ∼2% (fig. S12D) (*82*). Thus, (UG)_17_ displays undesirable properties and hence did not advance to studies with patient-derived neurons or mice.

## Short RNAs restore physiological TDP-43 localization in ALS patient-derived neurons

To further assess the translational potential of short RNAs, we utilized induced pluripotent stem cell (iPSC)-derived neurons from a healthy control or individuals with ALS genetically caused by a hexanucleotide repeat expansion in the *C9orf72* gene (C9-ALS). Compared to healthy control iPSC-derived neurons without RNA treatment (mock) or treated with the CTR RNA, C9-ALS iPSC-derived neurons exhibited TDP-43 pathology in the form of a decreased TDP-43 nuclear/cytoplasmic ratio (fig. S12E). Treatment of C9-ALS iPSC-derived neurons with Clip34 or Malat1_start, but not the CTR RNA, increased the TDP-43 nuclear/cytoplasmic ratio to the level of healthy control iPSC-derived neurons with mock treatment (Fig. 5, C and D). Thus, Clip34 and Malat1_start short RNAs can mitigate aberrant cytoplasmic mislocalization of TDP-43 and enable the return of TDP-43 to the nucleus in C9-ALS patient-derived neurons.

### Malat1_start mitigates neurodegeneration and reverses TDP-43 aggregation in mice

After establishing the ability of Malat1_start to mitigate aberrant TDP-43 phenotypes in human cells and ALS patient-derived neurons, we next tested this short RNA *in vivo*. We utilized an acute spinal expression paradigm (Fig. 6A) (*83*). We generated Adeno-associated virus (AAV) 9 containing the CMV-promoter driven TDP-43 NLS1 YFP plasmid (*84*), which results in cytoplasmic YFP-tagged TDP-43 due to a mutated nuclear localization signal (TDP-43^ΔNLS^).

**Fig. 6.**
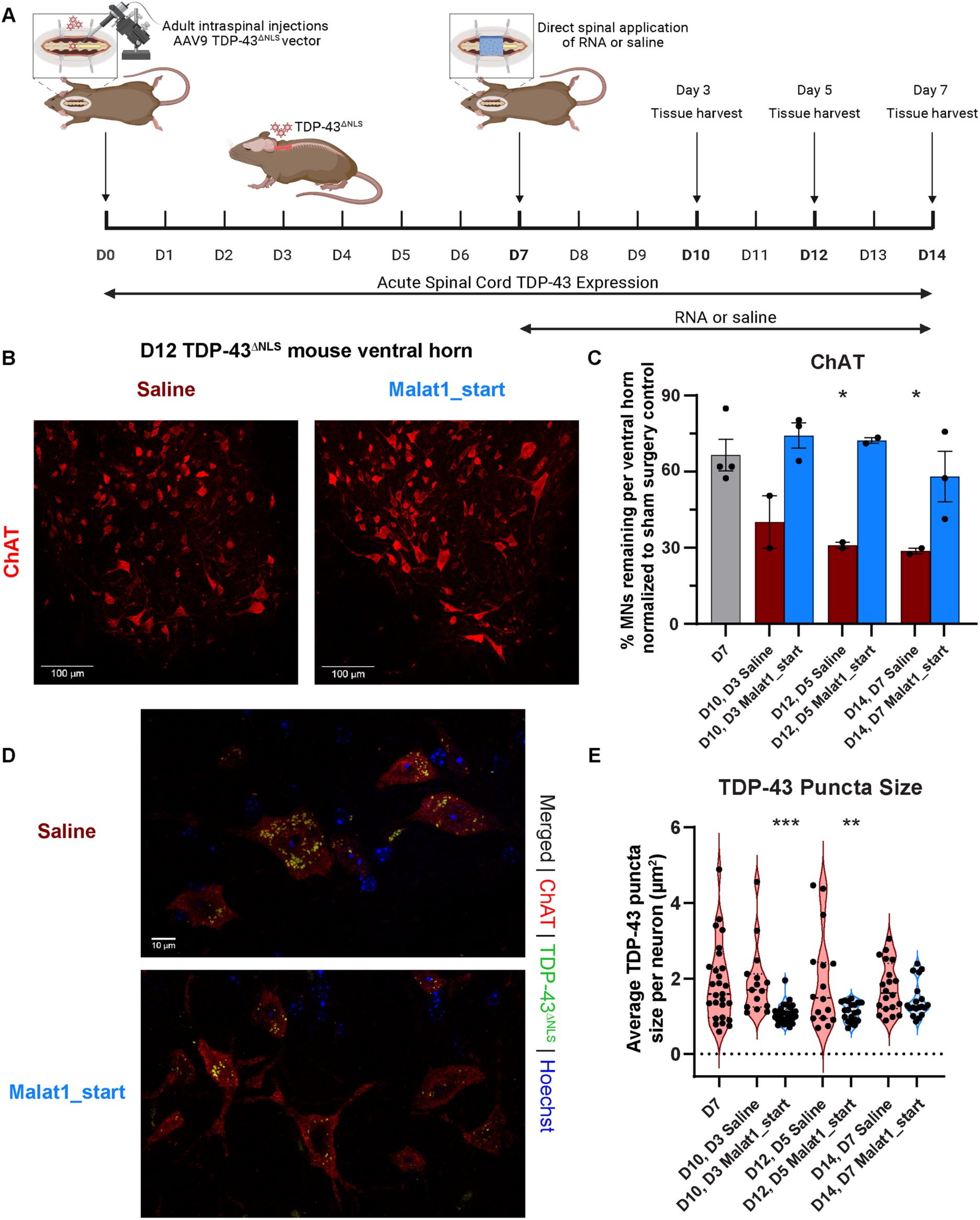
Malat1_start RNA mitigates neurodegeneration and TDP-43 aggregation in a mouse model of TDP-43 proteinopathy. **(A)** Schematic of experimental paradigm. On Day 0 (D0), animals undergo laminectomy and bilateral AAV9 viral injection across the C4-C6 region, to express TDP-43^ΔNLS^ throughout the ventral horns of cervical spine. On D7, animals undergo a second surgery to receive RNA or saline. Histology was assessed at days 3 (D10), 5 (D12), and 7 (D14) following treatment. **(B)** Representative immunohistochemistry images of ventral horns at D12, stained for choline acetyltransferase (ChAT). 20x magnification z-stack confocal images; scale bar indicates 100 μm. **(C)** ChAT^+^ motor neurons were manually counted within the ventral horn of spinal cord sections. Data are mean ± SEM (n=2-4 animals per condition; n=4 ventral horns per animal; shown: one-way ANOVA with Dunnett’s correction comparing to D7 TDP-43; *p ≤ 0.05; not shown: two-way ANOVA on D10, D12, and D14 data: treatment (Malat1_start vs saline): ***p ≤ 0.001). **(D)** Representative 60x magnification immunofluorescent staining images from 5-day (12-day expressing) saline-treated (top) and RNA-treated (bottom) animals. Scale bar indicates 10 μm. **(E)** TDP-43 positive puncta were assessed in ChAT^+^ motor neurons at 60x magnification in the ventral horn for each animal. Average puncta size was calculated per field of view. Data shown are mean ± SEM (n=2-3 animals per condition; 5-10 fields of view per animal; shown: one-way ANOVA with Dunnett’s correction comparing to D7 TDP-43: **p ≤ 0.01, ***p ≤ 0.001; not shown: two-way ANOVA on D10, D12, and D14 data: treatment (Malat1_start vs saline): ****p ≤ 0.0001).

This virus was bilaterally injected across six sites in the cervical spinal cord of p180 mice (Fig. 6A). At day 7 (D7), we observed highly efficient viral delivery to the ventral horn, and robust expression of TDP-43^ΔNLS^ (fig. S13A), which formed cytoplasmic puncta (fig. S13B). At D7, animals were then treated with either saline or Malat1_start.

We then determined the effect of RNA treatment on motor neuron loss and TDP-43 aggregation. We verified RNA penetration to the spinal cord ventral horn (fig. S13B). We counted motor neurons from these mice with two neuronal markers: choline acetyltransferase (ChAT), which denotes cholinergic motor neurons, and NeuN, which stains neuronal nuclei. Saline-treated TDP-43^ΔNLS^ animals displayed a progressive temporal loss of ChAT^+^ motor neurons, with the average percent of remaining motor neurons, compared to sham surgery controls, of 66.5% at D7 falling to 40.1% at D10, 31.0% at D12, and 28.7% at D14 (Fig. 6, B and C, and fig. S13C). By contrast, Malat1_start-treated TDP-43^ΔNLS^ animals maintained their ChAT^+^ motor neurons over this time course, with averages of 73.9% at D10, 72.2% at D12, and 58.1% at D14 (Fig. 6, B and C, and fig. S13C). Similarly, assessment of ventral horn NeuN^+^ neurons revealed that in saline-treated TDP-43^ΔNLS^ animals, the average of 70.4% of neurons at D7, compared to sham surgery controls, progressively decreased to 62.3% at D10, 48.2% at D12, and 41.8% at D14 (fig. S13D). NeuN^+^ neurons were also maintained in the Malat1_start-treated TDP-43^ΔNLS^ animals, with average neuronal values comparable to the D7 baseline across the weeklong time course (68.6% at D10, 66.8% at D12, and 63.0% at D14) (fig. S13D). Thus, Malat1_start mitigates TDP-43-driven neurodegeneration *in vivo*.

We next used three-dimensional image analysis to determine the average TDP-43 puncta size per motor neuron. The average TDP-43 puncta size in saline-treated animals was unchanged from the D7 baseline at all timepoints, whereas Malat1_start treatment reduced puncta size at D10 and D12 compared to the D7 baseline (Fig. 6, D and E). Thus, Malat1_start partially reversed TDP-43 aggregation, indicating that the short RNA solubilizes preformed aggregates *in vivo*. We also determined the change in the average number of TDP-43 puncta per motor neuron in each animal. The average total TDP-43 puncta per cell increased progressively in saline-treated animals, starting at 30.2 at D7 and increasing to 34.8 at D10, 38.5 at D12, and 40.6 at D14 (Fig. 6D and fig. S13E). However, at D10 and D12 in Malat1_start-treated animals, the number of TDP-43 puncta was still comparable to the D7 baseline (averages of 33.0 at D10, 34.3 at D12) (Fig. 6D and fig. S13E). By D14, puncta did begin to increase in Malat1_start-treated animals compared to the D7 baseline (average of 35.5 at D14) (fig. S13E). Nevertheless, these findings suggest that a single dose of Malat1_start RNA reduces TDP-43 aggregation and motor neuron degeneration *in vivo*.

## Discussion

In this study, we have defined allosteric crosstalk between the TDP-43 RRMs, PrLD, and RNA, which dictates the propensity of TDP-43 to access soluble or aggregation-prone states.

Specifically, the RRMs and PrLD form a sensitive interdependent module within TDP-43. When a short 34nt RNA binds to the TDP-43 RRMs, it acts as a molecular stabilizer, allosterically inducing the PrLD to populate disordered states with reduced propensity to self-assemble. For example, Clip34 engages and stabilizes the RRMs, and destabilizes a transient α-helix in the CR of the PrLD. This destabilization promotes solubility and disfavors self-assembly. Indeed, Clip34 binding elicits strong local destabilization of structure at residues Q331 and S332 in the CR. This allosteric effect suggests that short RNA binding does more than simply anchor TDP-43 to the nucleic acid. Rather, the short RNA alters TDP-43 conformation such that aggregation-prone states are depopulated.

Deletion of RRM2 more drastically impaired the ability of Clip34 to chaperone TDP-43 than deletion of RRM1. As RRM2 is proximal to the PrLD, we suggest that RRM2 transmits the effect of RNA binding to the PrLD. In the apo state, the CR region of the PrLD morphs from a disordered state to an α-helical form, which promotes intermolecular contacts between PrLDs that drive phase separation (*37, 38*). Ultimately, an aberrant phase transition occurs when the α-helical CR transitions to the intermolecular beta-sheet structure of TDP-43 fibrils (*16, 17*). By allosterically promoting disorder in this region, RNA effectively chaperones TDP-43 and prevents pathological aggregation. Intriguingly, although the destabilizing effect of Clip34 is specific to the CR when measured for soluble full-length TDP-43, Clip34 can also prevent aggregation of TDP-43 deletion constructs lacking the CR. Thus, Clip34 binding to the RRMs likely imposes additional allosteric effects on the PrLD beyond the CR that maintain intrinsic disorder, and thereby prevent formation of intermolecular contacts and structures that drive aggregation.

Conversely, the PrLD negatively regulates the affinity of the RRMs for RNA. The increase in RNA binding when the PrLD is deleted suggests that the PrLD likely moderates RNA-binding capacity of TDP-43, thereby maintaining a balanced engagement with RNA under physiological conditions. Within the PrLD, the CR plays an important role in this negative regulation, as deletion of the CR increases the affinity of TDP-43 for Clip34 RNA by ∼2.1-fold. Intriguingly, we found that TDP-43 remodels Clip34 by separating its 5’ and 3’ ends, an activity that was also negatively regulated by the CR and PrLD. It will be important to determine whether this RNA-remodeling activity is important for TDP-43 function. Consistent with our previous finding that the PrLD modulates TDP-43 binding and function at a subset of endogenous RNA regions (*59*), our findings suggest that the PrLD acts as a regulatory hub for tuning RNA interactions.

Importantly, deletion of the CR confers impaired neuronal function and behavioral abnormalities in mice, reinforcing the critical role of the CR in regulating TDP-43 function (*85*). Overall, our findings suggest that interplay between the RRMs, PrLD, and RNA maintains a precise balance between TDP-43 solubility and self-assembly propensity. TDP-43 likely responds dynamically to environmental cues, such as RNA availability, to maintain solubility and avoid aggregation.

We suggest that RNA-depleted environments, such as the cytoplasm of aging neurons or the interior of aging stress granules, place TDP-43 at risk for pathological aggregation (*13, 32, 86*). Indeed, pathological TDP-43 inclusions are typically depleted of RNA (*29*).

Importantly, our initial lead RNA, Clip34, is effective at preventing the pathological aggregation of diverse ALS/FTD-linked forms of TDP-43, including the WT protein, and several missense variants in RRM1 (P112H), the linker between RRM1 and RRM2 (K181E), and in the PrLD (G295R, G298S, A321V, Q331K, M337V, and A382T). Thus, our approach is likely to be broadly applicable to the majority of sporadic ALS cases in addition to rare familial forms caused by TDP-43 mutations. The ability to counter aggregation of TDP-43^P112H^ and TDP-43^K181E^ was surprising, as these TDP-43 variants have been reported to have reduced RNA binding (*68–70*), indicating that Clip34 can overcome this deficit and still chaperone effectively. Clip34 could also effectively chaperone pathological phosphomimetic variants of TDP-43 (S409/410E and S292/S409/410E) and the physiological PrLD arginine methylation mimetic (R293F). However, Clip34 was less effective against pathological RRM lysine acetylation mimetic (K145/192Q). This reduction in activity is problematic, as TDP-43 acetylated at K145 accumulates in pathological inclusions in ALS (*46*). Hence, we sought enhanced, short RNA chaperones that were also effective against TDP-43^K145/K192Q^.

By mining the sequence space of natural and synthetic short RNAs that engage TDP-43, we have uncovered many short RNAs that can effectively chaperone TDP-43. Of these, the two most potent were an engineered variant of Clip34, Clip34_UG6, and Malat1_start derived from the *MALAT1* long non-coding RNA. Clip34_UG6 and Malat1_start effectively chaperoned diverse disease-linked TDP-43 variants, including TDP-43^K145/K192Q^. A short synthetic RNA, (UG)_17_, was also effective. Importantly, Malat1_start and (UG)_17_ could also mitigate TDP-43 aggregation in an optogenetic model of TDP-43 proteinopathy in human cells. Thus, their activity extends beyond the pure protein level. However, a concern with using short RNAs in this way is that the RNA may remain too stably bound to TDP-43 and interfere with TDP-43 function. Indeed, using a sensitive CUTS reporter of TDP-43 functionality (*82*), we found that (UG)_17_ interfered with TDP-43 function in human cells. Thus, this synthetic RNA is unsuitable for further development. Importantly, neither Malat1_start nor Clip34 interfered with TDP-43 function (*87*).

Remarkably, treatment of C9-ALS iPSC-derived neurons with Clip34 or Malat1_start short RNAs restored nuclear TDP-43 localization to levels seen in control neurons. Thus, Clip34 and Malat1_start short RNAs can correct TDP-43 localization in patient-derived neurons. These findings also suggest that short RNA chaperones have applications in C9-ALS/FTD cases, which also present with TDP-43 proteinopathy.

Finally, we assessed the ability of Malat1_start to mitigate aberrant TDP-43 phenotypes in an acute spinal expression paradigm in mice in which AAV9 delivers TDP-43^ΔNLS^. TDP-43^ΔNLS^ aggregates in the cytoplasm and elicits motor neuron degeneration within 2 weeks. A challenge facing short RNA therapeutics is drug delivery, but we show here that 2’OMe-modified Malat1_start bearing five phosphorothioate linkages at each end can be readily delivered into the cytoplasm of neurons by direct spinal cord application. Thus, we delivered Malat1_start after 7 days of TDP-43^ΔNLS^ expression, at which time TDP-43 aggregates had already accumulated in the cytoplasm of motor neurons. Remarkably, Malat1_start reduced TDP-43 aggregation and prevented motor neuron degeneration in this model. Thus, a single dose of Malat1_start RNA reduces TDP-43 aggregation and motor neuron degeneration *in vivo*.

Our studies reveal mechanisms of short RNA chaperones and pave the way for their development as therapeutics for fatal TDP-43 proteinopathies. Our findings present a highly actionable therapeutic opportunity for diverse TDP-43 proteinopathies, including ALS/FTD, AD, LATE, and CTE. By transiently engaging TDP-43 with a short RNA therapeutic, we can shift the equilibrium back to soluble forms of TDP-43, which can be transported to the nucleus to mitigate TDP-43 loss of function. Short RNAs would preferentially engage TDP-43 in the cytoplasm where there is less competition from endogenous RNA, which is more concentrated in the nucleus (*51*). Moreover, as nuclear-import receptors bind solubilized TDP-43 in the cytoplasm, they will eject the bound RNA, such that the short RNA can be recycled for further rounds of TDP-43 solubilization in the cytoplasm. In this way, an apo form of TDP-43 is chaperoned back into the nucleus safely by nuclear-import receptors, where it can functionally engage RNA targets (*88*). Importantly, our short RNAs are similar in size and chemistry to FDA-approved ASOs, which can be readily delivered to the spinal cord and brain parenchyma of patients to exert therapeutic effects in ALS caused by SOD1 mutations and spinal muscular atrophy (*57*).

## Acknowledgements

We thank Linamarie Miller, Edward Barbieri, and JiaBei Lin for critiques, and Kristen Lynch for access to the Typhoon scanner. We thank Nikaela Bryan for preliminary digestion and HXMS conditions and HX rate determination of TDP-43. We thank Emily Smith for assistance with HMXS data visualization and MS data deposition. Some figure subpanels were created with BioRender.com.

## Author contributions

Conceptualization: KEC, CJD, & JS

Data curation: KEC, JCM, HD, MN, LX, AS, ML, CJD, BKJ, & JS

Formal analysis: KEC, JCM, HD, MD, AS, LM, ML, BKJ, & JS

Funding acquisition: KEC, HD, ML, BP, HMO, PP, NLF, BEB, CJD, BKJ, & JS

Investigation: KEC, JCM, HD, MN, LX, AS, MD, LM, ML, JDR, BP, BLL, HMO, & BKJ

Methodology: KEC, JCM, HD, MN, LX, LM, ML, JDR, BP, MH, JU, PP, YP, NLF, BEB, CJD, BKJ, & JS

Project administration: KEC, PP, NLF, BEB, CJD, BKJ, & JS

Resources: KEC, JCM, HD, LX, AS, LM, ML, JDR, MH, JU, PP, NLF, BEB, CJD, BKJ, & JS

Supervision: KEC, LM, PP, NLF, BEB, CJD, BKJ, & JS

Validation: KEC, JCM, HD, MN, LX, AS, LM, ML, JDR, BP, BLL, HMO, BKJ, & JS

Visualization: KEC, JCM, HD, LX, AS, LM, NLF, BEB, CJD, BKJ, & JS

Writing – original draft: KEC, HD, BKJ, & JS

Writing – review & editing: KEC, JCM, HD, MN, LX, AS, MD, LM, ML, JDR, BP, BLL, HMO, MH, JU, PP, YP, NLF, BEB, CJD, BKJ, & JS

## Data and materials availability

Plasmids generated in this study will be made readily available to the scientific community. All requests will be honored in a timely manner. Material transfers will be made with no more restrictive terms than in the Simple Letter Agreement or the Uniform Biological Materials Transfer Agreement and without reach through requirements. All data reported in this paper will be shared by the lead contact upon request. This paper does not report original code. Any additional information required to reanalyze the data reported in this paper is available from the lead contact upon request.

## Supplementary Materials for

### Materials and Methods

#### Animals

Experimental procedures were approved by the Institutional Animal Care and Use Committee (IACUC) at Thomas Jefferson University and were conducted in compliance with the Guide for the Care and Use of Laboratory Animals from the National Institutes of Health. Female non-transgenic C57BL/6J mice aged 180 days were acquired from Jackson Laboratories (https://www.jax.org/strain/000664) and housed in an animal facility with controlled humidity, temperature and light cycles, with access to *ad libitum* water and standard chow. As analgesics are delivered on the basis of animal weight, and to better control variance in animal starting weight, female mice were exclusively chosen for this present study.

This study represents a total of 26 animals having undergone the spinal surgeries described below. Initial characterization of viral expression used n=3 animals in sham surgery, and TDP-43^ΔNLS^ groups expressing virus for 7 days. In the main cohort of animals, there were n=2 sham non-injected animals and 18 animals which received injections to express TDP-43^ΔNLS^. After one week, all 20 animals underwent a second surgery. Sham animals again received no treatment, whereas TDP-43^ΔNLS^ animals were subdivided into saline and RNA treatment groups (n=9 for each). Of these two groups, n=2-3 were used for each of the 3-day, 5-day, and 7-day endpoints for immunostaining analysis. The n=2 sham animals were also collected at the 7-day endpoint.

#### Cell lines

Induced pluripotent stem cell (iPSC) lines CS15iCTR-5, CS29iALS-n1, and CS52iALS-n6A were obtained from the Cedars-Sinai RMI iPSC Core, and are male. Line JH034 was obtained from Johns Hopkins Hospital, and is female. The estimated G4C2 repeat expansion sizes are >2.5 kb for line JH034, and 6-8 kb for lines CS29iALS-n1 and CS52iALS-n6A.

iPSCs were differentiated following previously described protocols (*89, 90*). iPSCs were cultured in Matrigel (Corning) and mTeSR+ (StemCell Technologies) and kept in a humidified chamber with regulated levels of CO_2_ (5%) and temperature (37°C). For differentiation, 1×10^6 iPSCs were plated in 6-well plates. Once cells reached ∼90% confluency, media was changed from mTeSR+ to N2B27 media (50% DMEM:F12, 50% Neurobasal, plus NEAA, Glutamax, N2 and B27; all from Gibco) plus 10 μM SB431542 (StemCell Technologies), 100 nM LDN-193189 (Sigma-Aldrich), 1 μM RA (Sigma-Aldrich) and 1 μM Smoothened-Agonist (SAG, Cayman Chemical). Media was changed daily for a total of 6 days. Cells were then switched to N2B27 including 1 μM RA, 1 μM SAG, 4 μM SU5402 (Cayman Chemical) and 5 μM DAPT (Cayman Chemical), and media was changed daily until day 13. Neurons were dissociated on day 14 using TrypLE and DNAse I, and plated in Matrigel-coated 24-well plates with glass coverslips for confocal imaging studies. Cells were fed every other day and maintained for 13 days after plating in Neurobasal media + NEAA, Glutamax, N2, B27, plus 10 ng/mL BDNF, GDNF, CNTF (all from PeproTech) and 0.2 μg/mL Ascorbic acid (Sigma-Aldrich).

#### Microbe strains

*Escherichia coli* BL21 (DE3)-RIL cells (Agilent 230245) and BL21 Star (DE3) Chemically competent cells (Thermo Fisher C601003) were utilized for protein purification, with growth conditions as described in the purification sections. *Escherichia coli* XL10-Gold Ultracompetent cells (Agilent 200314) were utilized for cloning and plasmid propagation, and were grown at 37°C with the appropriate antibiotic.

#### Cloning

pJ4M was from Addgene (plasmid #104480; http://n2t.net/addgene:104480; RRID:Addgene_104480) (*91*). All other TDP-43 constructs purified were generated using the pJ4M plasmid. Partial PrLD deletion plasmids were generated previously (*59*). TDP-43^5FL^ was generated previously (*29*). TDP-43^S292E^, TDP-43^R293F^, TDP-43^S409/410E^, and TDP-43^S292/409/410E^ plasmids were generated previously (*67*). All other TDP-43 disease-relevant variants and domain deletion plasmids, as well as the MBP-His plasmid, were generated via QuikChange Site-Directed Mutagenesis (Agilent 210518). MBP-FUS was generated previously (*92*).

#### Purification of TEV protease

TEV protease was purified as previously described (*93*). His-TEV plasmid was transformed into BL21 (DE3)-RIL *E. coli* and grown on an LB-ampicillin plate at 37°C for 16 hr. The cells were then transferred to a starter culture of LB containing 100 µg/mL ampicillin and 34 µg/mL chloramphenicol, and incubated at 37°C for 2 hr while shaking at 250 rpm. After 2 hr, the starter culture was diluted 1:100 into the main culture of LB containing 100 µg/mL ampicillin and 34 µg/mL chloramphenicol. The main culture was shaken at 37°C and 250 rpm until the OD_600_ reached ∼0.7, then stored at 4°C for ∼30 min while the incubator cooled to 15°C. The culture was then induced with 1 mM IPTG (MilliporeSigma 420322), and grown shaking at 250 rpm for 16 hr at 15°C. After 16 hr, the culture was harvested by centrifugation at 4658 rcf at 4°C for 25 min. The pelleted cells were resuspended in 30 mL Lysis Buffer (500 mM NaCl, 25 mM Tris-HCl pH 8.0, supplemented with 10 mM β-mercaptoethanol and cOmplete, EDTA-free Protease Inhibitor Cocktail (MilliporeSigma (Roche) 5056489001) at 1 tablet/50 mL buffer). The resuspended cells were lysed on ice with 1 mg/mL lysozyme (MilliporeSigma L6876) for 30 min, then sonication. The lysate was then centrifuged at 30,597 rcf at 4°C for 20 min.

A CV of 2.67 mL of Ni-NTA resin (QIAGEN 30250) was utilized per 1 L prep, and the Ni-NTA resin was equilibrated with 10 CV of MilliQ and 6 CV of Lysis Buffer. The clarified supernatant was rotated with Ni-NTA resin for 1.5 hr at 4°C, then centrifuged at 179 rcf at 4°C for 5 min.

The Ni-NTA resin was then washed with 25 CV of Wash Buffer (500 mM NaCl, 25 mM Tris-HCl pH 8.0, 25 mM imidazole, supplemented with 10 mM β-mercaptoethanol), with centrifugations performed at 179 rcf at 4°C for 2 min. The Ni-NTA resin was then resuspended in 2 CV of Wash Buffer and applied to a chromatography column. Protein was eluted with 5 CV of Elution Buffer (500 mM NaCl, 25 mM Tris-HCl pH 8.0, 300 mM imidazole, supplemented with 10 mM β-mercaptoethanol). Eluted protein was pooled and concentrated to ∼10 mL utilizing an Amicon Ultra-15 Centrifugal Filter Unit, MWCO 30 kDa (Millipore UFC9030), by centrifugation at 716 rcf at 4°C. Concentrated protein was centrifuged at 716 rcf at 4°C for 3 min. Dialysis tubing was equilibrated in Dialysis Buffer (25 mM HEPES-NaOH pH 7.0, 5% glycerol, supplemented with 5 mM β-mercaptoethanol) for ∼10 min. The protein was dialyzed in 5 L of Dialysis Buffer overnight, stirring at 4°C.

Dialyzed protein was centrifuged at 716 rcf at 4°C for 10 min. Filtered supernatant was purified using an FPLC with a HiTrap SP XL column, equilibrated in Low-Salt ion exchange (IEX) Buffer (25 mM HEPES-NaOH pH 7.0, 5% glycerol, supplemented with 5 mM DTT). The column was washed with 2 CV of Low-Salt IEX Buffer, then protein was eluted utilizing a 0-80% gradient with Low-Salt IEX Buffer as the base buffer, and High-Salt IEX Buffer (750 mM NaCl, 25 mM HEPES-NaOH pH 7.0, 5% glycerol, supplemented with 5 mM DTT) as the elution buffer. Based on the chromatogram, elution fractions were pooled and concentrated to ∼40 mg/mL utilizing an Amicon Ultra-15 Centrifugal Filter Unit, MWCO 30 kDa (Millipore), by centrifugation at 716 rcf at 4°C. The concentrated protein was supplemented to 50% glycerol with 100% glycerol, then aliquoted, flash-frozen in liquid nitrogen, and stored at −80°C until use.

#### Purification of TDP-43-MBP-His

TDP-43-MBP-His, or MBP-His alone, plasmids were transformed into BL21 (DE3)-RIL *E. coli* and grown on LB-Kanamycin plates at 37°C for 16 hr. The cells were then transferred to a starter culture of LB containing 50 µg/mL kanamycin and 34 µg/mL chloramphenicol, and incubated at 37°C for 4 hr while shaking at 250 rpm. After 4 hr, the starter culture was diluted 1:100 into the main culture of 1 L LB containing 50 µg/mL kanamycin, 34 µg/mL chloramphenicol, and 0.2% glucose. The main culture was shaken at 37°C and 250 rpm until the OD_600_ reached ∼0.25, then continued to grow while cooling to 16°C, and induced with 1 mM IPTG after reaching 16°C and an OD_600_ of ∼0.5-0.6. The induced culture was grown shaking at 250 rpm for 16 hr at 16°C. After 16 hr, the culture was harvested by centrifugation at 4658 rcf at 4°C for 20 min. The pelleted cells were resuspended in 20 mL of Resuspension/Wash Buffer (1 M NaCl, 20 mM Tris-HCl pH 8.0, 10% glycerol, 10 mM imidazole pH 8.0, supplemented with 1 mM DTT, 5 µM Pepstatin A, 100 µM PMSF, and cOmplete, EDTA-free Protease Inhibitor Cocktail at 1 tablet/50 mL buffer). The resuspended cells were lysed on ice with 1 mg/mL lysozyme for 30 min, then sonication. The lysate was then centrifuged at 30,966 rcf at 4°C for 20 min.

A CV of 5 mL of Ni-NTA resin (QIAGEN) was utilized per 1 L prep, and the Ni-NTA resin was equilibrated with 18 CV of MilliQ and 3 CV of Resuspension/Wash Buffer. The clarified supernatant was rotated with Ni-NTA resin for 1 hr at 4°C, then centrifuged at 179 rcf (2000 rpm; 50 mL tubes) at 4°C for 4 min. The Ni-NTA slurry was then applied to a chromatography column, with the flow-through re-applied once. At 4°C, the column was washed with 10 CV of Resuspension/Wash buffer, then eluted in 3 CV of Nickel Elution Buffer (1 M NaCl, 20 mM Tris-HCl pH 8.0, 10% glycerol, 300 mM imidazole pH 8.0, supplemented with 1 mM DTT, 5 µM Pepstatin A, 100 µM PMSF, and cOmplete, EDTA-free Protease Inhibitor Cocktail at 1 tablet/50 mL buffer). Eluted fractions were stored overnight at 4°C.

Eluted fractions were pooled based on purity determined by SDS-PAGE. Approximate protein concentration was determined by Bradford, and 1 mL amylose resin (New England Biolabs E8021L) per 6 mg protein was utilized as the amylose resin CV. Amylose resin was equilibrated in ∼10 CV MilliQ and 3 CV Resuspension/Wash Buffer. The protein was rotated with amylose resin at 4°C for 30 min, then centrifuged at 179 rcf at 4°C for 4 min. The amylose slurry was then applied to a chromatography column, with the flow-through re-applied once. At 4°C, the column was washed with 5 CV of Resuspension/Wash buffer, then eluted in 3 CV of Amylose Elution Buffer (1 M NaCl, 20 mM Tris-HCl pH 8.0, 10% glycerol, 10 mM imidazole pH 8.0, supplemented with 1 mM DTT, 5 µM Pepstatin A, 100 µM PMSF, 10 mM maltose). Eluted fractions were pooled based on purity determined by SDS-PAGE, then concentrated utilizing an Amicon Ultra-15 Centrifugal Filter Unit, MWCO 50 kDa (Millipore UFC9050), by centrifugation at 716 rcf at 4°C, until a concentration of ∼150-200 µM was achieved. The protein was aliquoted, flash-frozen in liquid nitrogen, and stored at −80°C until use.

#### Purification of MBP-FUS

MBP-FUS was purified based on previous protocols (*92*). In brief, MBP-FUS plasmid DNA was transformed into One Shot BL21 Star (DE3) *E. coli* (Thermo Fisher Scientific) cells via heat shock, plated on LB agar plates containing 100 µg/ml ampicillin and incubated overnight at 37°C. The next day, bacterial cultures were scaled-up in LB media supplied with 100 µg/mL ampicillin and 0.2 % glucose and grown to an OD_600_ of 0.6 at 37°C. Expression was induced by adding 1 mM IPTG followed by incubation for 16 hr at 16°C and 250 rpm. Cells were harvested by centrifugation for 20 min at 4658 rcf and 4°C. Cells were resuspended in lysis buffer (20 mM HEPES, pH 7.4, 50 mM NaCl, 2 mM EDTA, 10% glucose, 2 mM DTT), supplied with 20 mg/mL lysozyme and incubated on ice for 30 min. After sonication, the lysate was centrifuged for 20 min at 30,966 rcf and 4°C.

The supernatant was pooled and added to 5 mL amylose beads (New England Biolabs) equilibrated with resuspension buffer and nutated for 2 hr at 4°C. Subsequently, the beads were washed with resuspension buffer and eluted using the same buffer supplied with 10 mM maltose. For further purification and RNA removal, the eluted sample was loaded onto a Heparin column (HiTrap Heparin HP, Cytiva) equilibrated with resuspension buffer using an FPLC. The sample was eluted with a linear gradient ranging from 0-80% high-salt buffer (20 mM HEPES, pH 7.4, 1 M NaCl, 2 mM EDTA, 10% glucose, 2 mM DTT) over 90 mL. Protein-containing fractions were pooled, concentrated using an Amicon spin concentrator (Merck Millipore, MWCO 50 kDa) and flash-frozen in liquid nitrogen.

#### Purification of TDP-43 RRMs utilized for NMR experiments

WT TDP-43 RRMs (102–269) were expressed via a codon-optimized sequence in the pJ411 vector. Protein growth and purification protocols were adapted from the literature (*35*). The protein was grown in BL21 Star (DE3) *E. coli* cells in M9 minimal media supplemented with ^15^NH_4_Cl for isotopic labeling. Bacterial cultures were grown at 37°C with agitation at 200 rpm to an optical density of 0.8. The cultures were induced with 1 mM IPTG and grown for 4 additional hours before harvesting by centrifugation at 6000 rpm and 4°C for 15 min. and resuspended in lysis buffer (20 mM HEPES pH 7.5, 1 M NaCl, 30 mM imidazole, 1 mM DTT) supplemented with an EDTA-free protease inhibitor cocktail (Roche). The cells were lysed using an EmulsiFlex C3 homogenizer (Avestin), and the lysate was cleared by centrifugation at 20,000 rpm and 4°C for 1 hr. The protein was eluted via a nickel HisTrap HP column (GE Healthcare) by affinity chromatography with a linear gradient of elution buffer (20 mM HEPES pH 7.5, 1 M NaCl, 300 mM imidazole, 1 mM DTT). The hexahistidine tag was cleaved by overnight dialysis with 0.03 mg/mL TEV protease at room temperature (20 mM HEPES pH 7.5, 500 mM NaCl, 1 mM DTT). The tag and TEV protease were removed with an additional elution over the HisTrap HP column. The purified protein was buffer exchanged into NMR buffer (20 mM KPi pH 6.8, 1 mM DTT), concentrated to ∼1 mM, flash frozen, and stored at −80°C.

#### RNA oligonucleotides

All RNA oligonucleotides utilized were purchased from Integrated DNA Technologies (IDT) or Horizon Discovery. All RNAs utilized for *in vitro* assays were unmodified and purified with standard desalting (except for RNAs utilized for NMR, which were HPLC purified), and were resuspended in RNase-free water, with nanodrop measurement performed to calculate the RNA concentration. RNAs utilized for cellular experiments were HPLC purified and were fully 2’OMe modified; the Clip34 RNA also had five phosphorothioate backbone modifications on each end of the RNA. RNAs utilized for mouse experiments were purified with *in vivo* HPLC, were fully 2’OMe modified, and had five phosphorothioate backbone modifications on each end of the RNA. A subset of RNA utilized for mouse experiments contained a 5’ Cy5 fluorophore.

#### *In vitro* TDP-43 aggregation prevention assay

RNA was thawed on ice, then serially diluted in water to achieve the desired working concentrations. Protein was thawed on ice, then centrifuged at 21,300 rcf for 10 min. at 4°C. Protein was then buffer exchanged into aggregation assay buffer (166.66 mM NaCl, 22.22 mM HEPES-NaOH pH 7.0, 1.11 mM DTT) using Micro Bio-Spin Chromatography Columns (BIO-RAD 7326200), following manufacturer’s instructions. After buffer exchange, nanodrop measurements were performed to calculate protein concentration. Aggregation assay buffer, and subsequently protein, were added to the tubes containing water/RNA, in order to achieve sample reactions with final concentrations of 5 µM TDP-43, 150 mM NaCl, 20 mM HEPES-NaOH pH 7.0, 1 mM DTT, with varying concentrations of RNA. The reactions were incubated for 15 min at RT after the addition of protein. To a 96-well nonbinding plate (Greiner Bio-One 655906), 0.25 µg TEV protease was added for a final concentration of 2.5 µg/mL, or TEV protease elution buffer for the No TEV control. The reactions were then added to the 96-well plate. The 96-well plate was sealed using parafilm. Turbidity was measured at absorbance 395 nm in a Tecan plate reader (Infinite M1000 or Safire2) for 16 hr, measuring every 1 min. Plate reader measurements were conducted at ambient temperature, typically ∼25-30°C.

For quantification, turbidity data was first standardized by setting the initial value for each well to 0. For the standardized data, any negative values were also set to 0. For normalization, the maximum value of the standardized No RNA condition data for a replicate was set to 100, with all other conditions for that protein in the replicate normalized based on this. Area under the curve (AUC) was calculated for the normalized data. To then normalize the AUC data, the AUC for the No RNA condition was set to 100. This analysis was performed separately for each replicate. The normalized AUC was then used to calculate an IC_50_ value for each replicate, utilizing nonlinear regression: [inhibitor] vs normalized response with variable slope. The IC_50_ value for each replicate was then combined to generate summary data.

#### *In vitro* FUS phase separation prevention assay

RNA was thawed on ice and diluted in water and 2x LLPS buffer to achieve the desired RNA concentration in 1x FUS LLPS buffer (20 mM HEPES-NaOH pH 7.4, 1 mM DTT). Protein was thawed on ice, then centrifuged at 21,300 rcf for 5 min at 4°C. Protein was diluted to 6 µM in FUS elution buffer (570 mM NaCl, 20 mM HEPES-NaOH pH 7.4, 2 mM EDTA, 10% glycerol, 0.5 mM DTT). TEV protease was diluted to 0.12 mg/mL in 1x FUS LLPS buffer. Equal volumes of protein and RNA were mixed, then transferred to a 384-well glass bottom plate (Azenta MGB101-1-2-LG-L). Diluted TEV protease was then added directly to the samples in the 384-well plate, to achieve final concentrations of 2 µM FUS protein, varying concentrations of RNA, 0.04 mg/mL TEV protease, 190 mM NaCl, 20 mM HEPES-NaOH pH 7.4, 0.83 mM DTT, 0.67 mM EDTA, and 3.33% glycerol. Turbidity was measured at absorbance 395 nm in a BMG Labtech plate reader (CLARIOstar Plus) for ∼2-2.5 hr at 26°C, measuring every 1 min. At the endpoint of turbidity measurements, samples were imaged within the plate by brightfield microscopy with a 100x objective (EVOS M5000).

#### Electrophoretic mobility shift assay

Protein was thawed on ice, then centrifuged at 21,300 rcf for 10 min at 4°C. Protein was then buffer exchanged into 150 mM NaCl, 20 mM HEPES-NaOH pH 7.0 (or pH 6.0 where indicated), 10% glycerol, 1 mM DTT using BIO-RAD Micro Bio-Spin Chromatography Columns, following manufacturer’s instructions. After buffer exchange, nanodrop measurements were performed to calculate protein concentration. Protein was diluted in buffer to achieve a working concentration of 50 µM, in EMSA assay buffer (150 mM NaCl, 20 mM HEPES-NaOH pH 7.0 (or pH 6.0 where indicated), 10% glycerol, 1 mM DTT, 20 ng/µL bovine serum albumin (BSA; Thermo Fisher 23209), 2.5 ng/µL yeast tRNA (Thermo Fisher AM7119), 0.4 U/µL RNasin (Promega N2511)). Protein was then serially diluted in EMSA assay buffer to achieve a range of protein concentrations. 20 µM 5’ 6-FAM RNA resuspended in RNase-free water was diluted to 1 µM in EMSA assay buffer (10x working concentration). 10x RNA was then added to protein samples to achieve 100 nM (1x) RNA and a range of protein concentrations in EMSA assay buffer. Samples were incubated at RT for 30 min. During this incubation, 6% DNA Retardation gels (Thermo Fisher EC63655BOX) were pre-run in 0.5x TBE buffer at 150 V for ∼20 min. 1x dye was prepared by dilution of 5x dye (20 mM EDTA, 50% sucrose, 0.25% bromophenol blue) in EMSA assay buffer. After incubation, heparin (MilliporeSigma H3393) was added to each sample to achieve a final concentration of 0.5 mg/mL heparin. 15 µL of 1x dye was loaded in the first lane to monitor sample progression, while 15 µL of undyed sample was loaded in remaining lanes. Gels were run at 150 V for 40 min. Gels were then imaged on a Typhoon Scanner using FAM fluorescence measurement. The signal of bound TDP-43 in each lane was quantified utilizing Image Studio Lite.

#### SDS-PAGE

Samples were diluted in 3x sample buffer (187.5 mM Tris-HCl, 6% SDS, 30% glycerol, 0.05% bromophenol blue, pH 6.8, 1.42 M β-mercaptoethanol) and boiled at 95°C for 5 min. Precision Plus Protein Dual Color Standard (BIO-RAD 1610374) and samples were loaded on Tris-HCl gels (4-15% or 4-20% as indicated) (BIO-RAD 3450027, 3450033), and run at 175 V for 1 hr 15 min. Gels were stained with Coomassie Brilliant Blue, followed by incubation with Destain I (40% methanol, 7% acetic acid), then Destain II (5% methanol, 7% acetic acid) overnight before imaging.

#### Fluorescence 5’ 6-FAM Clip34 3’ BHQ1 assay

Protein was thawed on ice, then centrifuged at 21,300 rcf for 10 min at 4°C. Protein was then buffer exchanged into fluorescence assay buffer (150 mM NaCl, 20 mM HEPES-NaOH pH 7.0, 1 mM DTT) using BIO-RAD Micro Bio-Spin Chromatography Columns, following manufacturer’s instructions. After buffer exchange, nanodrop measurements were performed to calculate protein concentration. Protein was diluted in fluorescence assay buffer to achieve a working concentration of 50 µM. Protein was then serially diluted in fluorescence assay buffer to achieve a range of protein concentrations. 20 µM 5’ 6-FAM 3’ BHQ1 RNA resuspended in RNase-free water was diluted to 1 µM in fluorescence assay buffer (10x working concentration). 10x RNA was then added to a 96-well nonbinding plate (Greiner). Protein samples were then also added to the 96-well plate, to achieve final concentrations of 100 nM 5’ 6-FAM Clip34 3’ BHQ1, and a range of protein concentrations, in 150 mM NaCl, 20 mM HEPES-NaOH pH 7.0, 1 mM DTT. Samples were incubated at RT for 30 min.

Fluorescence was measured in a Tecan plate reader (Spark) at 25°C. Excitation: 475 nm; bandwidth: 15 nm. Emission: 520 nm; bandwidth: 20 nm. A gain value of 80 was used for all trials. The turbidity value at 30 min was utilized. For each protein variant, it was validated that the signal was stable at the 30 min timepoint by measuring after sample addition to the plate, for 1 hr every 1 min, for at least one replicate. For quantification, (F-F_0_)/F_0_ values were calculated for each condition: the average signal for the No RNA condition was subtracted from the signal for a condition, which was then divided by the average signal for the No RNA condition. EC_50_ values were determined from this data, by performing nonlinear regression: [agonist] vs response with variable slope for each replicate.

#### Hydrogen/deuterium-exchange mass spectrometry (HXMS)

RNA was thawed on ice where needed. Protein was thawed on ice, then centrifuged at 21,300 rcf for 10 min at 4°C. For “free” conditions, protein was buffer exchanged into Non-Deuterated Buffer (150 mM NaCl, 20 mM HEPES-NaOH pH 7.0, 1 mM DTT) using BIO-RAD Micro Bio-Spin Chromatography Columns, following manufacturer’s instructions. After buffer exchange, nanodrop measurements were performed to calculate protein concentration. Protein was then diluted in Non-Deuterated Buffer to make a protein sample consisting of 20 µM TDP-43-MBP-His in 150 mM NaCl, 20 mM HEPES-NaOH pH 7.0, 1 mM DTT. Deuterium on-exchange was performed at 25°C by mixing 10 µL of sample with 40 µL of deuterium on-exchange buffer (D_2_O-based; 150 mM NaCl, 20 mM HEPES-NaOD pH 7.0, 1 mM DTT), resulting in a D_2_O concentration of 80%. At the indicated timepoint, the exchange reaction was quenched by addition of 10 µL of ice-cold 250 mM phosphoric acid, to achieve a final pH of pH 2.5. For non-deuterated samples, 10 µL of sample was mixed with 40 µL of Non-Deuterated Buffer, then quenched by addition of 10 µL of ice-cold 250 mM phosphoric acid. For the fully deuterated sample, 10 µL of sample was mixed with 40 µL of on-exchange buffer, incubated at 30°C for ∼18 hr, then quenched by addition of 10 µL of ice-cold 250 mM phosphoric acid. For “Clip34-bound” conditions, all procedures were the same, except that the protein was buffer exchanged into 166.67 mM NaCl, 22.22 mM HEPES-NaOH pH 7.0, 1.11 mM DTT, then diluted into the same buffer along with RNA and water, to achieve final sample concentrations of 20 µM TDP-43-MBP-His and 40 µM Clip34 in 150 mM NaCl, 20 mM HEPES-NaOH pH 7.0, 1 mM DTT.

HX measurements from 20 s to 14.5 hr were performed at pH 7.0. In order to measure less protected, faster exchanging parts of the protein, another set of measurements was performed at pH 6.0. Due to the direct dependence of the intrinsic exchange rate on OH^-^ concentration, these measurements can be put on the same time axis as the pH 7.0 measurements by dividing the actual exchange time by 10 (*94*). For the subset of timepoints done with pH 6-based buffer, all procedures were the same, except that the HEPES-NaOH component of both the Non-Deuterated Buffer and on-exchange buffer was at pH 6.0, and the quench reagent utilized was ice-cold 145 mM phosphoric acid (to achieve a final pH of pH 2.5). All 1 s, 2 s, 6 s, and 18 s timepoints were collected utilizing pH 6.0 buffer; 1 min and 3 min timepoints were collected with some replicates utilizing pH 6.0 buffer and others utilizing pH 7.0 buffer; 20 s, and 10 min and longer timepoints were collected utilizing pH 7.0 buffer. For example, the “1 min” timepoints were measured by 1 min of on-exchange at pH 7.0, or 10 min of on-exchange at pH 6.0. The agreement between these duplicated replicates indicates that the protein structural stability measured by HX is not affected by the pH change. In addition, WT TDP-43 and Clip34 were confirmed to maintain binding at pH 6.0.

For MS analysis, the sample was digested by loading 50 µL onto a homemade pepsin column maintained at 0°C, where pepsin was immobilized by coupling to POROS 20 AL support (Applied Biosystems) and packed into a column housing of 2 mm x 2 cm (64 µL) (Upchurch) (*95*). The protease-generated fragments were then collected onto a TARGA C8 5 µM Piccolo HPLC column (1.0 x 5.0 mm, Higgins Analytical) and separated on a C8 analytical column utilizing a shaped 10-45% Buffer B gradient at 8 µL/min (Buffer A: 0.1% formic acid; Buffer B: 0.1% formic acid, 99.9% acetonitrile). The effluent was electrosprayed into the mass spectrometer. Peptides were identified from non-deuterated samples by MS/MS (Thermo Q Exactive), by analyzing MS/MS data using SEQUEST Proteome Discoverer (ThermoFisher). Deuterated samples, and additional non-deuterated reference samples, were analyzed by MS (Thermo Q Exactive or Thermo Exactive Plus EMR).

HDExaminer software was utilized to process and analyze the HXMS data. The timepoints for samples performed at pH 6.0 were input as one-tenth of the actual on-exchange time. ExMS2, a MATLAB-based program, was used to prepare the peptide pool used by HDExaminer, from the SEQUEST output files for MS/MS data analysis. HDExaminer uses a non-deuterated sample as the reference for identifying deuterated peptides. Manual adjustment of retention times and m/z windows was performed as needed to correct any initial errors. Each deuterated peptide is corrected for back exchange after quenching, by normalizing to the maximal deuteration level of that peptide as detected in the fully deuterated sample. For calculating the peptide deuteration level at each timepoint, HDExaminer identifies the peptide envelope centroid values for both the non-deuterated and deuterated peptides.

#### HXMS data visualization

For visualizing the difference in peptide deuteration levels for each peptide, the HDExaminer data was visualized using MATLAB. At each timepoint, the average deuteration percent for a peptide in either the free or bound condition was calculated by taking the average deuteration percent of all replicates for the peptide at that timepoint that were identified with medium or high confidence. These values were then analyzed in MATLAB by subtracting the average deuteration percent of the peptide in the Clip34-bound state from the average deuteration percent of the peptide in the free state. This data is plotted in MATLAB according to the colors shown in the color legends in the figures (e.g. Fig. 2). Peptides of the same sequence but different charge states are plotted to allow visualization of the agreement across separate charge states for a unique peptide sequence.

To generate plots of consensus exchange difference for each timepoint, the exchange differences of the peptides at that timepoint were manually analyzed, with the consensus exchange difference determined based on the average classification of all peptides including a specific residue. This was done via a scoring system, where a peptide with a difference of less than 10% receives a score of 0, a peptide with a difference of ≥ 10% (light blue according to legend) receives a score of −1, a peptide with a difference of ≥ 20% (medium blue) receives a score of −2, a peptide with a difference of ≥ 30% (dark blue) receives a score of −3, a peptide with a difference of ≤ −10% (light red) receives a score of +1, a peptide with a difference of ≤ −20% (medium red) receives a score of +2, and a peptide with a difference of ≤ −30% (dark red) receives a score of +3. As peptides were binned according to this scoring system, the displayed consensus percentage differences in exchange do not report the exact value of the percentage difference for each peptide, and do not report the proximity of each peptide’s behavior to the cutoff value for each score. The average score for each residue was rounded to the nearest whole number, and this data was plotted in GraphPad Prism, with the rounded score value for each residue colored according to the same scoring system as described for the manual analysis above.

To generate plots of HX data for representative peptides displaying exchange as the number of deuterons, the HDExaminer output of this data was visualized using GraphPad Prism. For a peptide, the values for the number of deuterons from HDExaminer was taken for each replicate with high or medium confidence at each timepoint, for both free and bound states. This was then plotted in GraphPad Prism.

To generate plots of mass spectra, HDExaminer output was visualized in GraphPad Prism. The raw mass spectrum data for a particular replicate of a specific timepoint in either the free or bound state was copied into GraphPad Prism. This data was manually analyzed to determine the signal that corresponded to the desired peptide, based on the signal corresponding to the appropriate m/z values. All signal is displayed in the mass spectra plots, but the signal determined to correspond to the correct peptide is colored red, for ease of visualization. Dashed guidelines for visualization are also displayed, with the blue line corresponding to the value of the monoisotopic peak for that peptide, and the purple line corresponding to the centroid value of the peptide in the fully deuterated condition. Representative spectra for each state at each timepoint were chosen by determining the average value of the centroid for all replicates with peptides of high or medium confidence, for that state and timepoint. The spectrum displayed as the representative spectrum corresponds to the replicate with the centroid value closest to the average of these centroid values.

#### Sedimentation analysis

At the end timepoint of aggregation prevention assays (t = 16 hr), a portion of select conditions was transferred to a tube. Tubes were spun at 21,300 rcf for 10 min at RT to sediment the pellet. The supernatant was transferred to a fresh tube. The pellet was resuspended in assay buffer of equal volume. Equal volumes of supernatant from multiple conditions were then run on SDS-PAGE gels. To determine the relative amounts of protein in the supernatant, Image Studio Lite was utilized to quantify the TDP-43 band signal for supernatant samples for each condition.

#### NMR data collection and processing

All NMR experiments were heteronuclear single quantum coherence (HSQC) spectra conducted on Bruker Avance 600 MHz ^1^H Larmor frequency spectrometers with HCN TCI z-gradient cryoprobe at 298K. TDP-43 RRMs NMR samples contained 50 µM protein in 20 mM KPi pH 6.8, 1 mM DTT, 5% D_2_O (v/v). NMR samples of RRMs with RNA included 100 µM (2x molar equivalent) RNA. Backbone chemical shift assignments were transferred from BMRB deposited data (BMRB ID 27613). NMR data were processed and analyzed with Bruker TopSpin, NMRPipe (*96*), and CCPNMR (*97*). Chemical shift perturbations were quantified by comparison of the ^1^H–^15^N cross-peak measurements in the RNA-containing and RNA-free HSQCs. Intensity ratios were calculated from the intensity of the ^1^H–^15^N cross-peaks with the formula I/I_o_ where I is the RNA-containing sample and I_o_ is the RNA-free control sample.

#### G-quadruplex RNA annealing

RNA was thawed on ice. RNA was diluted to achieve working concentrations of 20 µM RNA, 150 mM NaCl, 20 mM HEPES-NaOH pH 7.0, 1 mM DTT. RNA was then annealed in a PCR machine by heating at 95°C for 2 min, followed by decreasing temperature at a rate of 1°C per minute, until reaching RT. RNA was added to the desired assay within a maximum of 30 min after the end of the annealing process.

#### Circular Dichroism

RNA was first prepared as described in the above “G-quadruplex RNA annealing” methods section. RNA was then diluted to achieve a final concentration of 5 µM RNA, 150 mM NaCl, 20 mM HEPES-NaOH pH 7.0, 1 mM DTT. The absorbance spectra were recorded in a 1 mm pathlength cuvette at 25°C with an Aviv Circular Dichroism Spectrometer, Model 202.

Parameters for measuring the spectra were a measurement range of 220-320 nm, a bandwidth of 2 nm, a wavelength step of 2 nm, and an averaging time of 60 s. Data was standardized by subtracting the absorbance spectrum of the blank. The standardized data was then normalized utilizing the equation: Δε(M^-1^cm^-1^) = θ/(32980*c*l), where θ is the reported CD signal in millidegrees, c is the RNA molar concentration, and l is the pathlength in cm.

#### HEK293 cell oligonucleotide treatment

Glass bottom 24-well plates were coated with 50 µg/mL collagen overnight. OptoTDP-43 stable HEK293 cells were plated at 150,000 cells/well in DMEM (Fisher Scientific) with 10% BGS (Hyclone; Fisher Scientific) and 1% Glutamax (Thermo Fisher). 16 hours after plating, optoTDP-43 expression was induced by media change to phenol-free DMEM/10%BGS/1%Glutamax with 750 ng/mL doxycycline-hyclate. Immediately after the media change, oligo treatments were started. 2’OMe_RNA oligos were transfected using lipofectamine RNAiMAX according to manufacturer’s instructions (Invitrogen). Briefly, 500 nM of each oligo was diluted in OptiMEM (Thermo Fisher) and mixed with 1 µL lipofectamine per well, incubated at room temperature for 10 minutes, and added dropwise to the cells. Plates were loosely wrapped in aluminum foil to prevent light exposure and subsequent light-induced TDP-43 oligomerization. 43 hours after doxycycline-hyclate induction, plates were removed from the foil and placed on an LED array (Amuza) for blue light stimulation (465 nm) for 5 hours at 37°C. After blue light stimulation, cells were washed once with PBS and fixed with 4% PFA (in PBS) for 20 minutes at room temperature. Cells were permeabilized in 0.3% Triton X-100 in PBS and stained with Hoechst (1:1000; Thermo Fisher) overnight.

#### HEK293 cell imaging and analysis

Image acquisition was performed using a Nikon Eclipse Ti2 Inverted Microscope with a 40X air objective. 20 fields of view (FOVs) were randomly selected by the NIS-Elements software per well. All image visualization and quantification were performed using NIS-Elements AR1 Analysis 4.51. The microscopy images were collected across three independent experiments and maximum intensity projection images were used for analysis. Binary thresholds (594 nm and 405 nm channel) and spot detection were used to capture and separate nuclei and puncta objects.

Puncta overlapping with the nuclear signal were removed from the analysis, leaving only cytoplasmic puncta for quantification. Puncta area (µm^2^) was divided by nuclei count and expressed as puncta area per cell. Puncta area/cell was normalized against the average value for the control (CTR) oligo. Out-of-focus images were removed, and FOVs with mean puncta area >100 µm^2^/cell were excluded and considered outliers. Twenty FOVs were analyzed per well, and at least two to three wells were imaged per experiment. Mean values per experiment were normalized to control oligonucleotide treatment and considered a biological replicate, and the mean values of three biological replicates (n=3 experiments) were used to analyze the effect of the oligonucleotide.

#### Stable HEK293 CUTS cell line

Stable HEK293 cells expressing CUTS were generated as previously described (*82*). Briefly, HEK293 cells were seeded in 6-well plates and transfected at roughly 70% confluency with 2.5 μg of PiggyBac plasmids encoding CUTS along with 0.5 μg of a Super PiggyBac Transposase Expressing plasmid (PB200PA-1), using Lipofectamine 3000 (Invitrogen) as per the manufacturer’s instructions. A control group lacking the transposase plasmid was included. After 48 hours, cells were subjected to selection with 5 μg/mL puromycin (Sigma, P8833), with media being refreshed every two days. Non-transfected control cells typically died within five days under selection. Surviving cells were expanded and cultured in media containing a reduced puromycin concentration (2.5 μg/mL) to establish stable cell lines. Successful transgene expression was validated through live imaging.

#### Live Confocal Microscopy of HEK CUTS cell line

Live-cell imaging was carried out using a Nikon A1 laser-scanning confocal microscope equipped with a 10X objective lens. Environmental conditions during imaging were maintained using a Tokai HIT stage-top incubator. Images were acquired and analyzed using Nikon Elements software. Representative images were selected from a minimum of three independent experiments to ensure reproducibility.

#### siRNA Reverse Transfection and RNA oligonucleotide transfection of HEK CUTS cell line

siRNA-mediated gene knockdown was performed via reverse transfection using Lipofectamine RNAiMAX reagent (Invitrogen, 13778150), following the manufacturer’s instructions. To reduce TDP-43 expression, ON-TARGETplus SMARTpool siRNA targeting *TARDBP* (Dharmacon, L-012394-00-0005) was employed. Non-targeting siRNA (Dharmacon, D-001206-13-05) served as a control in these experiments. RNA oligonucleotide transfections were performed using Lipofectamine RNAiMAX reagent (Invitrogen) in accordance with the manufacturer’s protocol.

#### iPSC-derived neuron treatment and immunostaining

RNA treatments started on day 13 after plating (DIV27) and lasted 24 hr. RNAs were transfected using Lipofectamine RNAiMAX (Invitrogen) according to the manufacturer’s instructions.

Briefly, each RNA was diluted in OptiMEM (Gibco) and combined with 1 µL Lipofectamine per well, also diluted in OptiMEM. The mixture was incubated at RT for 10 min, and then added dropwise to the cells with each RNA at a final concentration of 500 nM. Neurons were fixed 24 hr after RNA treatment on day 14 after plating (DIV28).

On DIV28, cells were washed once in PBS (Gibco) and fixed in 4% paraformaldehyde (PFA) (Electron Microscopy Sciences) immediately after treatments ended. Cells were kept in PFA for 20 min, then washed three times in PBS and blocked with 5% Donkey Serum (Jackson ImmunoResearch) + 0.3% TX-100 (Sigma-Aldrich) in PBS for 30 min at RT. Primary antibodies (goat MAP2 1:1000, Phosphosolutions; rabbit TDP-43 1:300, Proteintech) were diluted in blocking solution and incubated overnight at 4°C. Secondary antibodies (donkey Alexa Fluor, Jackson ImmunoResearch) were used at 1:1000 dilution in blocking solution and incubated for 60 min at RT. All treatments and cell lines were treated and probed simultaneously to decrease variability. Coverslips were mounted on slides using Prolong Glass mounting media (Invitrogen).

Images were acquired (20 per group) using an A1R Nikon Confocal Microscope and fields of view (FOV) were processed for analyses using Nikon NIS Elements Software. Settings were kept consistent across treatments. Within each FOV, neurons that were isolated (not over glial cells or other neurons) were selected and nuclear TDP-43 signal was measured by overlaying an ROI using DAPI as a guide. Cytosolic area was hand-drawn using MAP2 signal as a guide. Raw intensity values for nuclear signal in the 488 channel (TDP-43) were normalized against cytosolic intensity values and the output was referred to as “nuclear/cytosolic ratio.”

#### Virus Production

AAV9 virus was generated by Vector Biolabs using a plasmid designed as follows. The CMV-promoter driven pcDNA3.2 TDP-43 NLS1 YFP plasmid, a gift from Aaron Gitler (Addgene plasmid #84912; RRID:Addgene_84912) (*84*), was packaged into AAV9 viral particles.

#### Intraspinal delivery of AAV9 Virus

Intraspinal delivery of AAV9 in p180 mice was carried out as previously described (*83*). Mice deeply under anesthesia underwent an incision of their dorsal skin and underlying muscle with retraction, revealing the spinous processes between vertebrae C2 and T1. Following laminectomy at spinal levels C4, C5, and C6, six total bilateral injections were given across this area. Each injection contained 1×10^11^ GC of the AAV9-TDP-43 NLS1 virus (TDP-43^ΔNLS^) in a 1 μL total volume. A gas-tight Hamilton syringe mounted on a UMP3 electronic micropump (World Precision International) was used for these injections, with a 33-gauge 45° beveled needle. Targeting of injections was guided on the lateral axis by the midpoint of each spinal segment, and on the rostral-caudal axis by the location of dorsal root entry for C4, C5, and C6.

The needle was lowered to a depth 0.8 mm below the dorsal surface for ventral horn targeting, with injections then delivered over a 5-minute interval at a constant rate. Sham surgery control animals underwent identical procedures and laminectomies, as well as needle placement and insertion. In sham animals, the Hamilton syringe was filled with sterile PBS and the micropump was not initiated. Following the final injection, the dura was removed from the dorsal spinal cord of the injection region, and a non-adhering dressing (Adaptic non-adhering dressing by Systagenix) was applied. Overlying muscles were then closed in layers, using sterile silk sutures. The skin incision was also closed using both sutures and sterile wound clips. Animals recovered on a heating pad until awake, and were then returned to their home cage. To minimize pain and distress, at the time of surgery and at 12-hour intervals for the first 24 hours following surgery, animals were given subcutaneous sterile saline for fluid balance, buprenorphine analgesic (0.05 mg/kg), and cefazolin antibiotic (10 mg/kg). Animals were monitored daily and were checked for signs of pain and/or distress, as well as ambulatory potential and ability to obtain food/water.

#### Spinal delivery of sRNA

After one week of viral expression, animals were again deeply anesthetized. The original surgical incision was reopened and skin and muscle retracted to expose the spinal cord. The non-adhering dressing was removed from the spinal cord surface, and was replaced with a pre-saturated gelfoam sponge (sterile gelfoam Dental sponge, Pharmacia & Upjohn). TDP-43^ΔNLS^-expressing animals were randomized into saline control, or RNA treatment groups, with the sponge being pre-soaked either in sterile saline solution, or sterile saline reconstituted RNA at a 100 μg/mouse dosing. This dosing penetrates to the ventral spinal cord by 3-days post-application in a robust and reproducible manner, and causes no detrimental side-effects when evaluated out to two-weeks post-administration. Sham surgery control animals underwent identical procedures, and received a saline-saturated gelfoam sponge. Following this application, overlying muscles and skin were again closed with sterile silk sutures, with the skin also being bound with sterile wound clips. Animals again recovered on a heating pad until awake, prior to return to home cages. Animals received the same set of compounds to minimize pain and distress, at the time of surgery and at 12-hour intervals for the first 24 hours following surgery: subcutaneous sterile saline for fluid balance, buprenorphine analgesic (0.05 mg/kg), and cefazolin antibiotic (10 mg/kg). Animals were again monitored daily for signs of pain and/or distress, as well as ambulatory potential and ability to obtain food/water. Assessment of motor neuron numbers was carried out at 3-, 5-, and 7-days following sRNA application, to determine potential beneficial therapeutic effects and longevity.

#### Animal Harvesting for Spinal Cord Immunofluorescence

Mice were euthanized using carbon dioxide asphyxiation, and perfused and fixed following standard laboratory procedures. A perfusion needle connected to a peristaltic pump was inserted into the left ventricle of the heart, and animals were then perfused with approximately 20 mL of PBS followed by 25 mL 4% paraformaldehyde. The animal was dissected to obtain the cervical spinal cord, which was then placed in 4% paraformaldehyde overnight. Paraformaldehyde was briefly rinsed off the tissue with PBS, with spinal cords then placed in 30% sucrose until tissue sinking (24-48 hours). Spinal cords were frozen into Tissue-Tek OCT solution and were subsequently sectioned using a Cryostar NX50 cryostat (epredia) at a section depth of 30 mm. Sections were placed onto charged glass slides. Tissue blocking, permeabilization, and staining were performed according to laboratory standard protocols and according to antibody manufacturer recommendations. 30 µm spinal cord sections on slides were heated overnight at 55°C and were then rinsed with PBS. Sections were next blocked in 5% BSA for 1 hour at room temperature, and then incubated in primary antibodies at 4°C overnight (NeuN) or for 48 hours (ChAT). Primary antibodies included: anti-ChAT (Millipore RRID:AB_2079751, 1:1,000) and anti-NeuN (Cell Signaling Cat# 24307, RRID:AB_2651140, 1:400). Following this incubation and PBS washing, secondary labeling for visualization was attained with AlexaFluor594 (Life Technologies). To label cell nuclei, Hoechst stain (ThermoFisher) was used. Slides were mounted with coverslips using Citifluor AF3 (Electron Microscopy Sciences). Microscopy imaging was accomplished using a Nikon A1+ confocal microscope and NIS-Elements software. ChAT^+^ and NeuN^+^ cells were assessed using bilateral ventral horn images with manual counting by a blinded assessor. TDP-43 puncta, visualized using the YFP tag of the virally expressed protein, were assessed using three-dimensional Imaris Imaging Software (Version 10.1.0, Oxford Instruments).

#### Animal Data Analysis Considerations

Animals were numbered according to surgery order and were randomized into treatment groups. Counting of motor neuron numbers and TDP-43 puncta were performed by a second, blinded individual.

#### Quantification and Statistical Analysis

All statistical details of experiments can be found in the figure legends. Data visualization and statistical analyses were performed with GraphPad Prism (GraphPad Software Inc.; La Jolla, CA, USA). Quantification of gel images was performed with Image Studio Lite (LI-COR Biosciences; Lincoln, NE, USA). HX analysis was performed with HDExaminer (Trajan Scientific and Medical; Melbourne, Australia). MATLAB was utilized for some HX data visualization (MathWorks; Natick, MA, USA). ExMS2, a MATLAB-based tool, was utilized to prepare the peptide pool for HX (*98*). Immunofluorescence analysis was performed utilizing NIS-Elements (Nikon; Minato City, Tokyo, Japan) and Imaris (Oxford Instruments; Abingdon, United Kingdom).

---

**Fig. S1.**
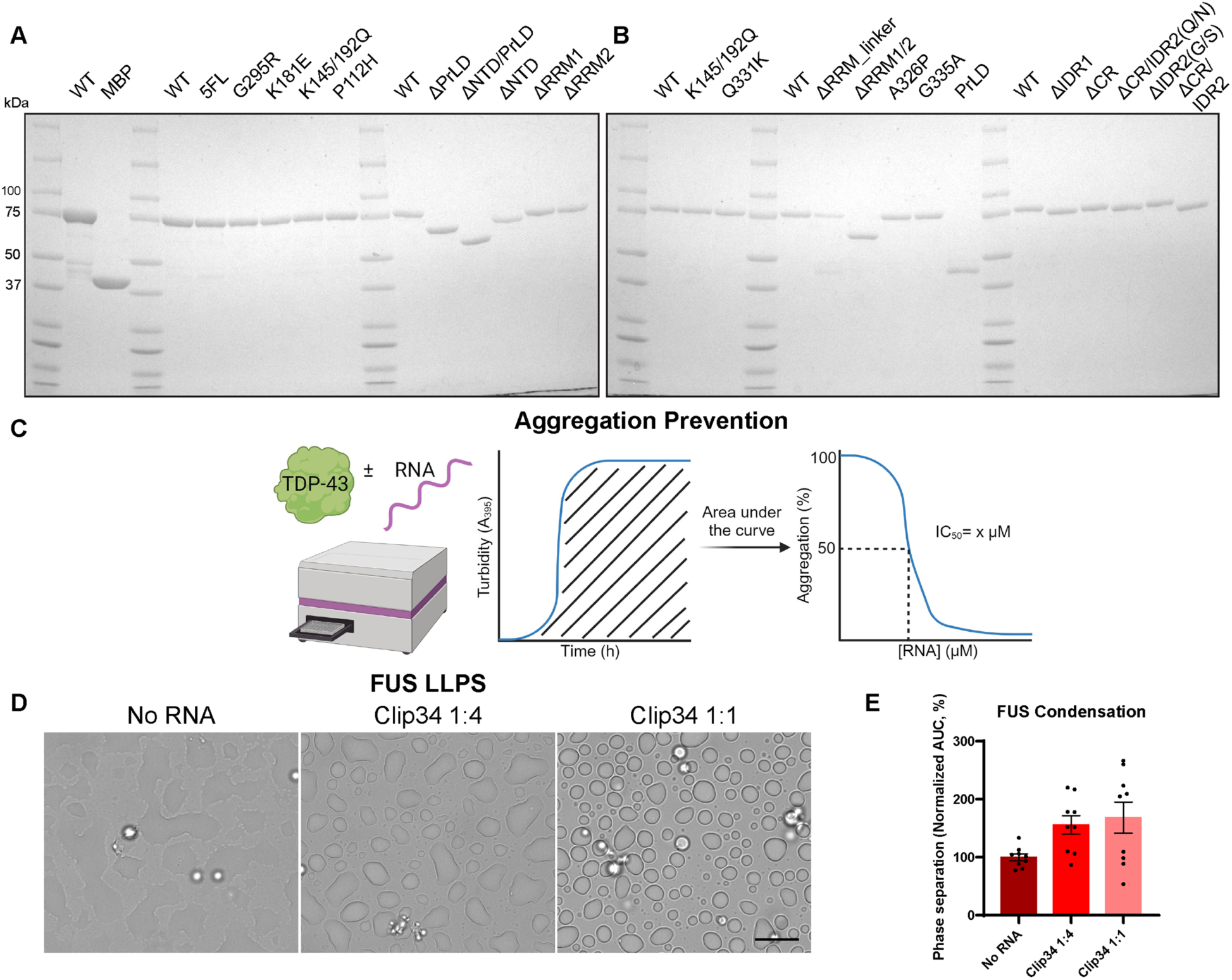
TDP-43 purification, aggregation assay, and inefficacy of Clip34 against FUS phase separation. **(A,B)** 4-15% Tris-HCl SDS-PAGE gels loaded with 1 µg of each indicated purified TDP-43-MBP-His variant (or MBP-His alone) and subsequently stained with Coomassie Brilliant Blue. **(C)** Schematic of *in vitro* TDP-43 aggregation prevention assay. TDP-43-MBP-His (5 µM) was incubated with TEV protease in the presence or absence of RNA for 16 hr, measuring every minute in a plate reader. The standardized turbidity data is normalized so that the No RNA condition maximum value is set to 100; subsequently, the normalized area under the curve (AUC) of this data is taken, which is then utilized to calculate an IC_50_ value (nonlinear regression: [inhibitor] vs normalized response with variable slope). **(D)** Representative 100x brightfield microscopy images of FUS condensates in the presence or absence of Clip34 RNA, after ∼2-2.5 h of measurement in the plate reader. The scale bar represents 10 µm (n=9; 3 biological replicates, each consisting of 3 technical triplicates; [RNA]:[FUS]; 2 µM FUS). **(E)** AUC of turbidity data for the same FUS samples imaged in (D), normalized to the average No RNA value for the respective technical triplicate. Data are mean ± SEM (n=9; 3 biological replicates, each consisting of 3 technical triplicates; [RNA]:[FUS]; 2 µM FUS).

**Fig. S2.**
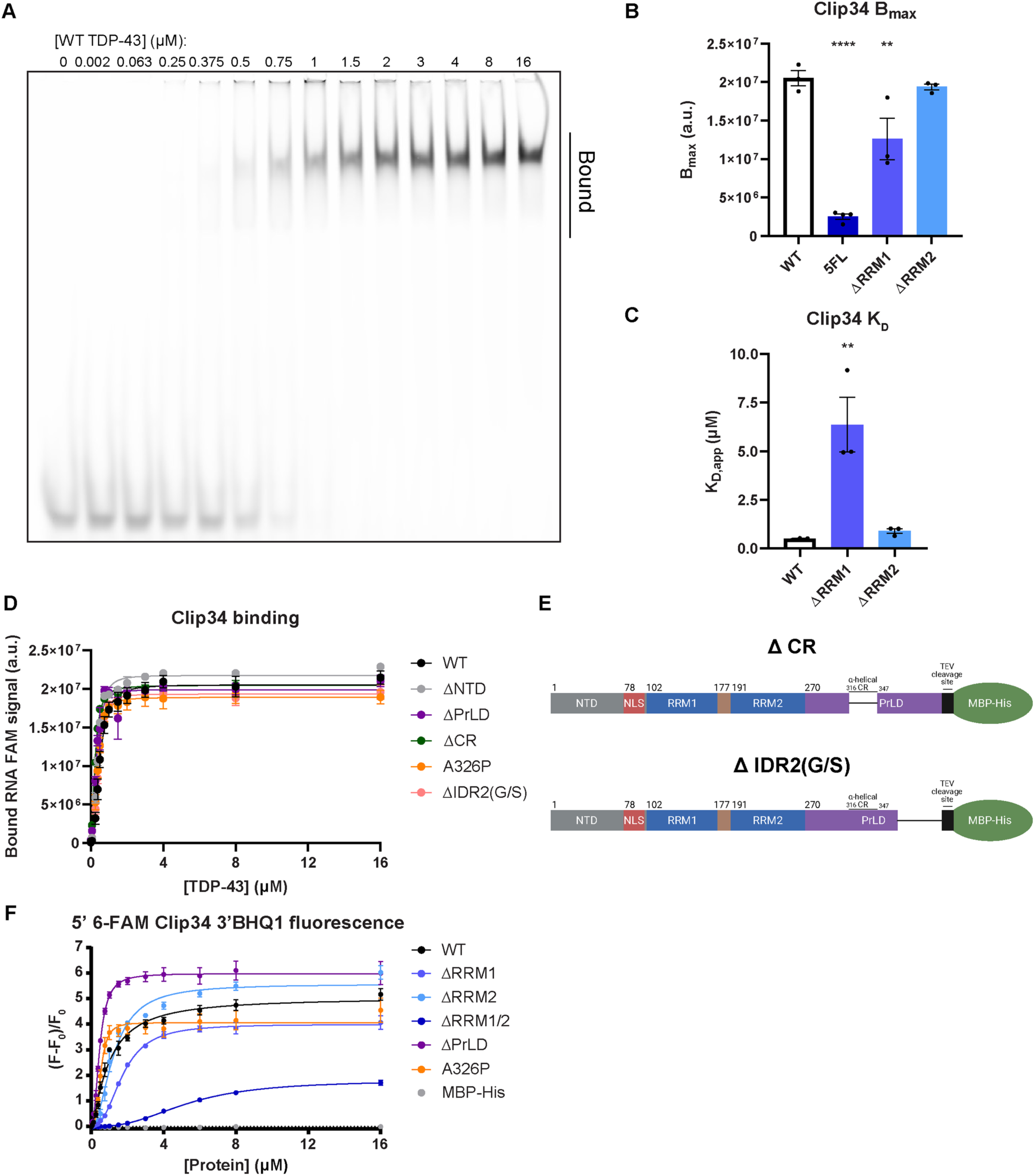
RNA binding and remodeling activity of TDP-43 variants. **(A)** Representative image of an EMSA for 100 nM 5’ 6-FAM Clip34 with indicated WT TDP-43-MBP-His concentrations, run on a 6% DNA Retardation gel. The vertical bar indicates the approximate area quantified for the bound fraction. **(B, C)** B_max_ (B) and apparent K_D_ (C) values calculated from the bound signal from individual replicates of EMSAs performed with 5’ 6-FAM Clip34 and WT TDP-43-MBP-His or the indicated TDP-43 variant, corresponding to summary data in Fig. 1J. WT data shown is the same data as in Fig. 1L. Data are mean ± SEM (n=3-4; one-way ANOVA with Dunnett’s correction comparing to WT; **p≤ 0.01; ****p ≤ 0.0001). **(D)** Bound 5’ 6-FAM Clip34 signal for EMSAs performed with indicated TDP-43-MBP-His variants. WT data shown is the same in Fig. 1J. This is the summary data corresponding to K_D_ values calculated from individual replicates, shown in Fig. 1L. Data are mean ± SEM (n=3; shown is the nonlinear regression: [agonist] vs response with variable slope, of the combined replicates). **(E)** Domain maps of TDP-43 partial PrLD deletion constructs. Amino acids deleted, inclusive, are: ΔCR, aa316-346; ΔIDR2(G/S), aa367-414. **(F)** Relative fluorescence intensity values for 5’ 6-FAM Clip34 3’ BHQ1 with indicated TDP-43-MBP-His variants or MBP-His. This is the summary data corresponding to EC_50_ values calculated from individual replicates, shown in Fig. 1N. Data are mean ± SEM (n=3 for TDP-43 variants; n=2 for MBP-His; 100 nM RNA; shown is the nonlinear regression: [agonist] vs response with variable slope, of the combined replicates).

**Fig. S3.**
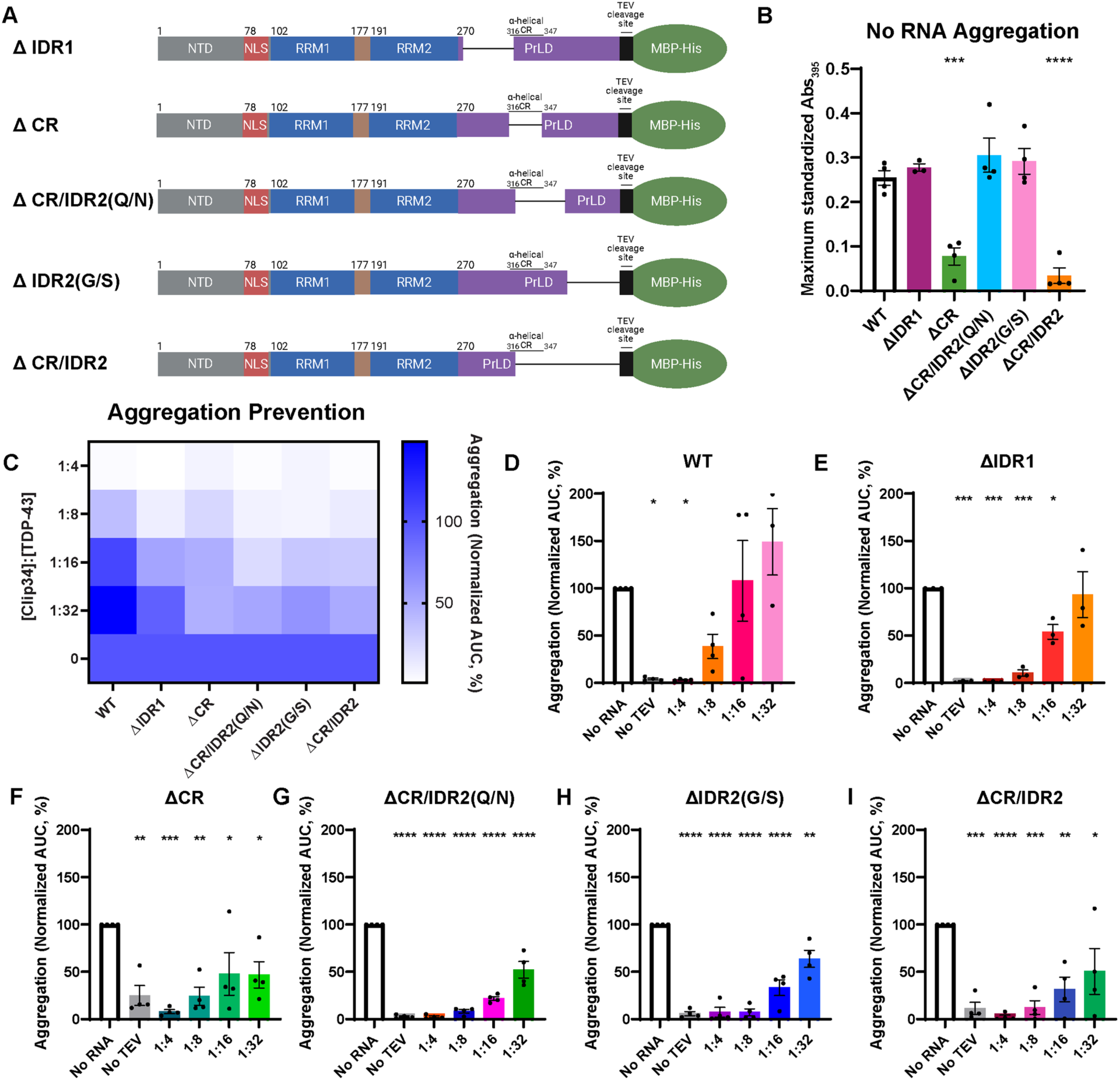
Clip34 exhibits enhanced chaperone activity against partial PrLD-deletion variants. **(A)** Domain maps of the TDP-43 partial PrLD deletion constructs. Amino acids deleted, inclusive, are: ΔIDR1, aa274-320; ΔCR, aa316-346; ΔCR/IDR2(Q/N), aa321-366; ΔIDR2(G/S), aa367-414; and ΔCR/IDR2, aa321-414. **(B)** Maximum standardized Abs_395_ value recorded for the No RNA condition for each TDP-43 variant tested by *in vitro* aggregation prevention assays. Data are mean ± SEM (n=3-4; one-way ANOVA with Dunnett’s correction comparing to WT; ***p ≤ 0.001, ****p ≤ 0.0001). **(C)** Heatmap displaying the mean AUC values from *in vitro* aggregation prevention assays for each TDP-43 variant at the indicated molar concentration ratios, normalized to the respective variant’s No RNA condition. **(D-I)** Individual replicates of the summary data shown in (C) for each TDP-43 variant. Data are mean ± SEM (n=3-4; one-way ANOVA with Dunnett’s correction comparing to No RNA; *p ≤ 0.05, **p ≤ 0.01, ***p ≤ 0.001, ****p ≤ 0.0001).

**Fig. S4.**
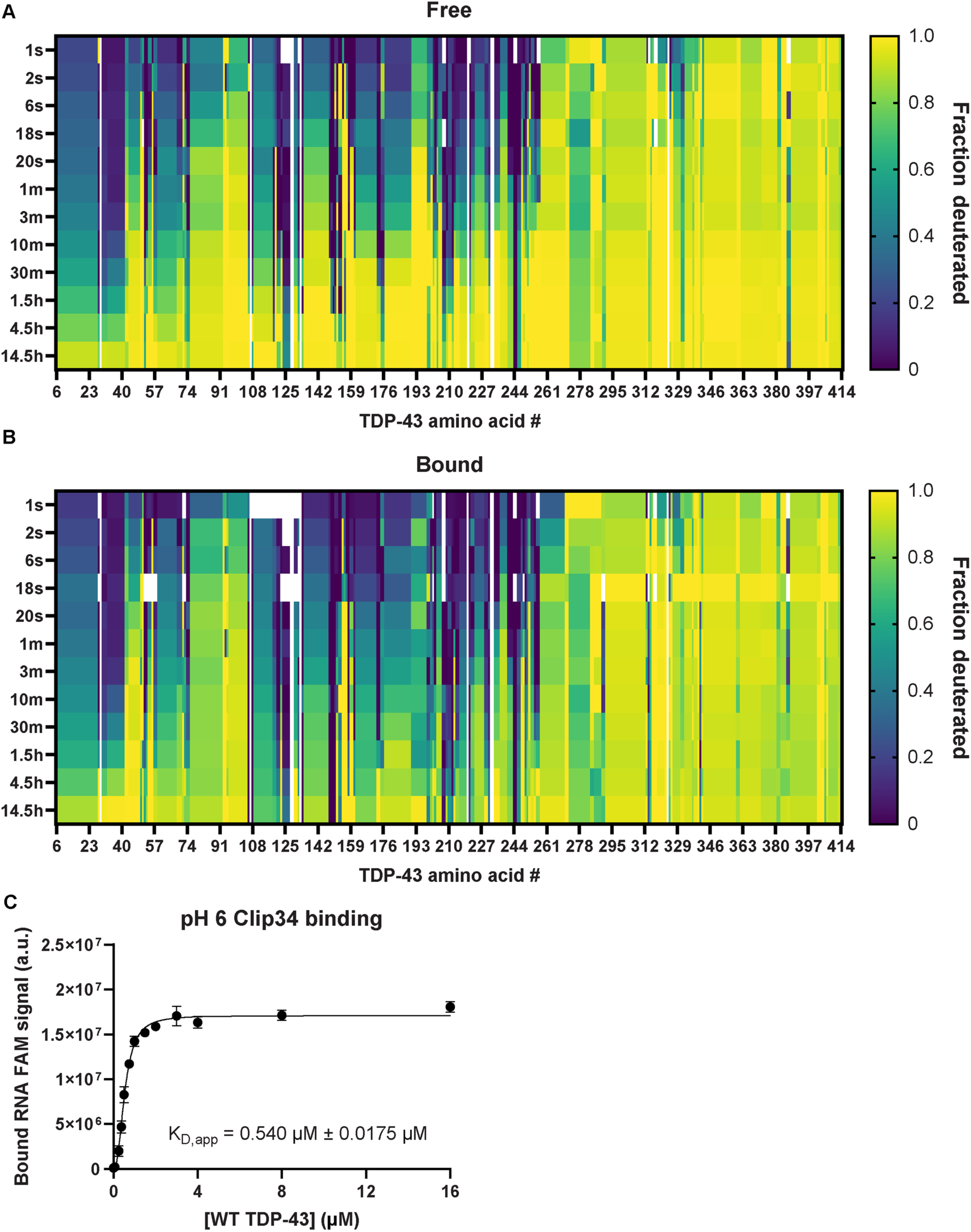
Hydrogen/deuterium-exchange for TDP-43 in the absence or presence of Clip34 RNA. **(A,B)** Heatmaps displaying hydrogen/deuterium-exchange for each TDP-43 amino acid in the free (A) and bound (B) states at each timepoint, normalized so that 1 represents fully deuterated and 0 represents not deuterated. White spaces represent small gaps in coverage. Sub-localization between peptides to determine the exchange value at each amino acid was performed by HDExaminer. All 1 s, 2 s, 6 s, and 18 s timepoints were collected utilizing pH 6.0 buffer; 20 s, 1 min, and 3 min timepoints were collected with some replicates utilizing pH 6.0 buffer and others utilizing pH 7.0 buffer; 10 min and longer timepoints were collected utilizing pH 7.0 buffer. **(C)** Bound 5’ 6-FAM Clip34 signal for EMSAs performed with WT TDP-43 in pH 6.0 buffer. Data are mean ± SEM (n=3; shown curve is the nonlinear regression: [agonist] vs response with variable slope, of the combined replicates; apparent K_D_ value was calculated from individual replicates).

**Fig. S5.**
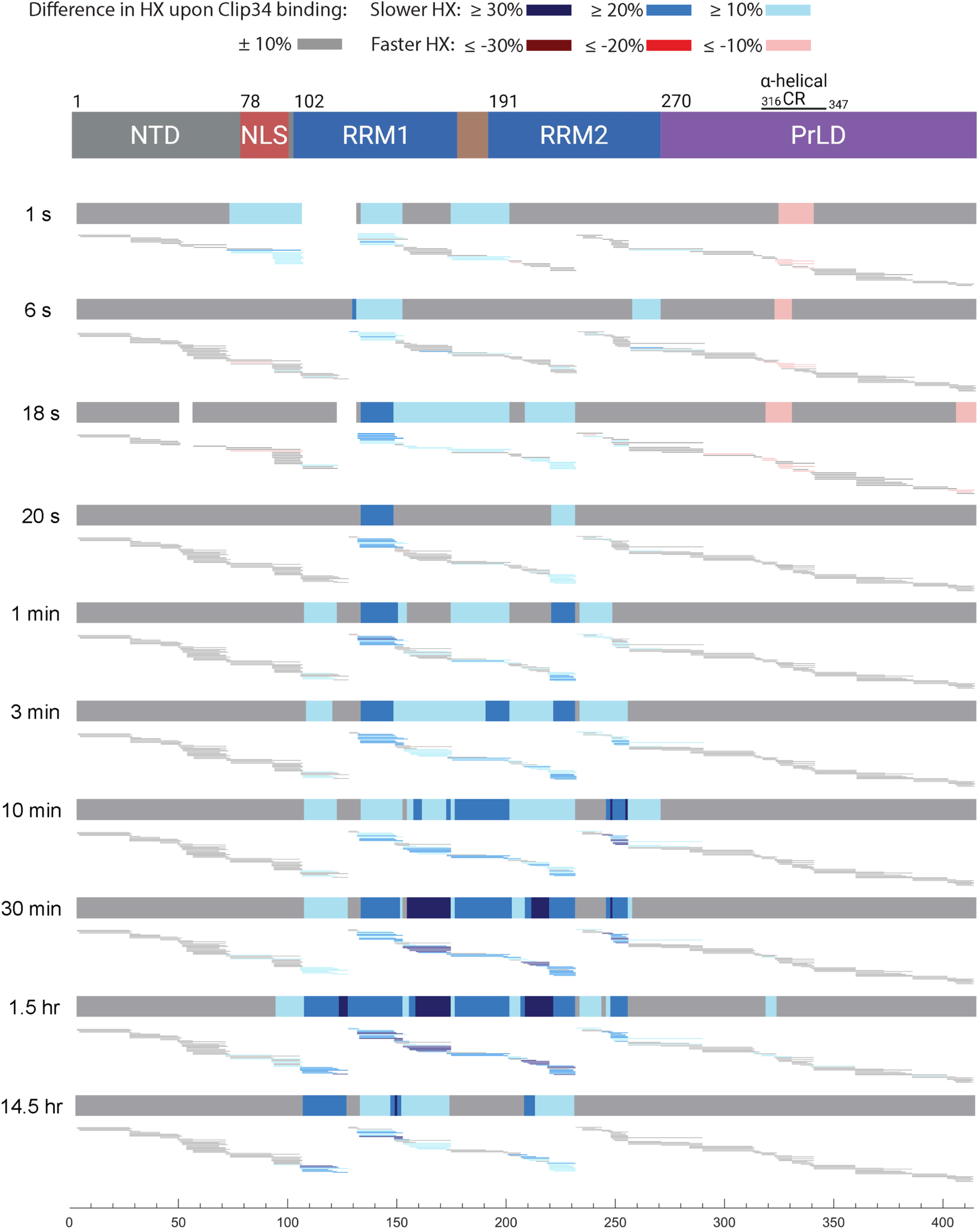
Hydrogen/deuterium-exchange reveals stabilization of the TDP-43 RRMs and destabilization of the α-helical conserved region in the PrLD upon Clip34 RNA binding. *Top:* domain map of TDP-43, aligned with data underneath. *Bottom:* The consensus percentage difference in exchange between the Clip34-bound and free states at each indicated timepoint is shown again as in Fig. 2A. Aligned beneath each consensus percentage difference in exchange plot are the peptides analyzed at the respective timepoint, with percentage differences in exchange between the Clip34-bound and free states for each peptide colored as shown in the legend. The amino acid number is indicated on the axis below.

**Fig. S6.**
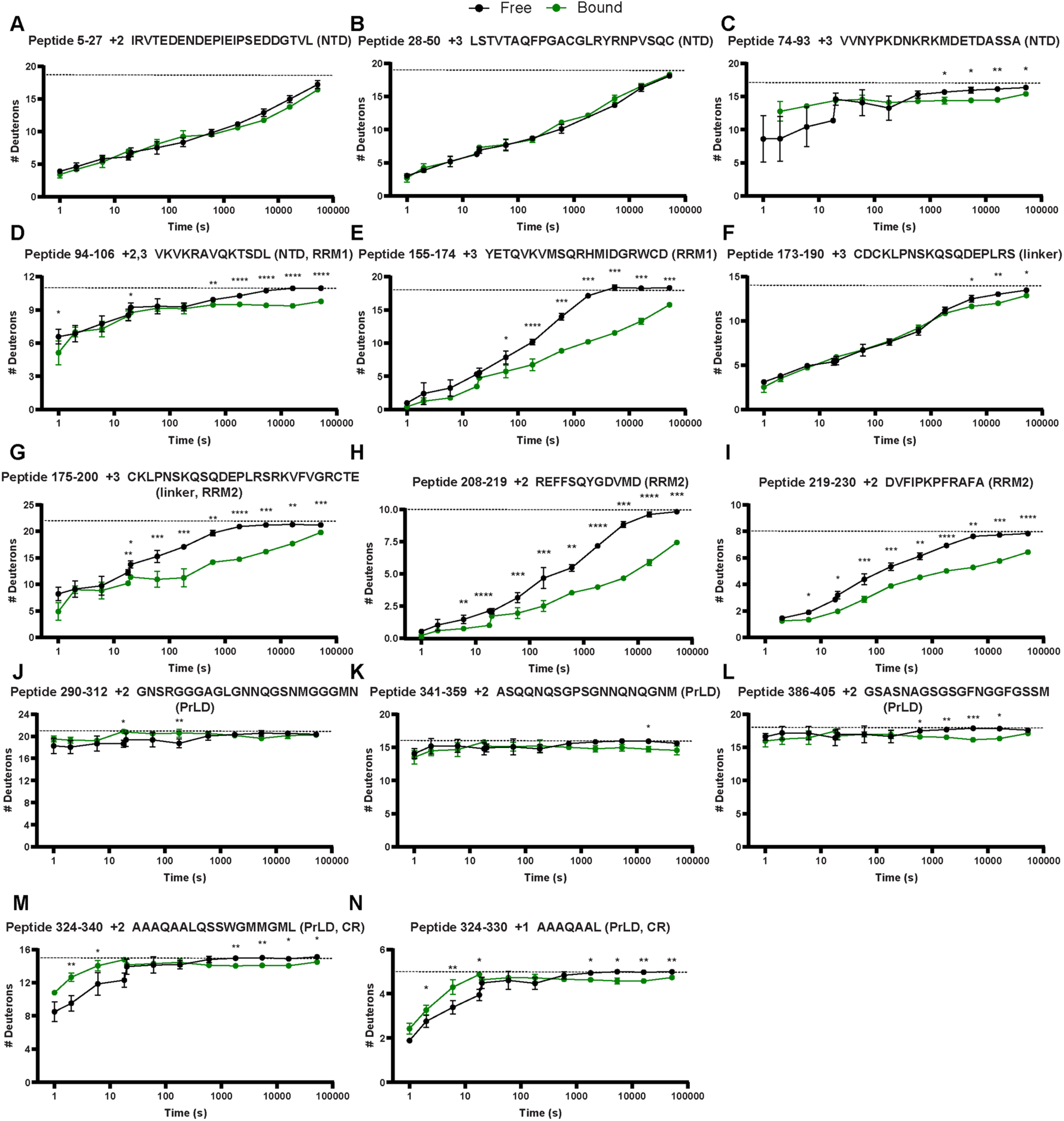
Hydrogen/deuterium-exchange deuteration curves illustrate the effects of Clip34 RNA binding on TDP-43 structure. (A-N) HX for representative peptides, with charge state and peptide sequence indicated in the graph titles. Representative examples are shown of peptides in the NTD (A-C); bridging the NTD and RRM1 (D); in RRM1 (E); bridging RRM1 and the linker between RRMs (F); bridging RRM1, the linker between RRMs, and RRM2 (G); in RRM2 (H, I); in the PrLD primarily outside of the CR (J-L); and primarily within the α-helical conserved region (CR) of the PrLD (M, N). Other regional features of interest include peptides with residues in the nuclear localization sequence (NLS) (C, D) and peptides located in different parts of the PrLD: within IDR1 (J), at the end of the CR and extending into IDR2 (K), and within the glycine- and serine-rich region of IDR2 (L). The dashed line represents the fully-deuterated condition. Data are mean ± SD (n=3-7 replicates run on mass spectrometry per timepoint; error is too small to visualize for a subset of timepoints; Welch’s t-test comparing bound and free at each timepoint; *p ≤ 0.05, **p ≤ 0.01, ***p ≤ 0.001, ****p ≤ 0.0001).

**Fig. S7.**
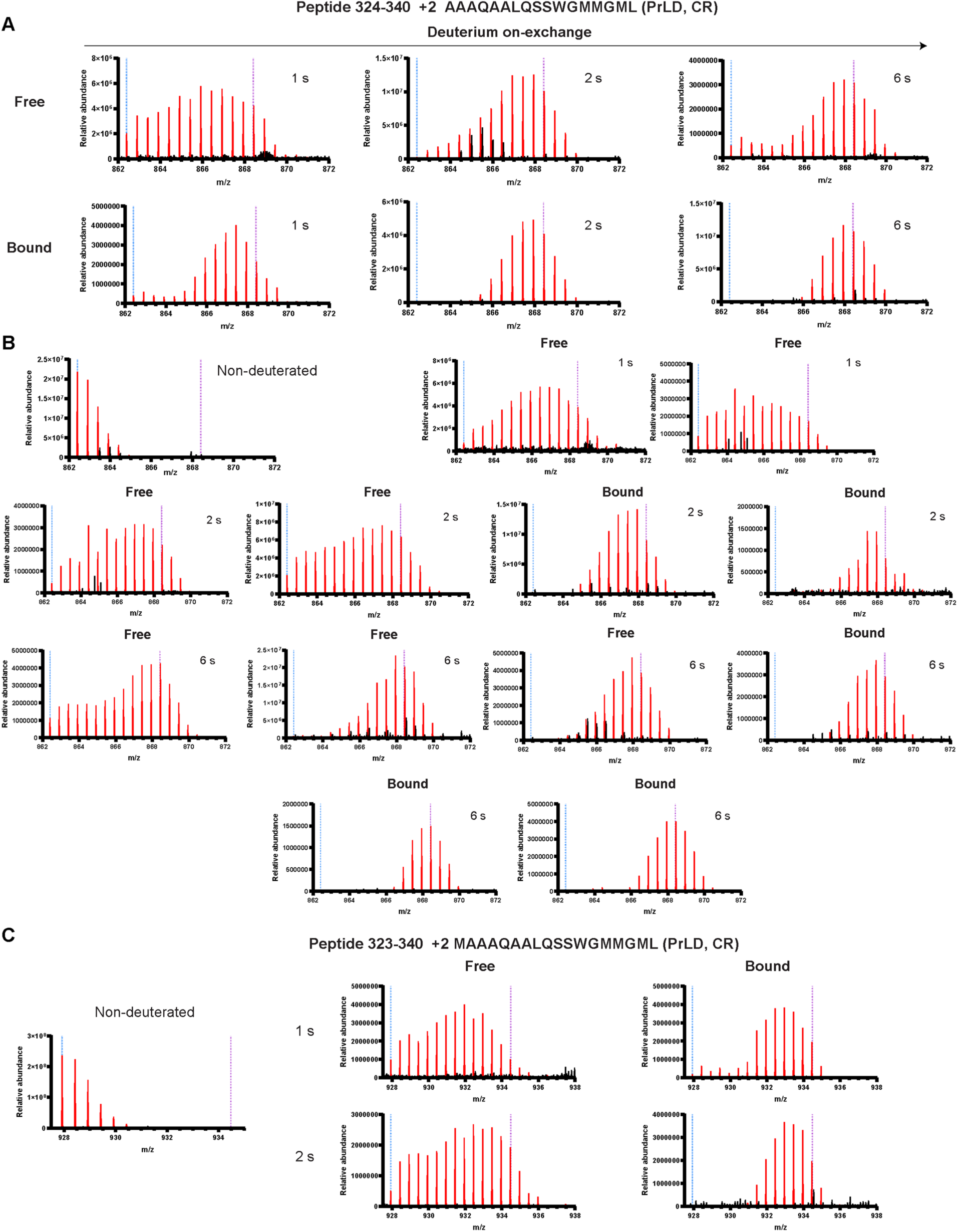
Mass spectra illustrate the effects of Clip34 RNA binding on TDP-43 structure. **(A-C)** Raw mass spectra data, with signal corresponding to the appropriate peptide as determined by appropriate m/z values colored red, and noise from overlapping peptide(s) colored black. The two dashed lines serve as visual guides; the blue dashed line indicates the monoisotopic peak, whereas the purple dashed line indicates the centroid value of the peptide in the fully deuterated sample. Spectra at 1 s, 2 s, and 6 s timepoints for the free and bound states for the indicated peptide in the CR of the PrLD, in addition to the representative spectra shown in Fig. 2G (A). The raw mass spectrum data of a representative non-deuterated sample, and remaining raw mass spectra data for replicates at 1 s, 2 s, and 6 s timepoints for the free and bound states of the indicated peptide, in addition to those shown in Fig. 2G and fig. S7A (B). The raw mass spectrum data of a representative non-deuterated sample, and representative spectra at 1 s and 2 s timepoints for the free and bound states for the indicated peptide in the CR of the PrLD (C).

**Fig. S8.**
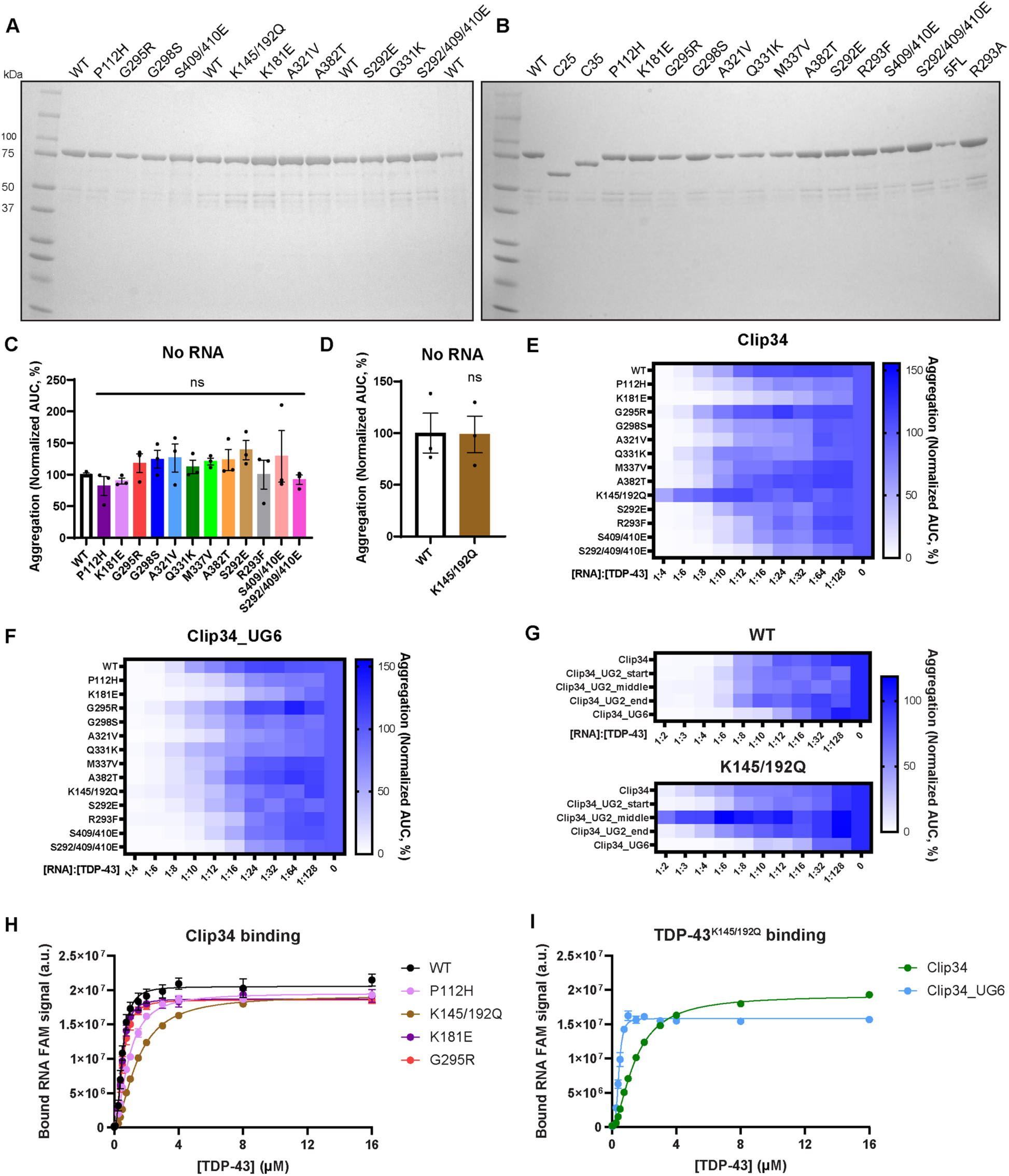
Clip34 and Clip34_UG6 effectively prevent aggregation of diverse disease-linked TDP-43 variants. **(A,B)** 4-20% Tris-HCl SDS-PAGE gels loaded with 1 µg of each indicated purified TDP-43 variant and subsequently stained with Coomassie Brilliant Blue. **(C, D)** AUC values for No RNA conditions for all examined TDP-43 variants, normalized to WT. For each graph, all data for each replicate was collected within the same experiment. Data are mean ± SEM (n=3; one-way ANOVA with Dunnett’s correction comparing to WT (C) or unpaired t-test (D)). **(E, F)** Heatmaps displaying the mean normalized AUC value at each tested molar concentration ratio of Clip34 (E) or Clip34_UG6 (F) to TDP-43, across individual replicates of aggregation assays. Data is normalized to the respective variant’s No RNA condition. This data was utilized to calculate the IC_50_ values depicted in Fig. 3, C and D. **(G)** Heatmaps displaying the mean normalized AUC value at each tested molar concentration ratio of Clip34 variants to WT TDP-43 or TDP-43^K145/192Q^, across individual replicates of aggregation assays. Data is normalized to the respective variant’s No RNA condition. This data was utilized to calculate the IC_50_ values depicted in Fig. 3, E and F. **(H)** Bound 5’ 6-FAM Clip34 signal for EMSAs performed with the indicated TDP-43-MBP-His variants. This is the summary data corresponding to K_D_ values calculated from individual replicates, shown in Fig. 3G. WT data is the same as shown in Fig. 1J and fig. S2D. Data are mean ± SEM (n=3; shown is the nonlinear regression: [agonist] vs response with variable slope, of the combined replicates). **(I)** Bound 5’ 6-FAM RNA signal for EMSAs performed with TDP-43^K145/192Q^. This is the summary data corresponding to K_D_ values calculated from individual replicates, shown in Fig. 3H. The Clip34 curve data is the same as shown in (H), included for reference. Data are mean ± SEM (n=3; shown is the nonlinear regression: [agonist] vs response with variable slope, of the combined replicates).

**Fig. S9.**
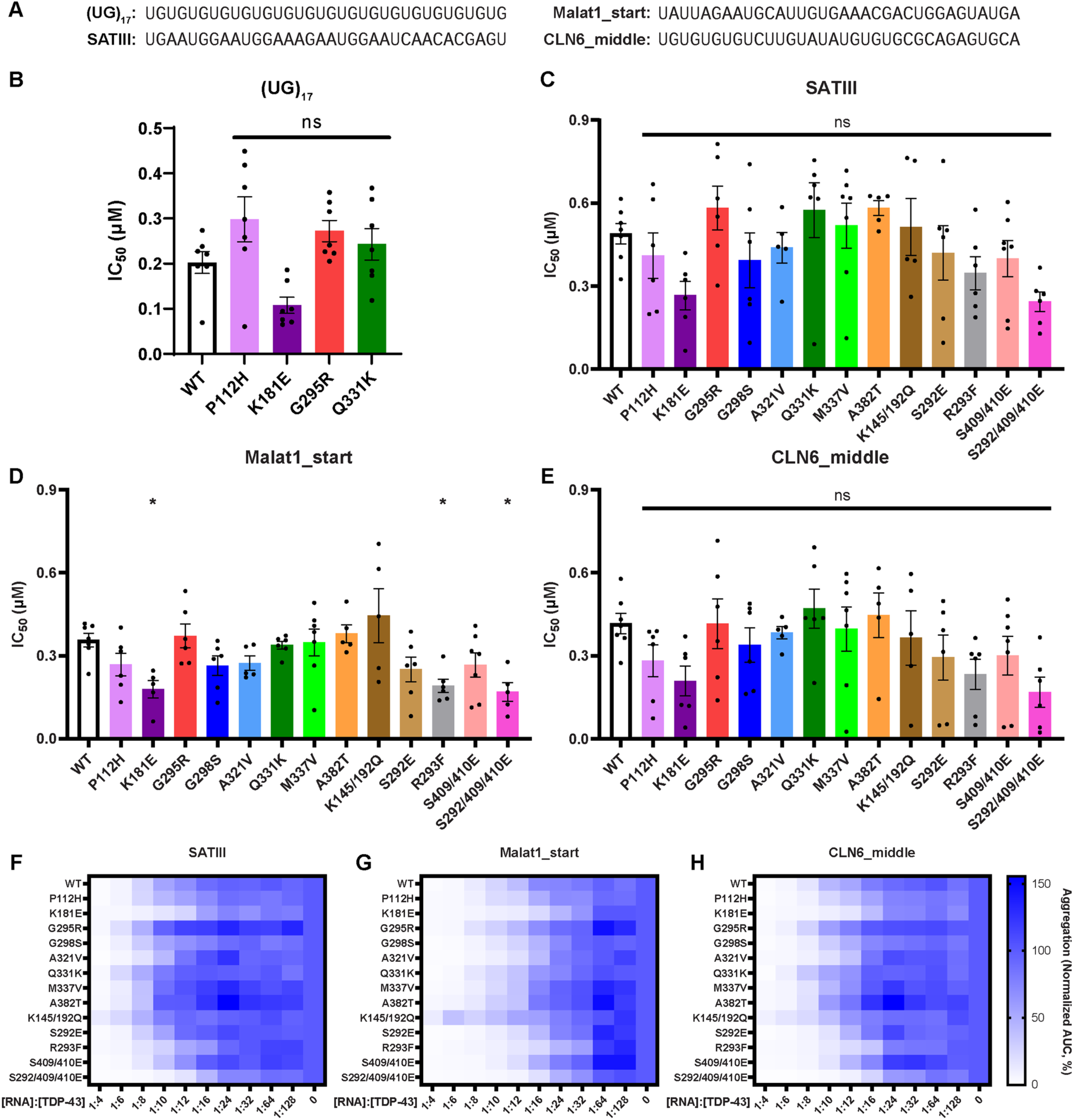
Malat1_start, SATIII, CLN6_middle, and (UG)_17_ effectively prevent aggregation of diverse disease-linked TDP-43 variants. **(A)** RNA sequences of tested RNAs. **(B)** IC_50_ values calculated for the indicated TDP-43 variants with (UG)_17_. Data are mean ± SEM (n=7; one-way ANOVA with Dunnett’s correction comparing to WT). **(C-E)** IC_50_ values for SATIII (C), Malat1_start (D), or CLN6_middle (E) RNAs with each TDP-43 variant. Data are mean ± SEM (n=5-7; one-way ANOVA with Dunnett’s correction comparing to WT; *p ≤ 0.05). **(F-H)** Heatmaps displaying the mean normalized AUC value at each tested molar concentration ratio of SATIII (F), Malat1_start (G), or CLN6_middle (H) RNAs to TDP-43, across individual replicates of aggregation assays. Data is normalized to the respective variant’s No RNA condition. This data was utilized to calculate the IC_50_ values depicted in (C-E).

**Fig. S10.**
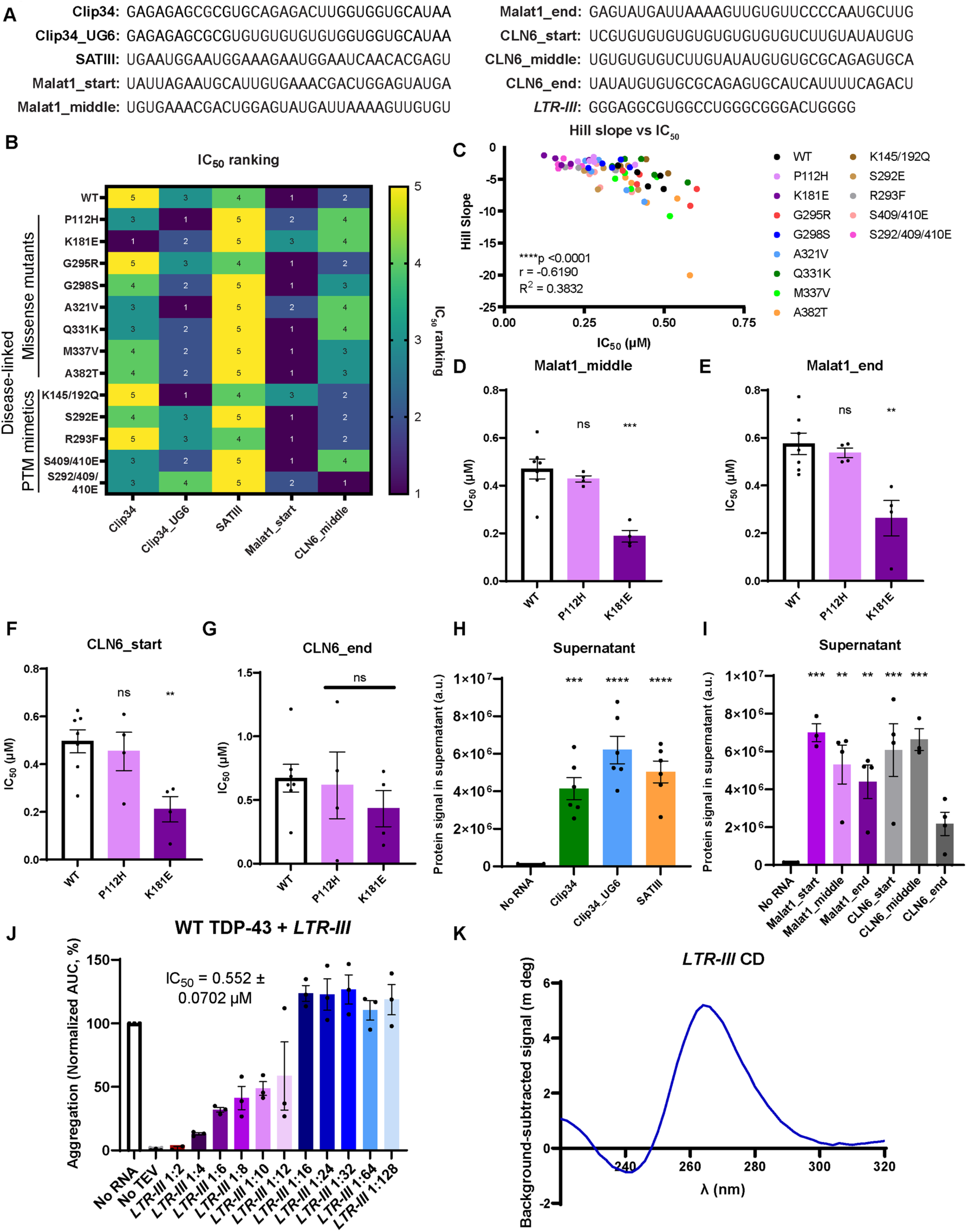
Several short RNA chaperones effectively prevent aggregation of diverse disease-linked TDP-43 variants. **(A)** RNA sequences of tested RNAs. **(B)** Heatmap displaying rankings of the IC_50_ data shown in Fig. 4B. Rankings of RNA IC_50_ values are determined for each TDP-43 variant, with a ranking of 1 corresponding to the “best” (lowest IC_50_ value), and a ranking of 5 corresponding to the “worst” (highest IC_50_ value). **(C)** The IC_50_ for each TDP-43 variant:RNA pair is plotted against the hill slope for that same pair, as calculated by nonlinear regression: [inhibitor] vs normalized response with variable slope, from the combined data of all replicates (Pearson correlation; ****p ≤ 0.0001). **(D-G)** IC_50_ values calculated for the indicated TDP-43 variants with Malat1_middle (D), Malat1_end (E), CLN6_start (F), or CLN6_end (G) RNAs. Data are mean ± SEM (n=4-7; one-way ANOVA with Dunnett’s correction comparing to WT; **p ≤ 0.01, ***p ≤ 0.001). **(H, I)** Quantification of the WT TDP-43 signal in the supernatant after sedimentation was performed at the end timepoint of aggregation assays. 4-20% Tris-HCl SDS-PAGE gels were loaded with equal volumes of supernatant from each sample, and subsequently stained with Coomassie Brilliant Blue. Replicates within each graph were collected within the same experiment. Data are mean ± SEM (n=3-6; one-way ANOVA with Dunnett’s correction comparing to No RNA; **p ≤ 0.01, ***p ≤ 0.001, ****p ≤ 0.0001). **(J)** AUC of turbidity data for WT TDP-43 with annealed *LTR-III* RNA, normalized to the No RNA condition. Data are mean ± SEM (n=3). **(K)** Circular dichroism spectrum for 5 µM *LTR-III* RNA after annealing.

**Fig. S11.**
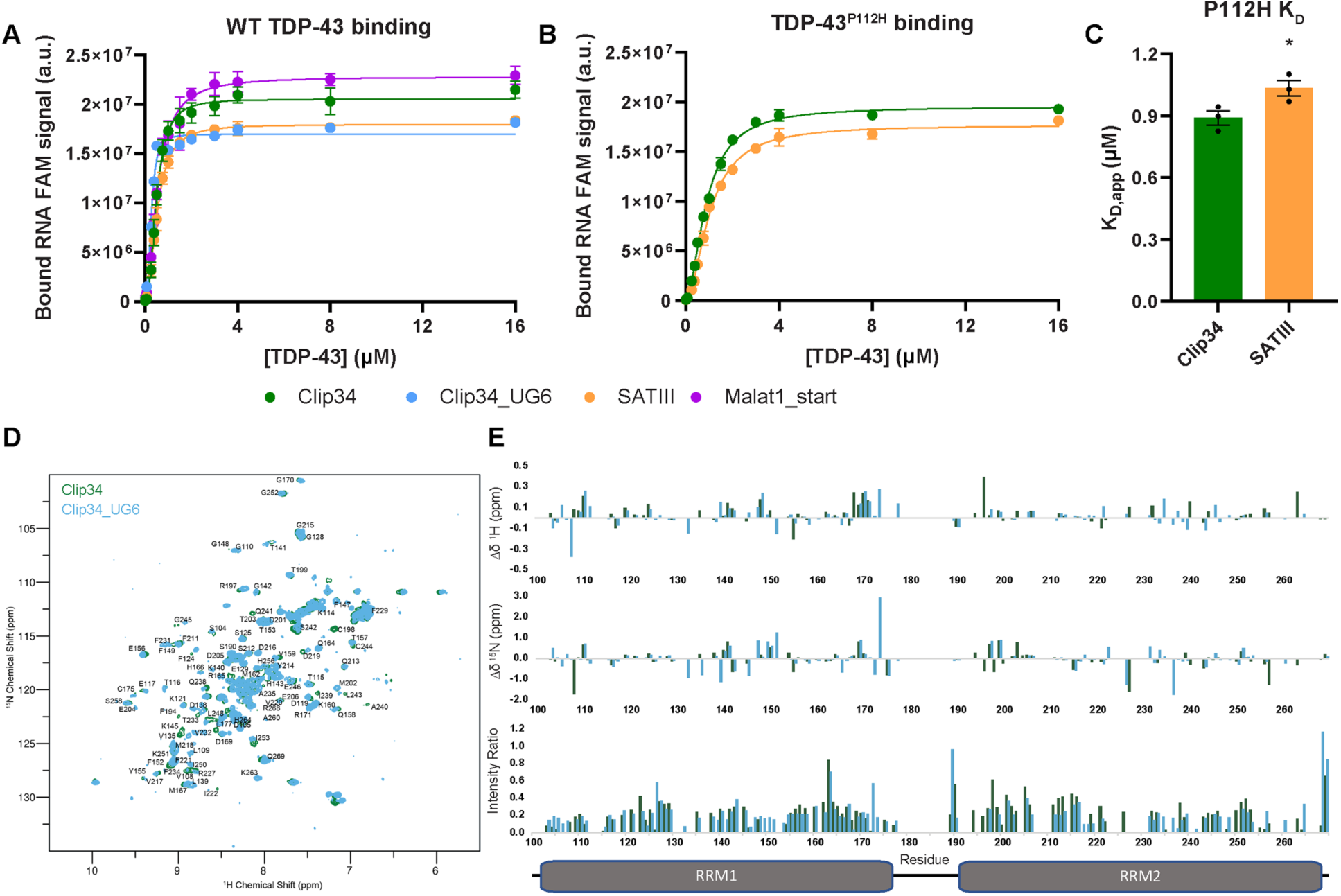
Short RNA chaperone engagement of the TDP-43 RRMs. **(A,B)** Bound 5’ 6-FAM RNA signal for EMSAs performed with WT TDP-43 (A) or TDP-43^P112H^ (B). The Clip34 curve data is the same as shown in Fig. 1J, fig. S2D, and fig. S8H, included for reference. Data are mean ± SEM (n=3-4; shown is the nonlinear regression: [agonist] vs response with variable slope, of the combined replicates). **(C)** Apparent K_D_ values calculated from bound 5’ 6-FAM signal of the indicated RNAs, from individual replicates of EMSAs performed with TDP-43^P112H^, corresponding to summary data shown in (B). Clip34 data is the same as shown in Fig. 3G. Data are mean ± SEM (n=3; unpaired t-test; *p ≤ 0.05). **(D)** Overlay of ^1^H-^15^N heteronuclear single quantum coherence (HSQC) spectra of TDP-43 RRMs with Clip34 RNA (green, 2:1::[Clip34]:[TDP-43]) and Clip34_UG6 RNA (blue, 2:1::[Clip34_UG6]:[TDP-43]). **(E)** ^1^H and ^15^N chemical shift perturbations (Δδ^1^H (*top*) and Δδ^15^N (*middle*), respectively), and intensity ratios (*bottom*) of TDP-43 RRMs upon binding of Clip34 (green) and Clip34_UG6 (blue) RNA. Domain map of TDP-43 RRMs shown at the bottom, aligned to x-axes of graphs.

**Fig. S12.**
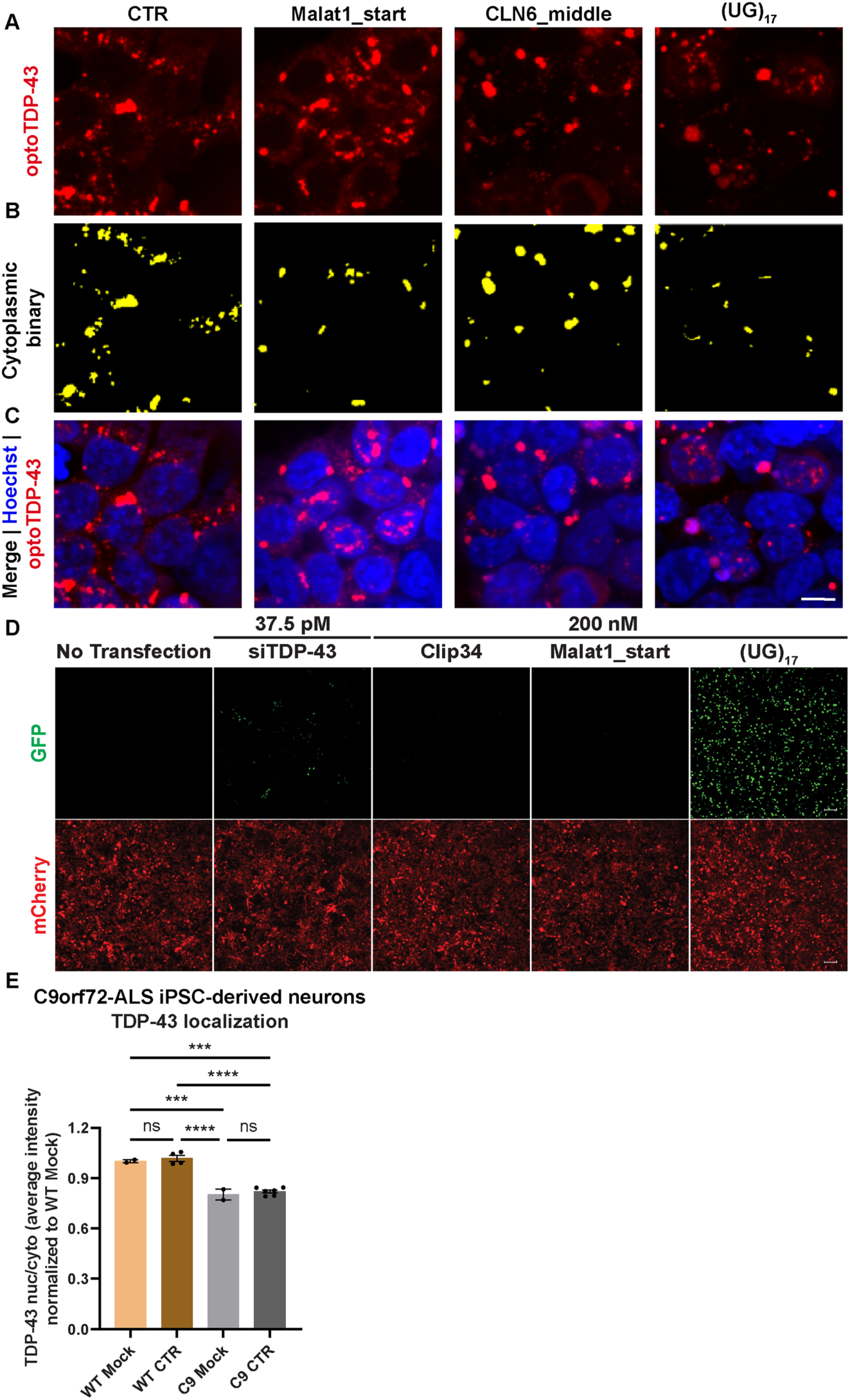
Clip34 and Malat1_start mitigate aberrant TDP-43 phenotypes in an optogenetic human cell model and in patient-derived neurons without interfering with TDP-43 function. (A-C) OptoTDP-43 stable HEK293 cells were treated with the indicated RNA, followed by blue light exposure to induce Cry2olig oligomerization, and imaged after fixation. Representative images corresponding to quantification in Fig. 5B, for optoTDP-43 (A), the binary signal of cytoplasmic optoTDP-43 puncta (B), and the merged image showing optoTDP-43 and Hoechst (C). Scale bar indicates 10 μm. **(D)** Stable HEK293 cells with inducible CUTS biosensor were transfected with 200 nM of Clip34, Malat1_start, and (UG)_17_ RNA oligos for 48 hours in doxycycline supplemented media (1000 ng/mL). The cells were also reverse transfected with a low dose (37.5 pM) siRNA of TDP-43 for 72 hours, together with 48 hours of doxycycline as a positive control. Cells experiencing TDP-43 loss of function will have green nuclei due to the expression of GFP-NLS that would ordinarily be repressed by a TDP-43-regulated cryptic exon (*82*). Live-imaging of CUTS HEK cells transfected with siRNA of TDP-43, Clip34, Malat1_start, and (UG)_17_. Green = GFP; red = mCherry. Scale bar indicates 100 µm. **(E)** The average ratio of TDP-43 nuclear to cytoplasmic signal normalized to healthy control iPSC-derived neurons without RNA treatment. C9 CTR condition data are the same as shown in Fig. 5D. Data are mean ± SEM (n=2-6 technical replicates; one-way ANOVA with Tukey’s correction; ***p ≤ 0.001, ****p ≤ 0.0001).

**Fig. S13.**
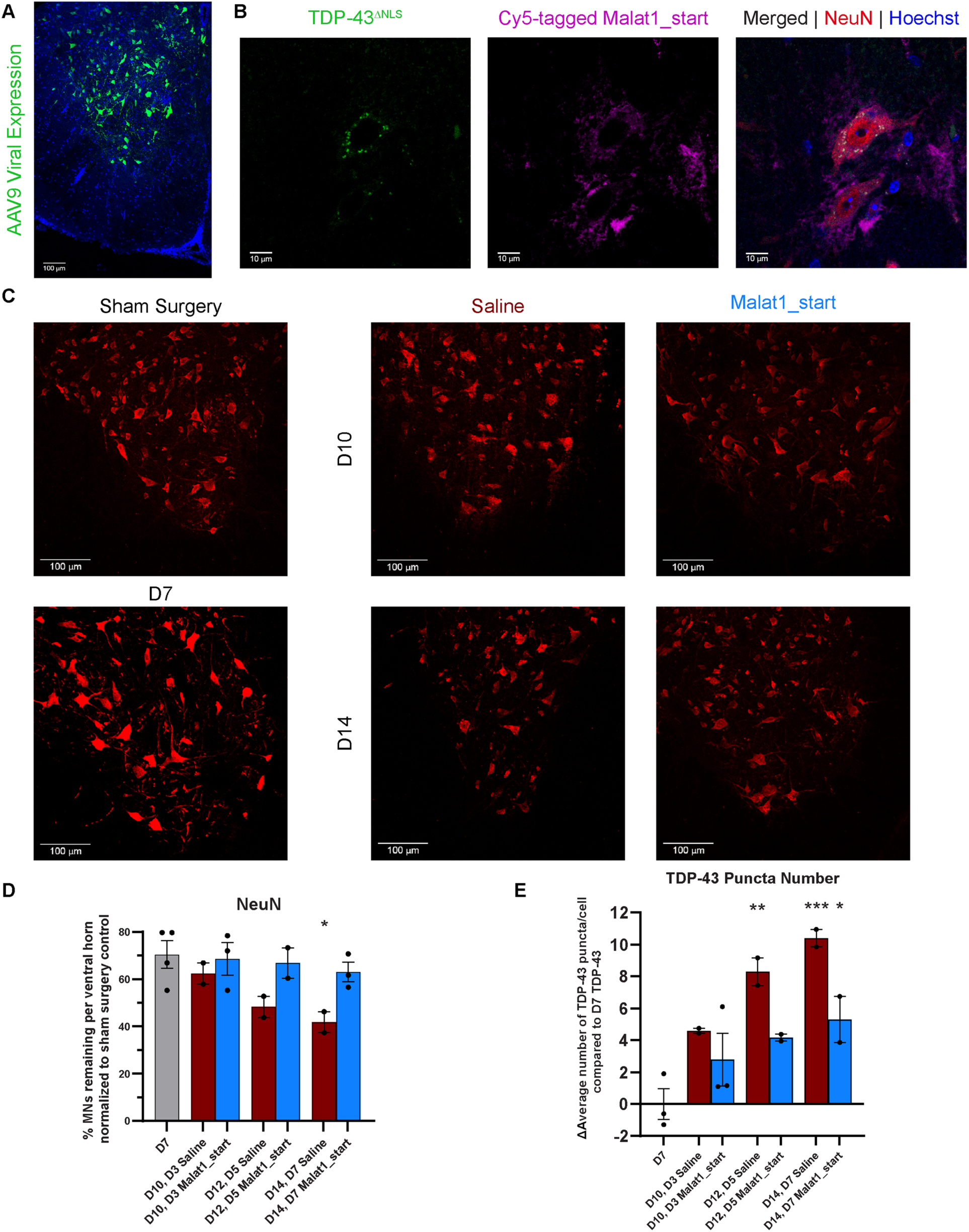
Malat1_start RNA mitigates neurodegeneration and TDP-43 aggregation in a mouse model of TDP-43 proteinopathy. **(A)** Representative transduction profile showing 10x magnification images following six 1×10^^11^ GC injections of AAV9-TDP-43^ΔNLS-YFP^ in p180 female mice. Shown here is baseline 7 days of expression, 30 µm spinal cord slice. Scale bar indicates 100 μm. **(B)** Representative image showing TDP-43 puncta (green), as well as Cy5-tagged Malat1_start RNA (magenta) in NeuN positive (red) motor neurons of the ventral horn of the spinal cord. Shown is a 5-day (12-day expressing) treated animal, demonstrating effective penetration of RNA to this region. 60x magnification; scale bar indicates 10 μm. **(C)** Representative immunohistochemistry images of ventral horns, stained for choline acetyltransferase (ChAT). 20x magnification z-stack confocal images; scale bar indicates 100 μm. **(D)** Immunostaining quantification for NeuN^+^ neurons manually counted within the ventral horn region of spinal cord sections. Data are mean ± SEM (n=2-4 animals per condition; n=4 ventral horns per animal; shown: one-way ANOVA with Dunnett’s correction comparing to D7 TDP-43: *p ≤ 0.05; not shown: two-way ANOVA on D10, D12, and D14 data: treatment (Malat1_start vs saline): *p ≤ 0.05). **(E)** TDP-43 positive puncta were assessed in ChAT^+^ motor neurons. Ten fields at 60x magnification were taken in the ventral horn for each animal (same fields as for Fig. 6E). For each animal, the average value of puncta per neuron was standardized to set the average D7 TDP-43 value to 0. Data shown are mean ± SEM (n=2-3 animals per condition; shown: one-way ANOVA with Dunnett’s correction comparing to D7 TDP-43: *p ≤ 0.05, **p ≤ 0.01, ***p ≤ 0.001; not shown: two-way ANOVA on D10, D12, and D14 data: treatment (Malat1_start vs saline): *p ≤ 0.05).

**Table S1.**
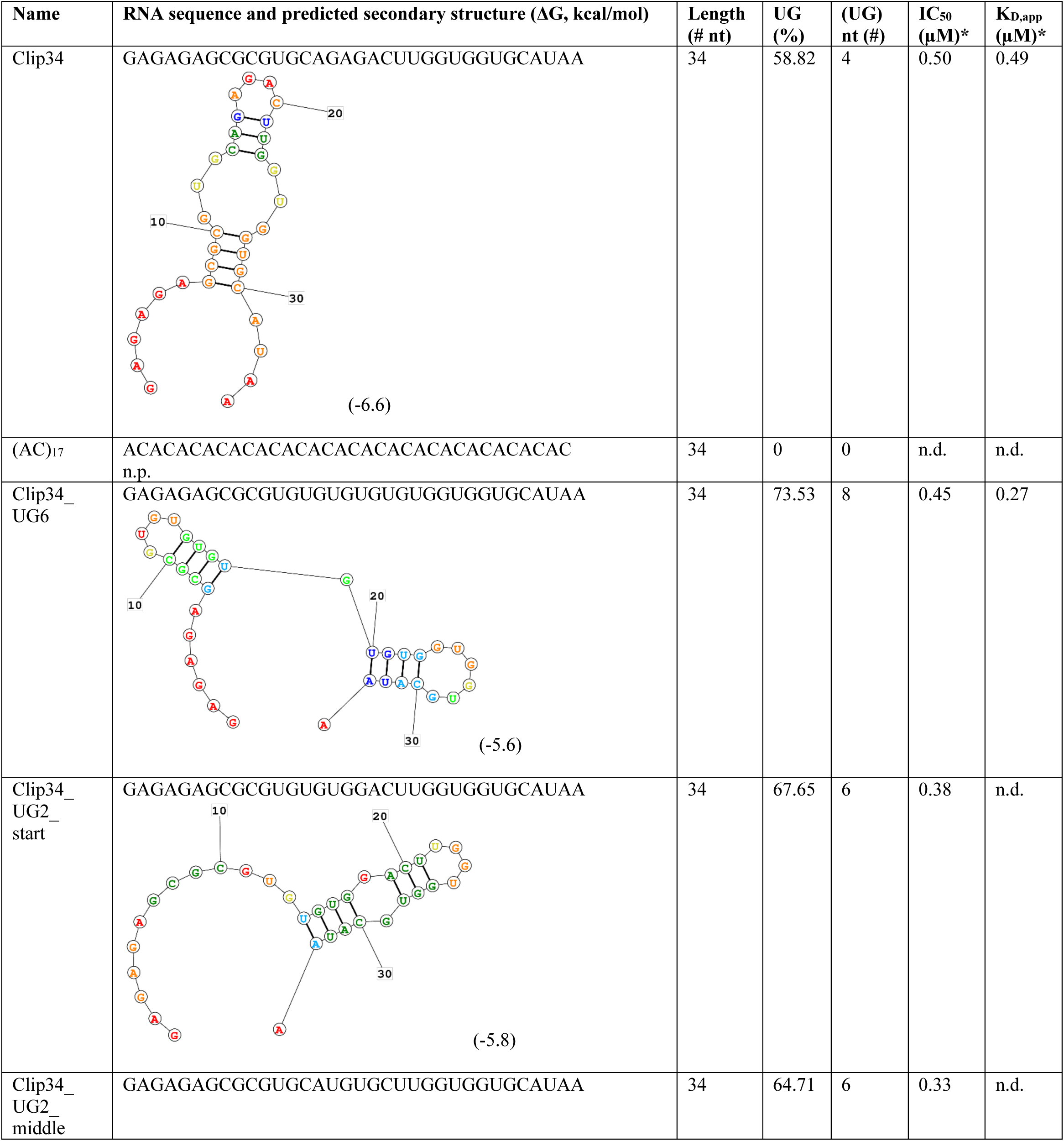
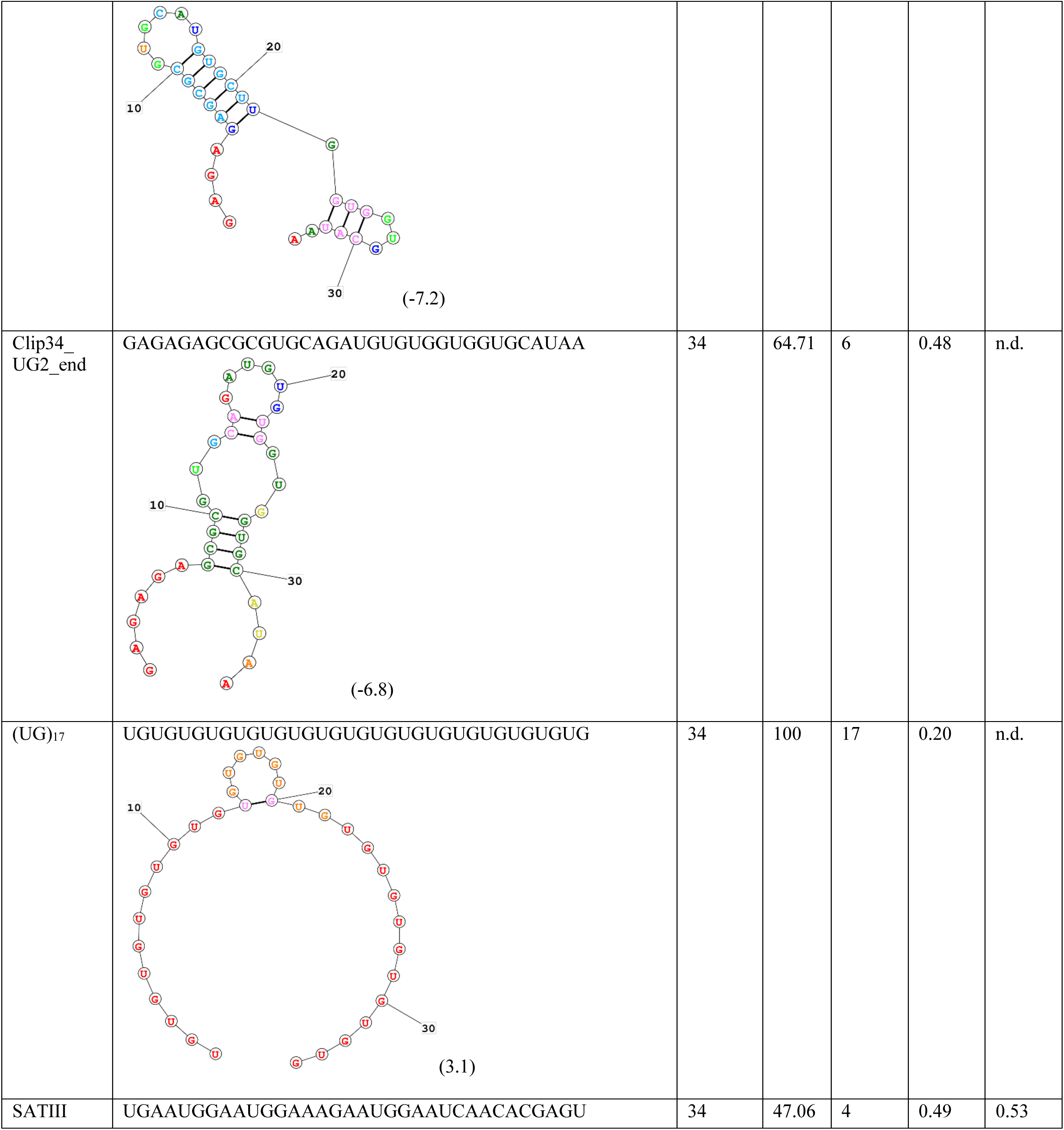
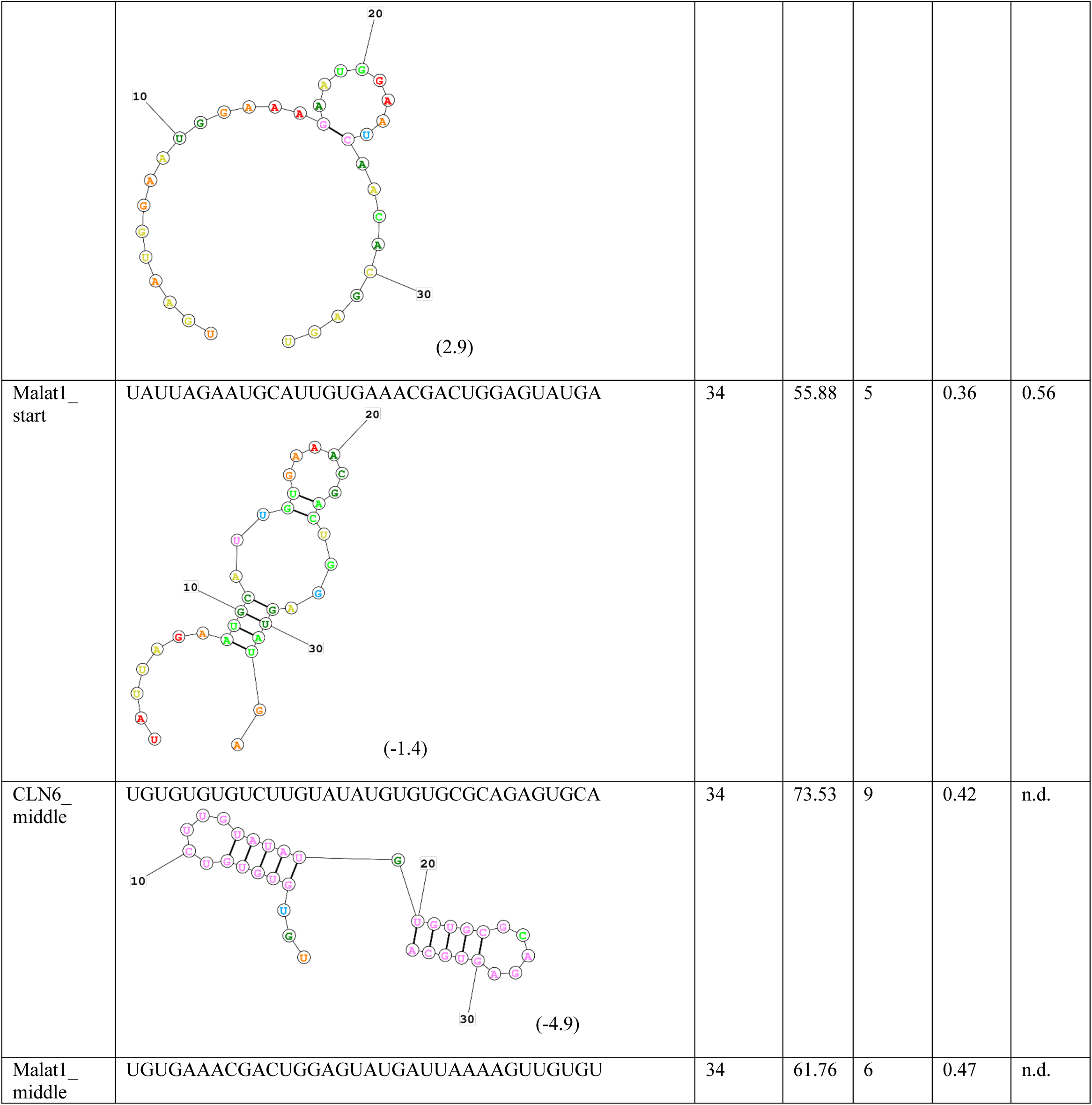
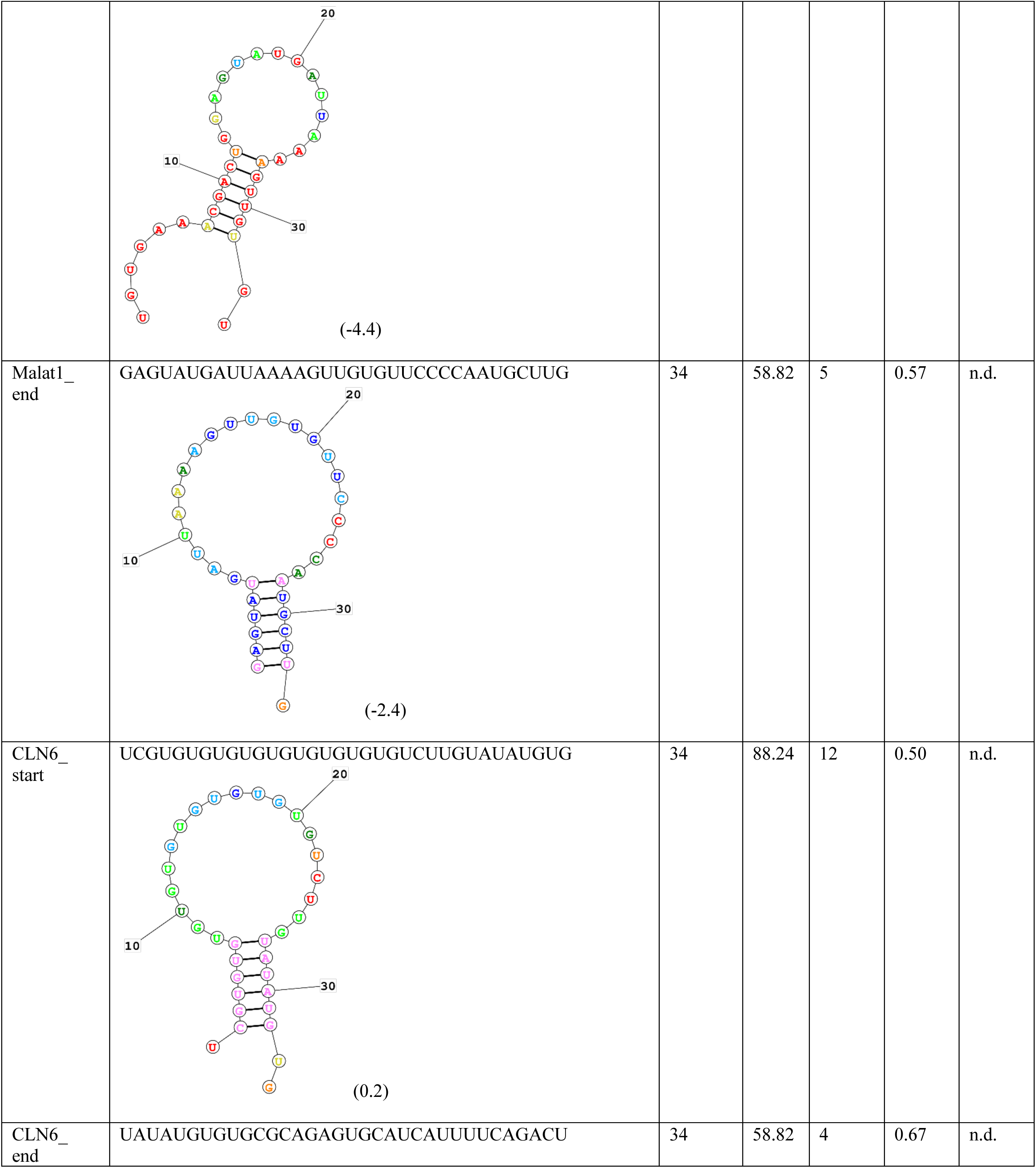
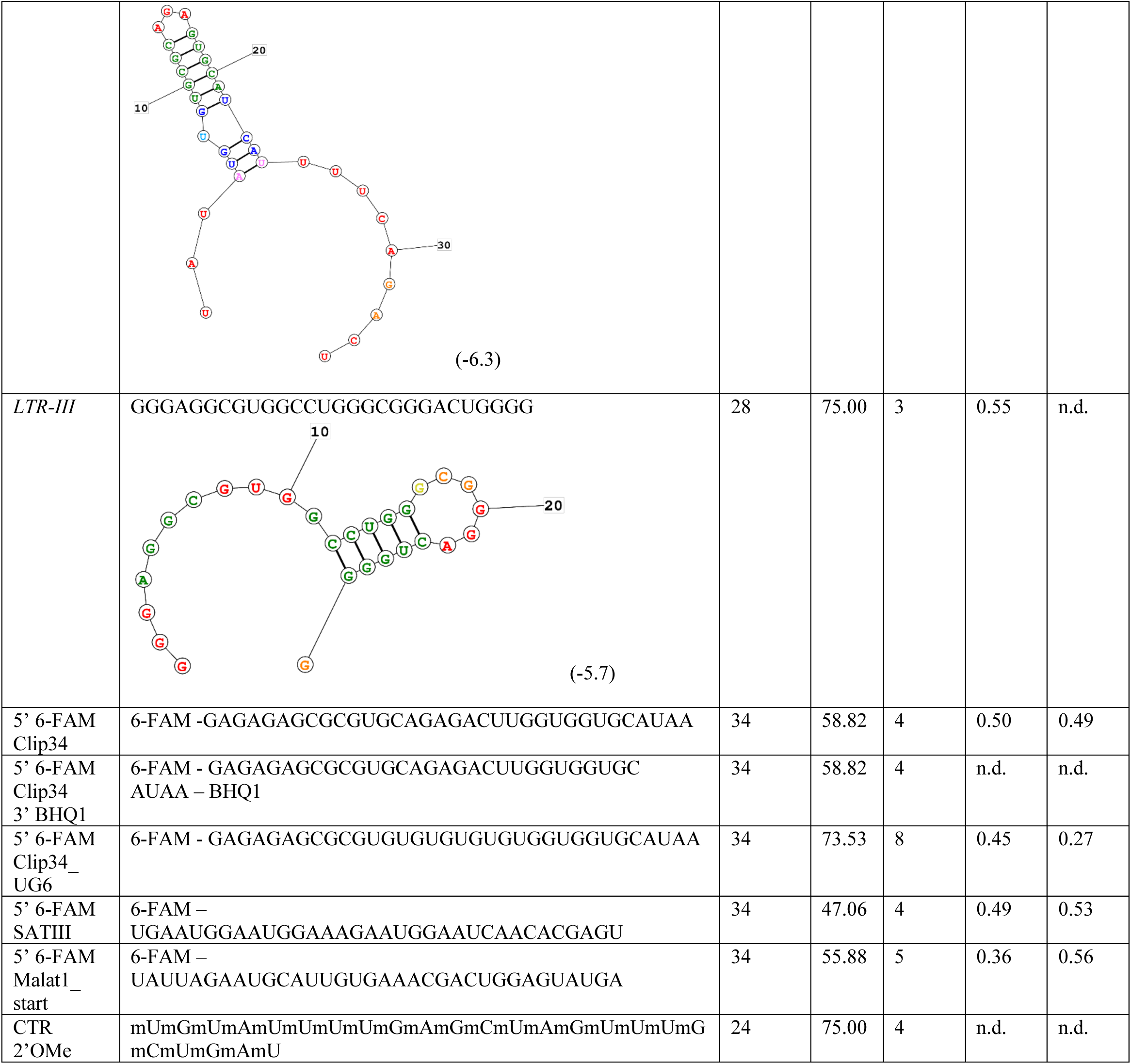
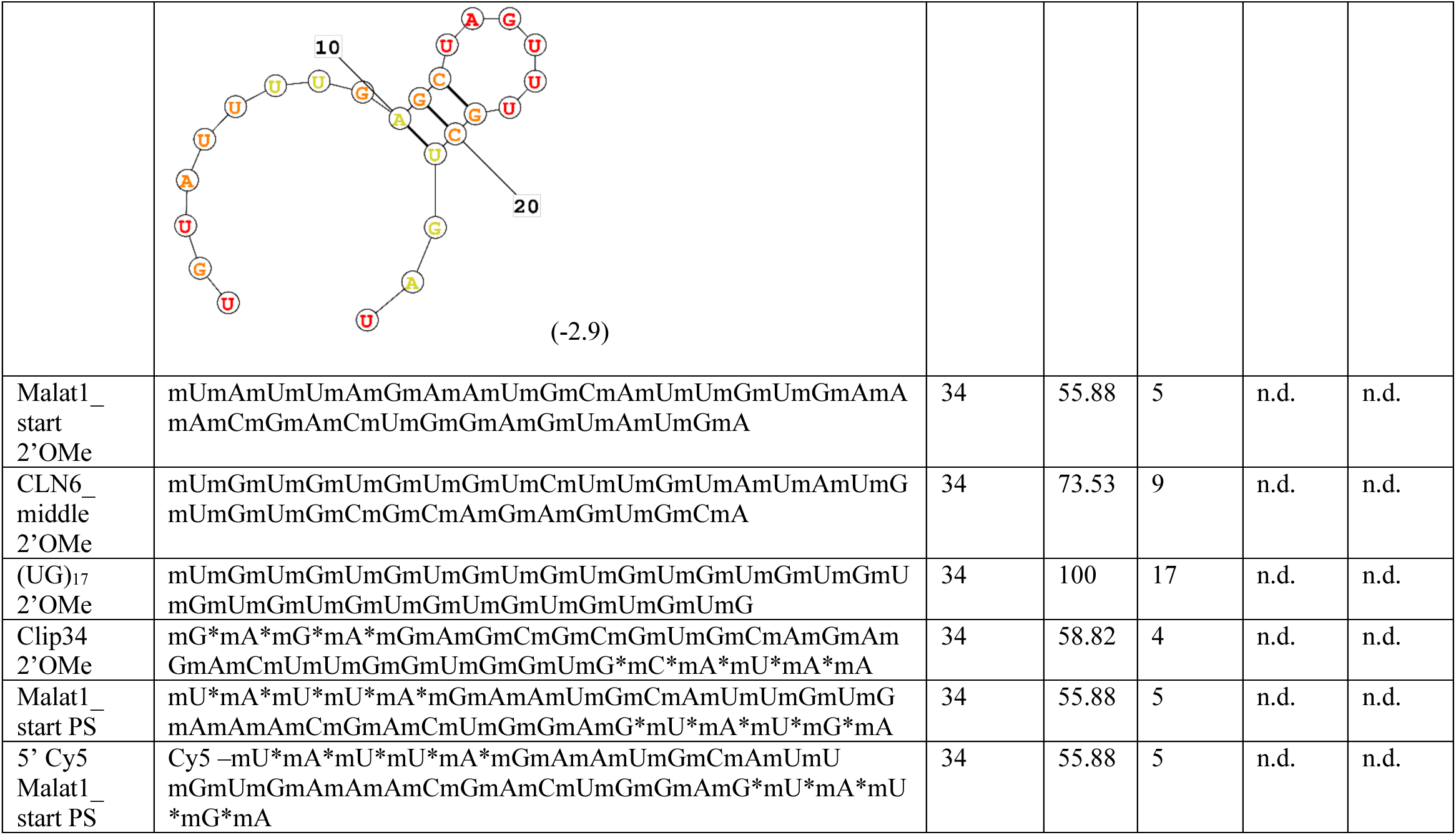
RNA oligonucleotides utilized in this study. The sequences of all RNA oligonucleotides used in this study are provided. ‘m’ indicates the nucleotide has a 2’-O-Methyl modification. ‘*’ indicates a phosphorothioate bond modification. RNA secondary structure predictions were performed utilizing the RNAstructure web server, and reported ΔG values (in kcal/mol) are listed. Nucleotide color corresponds to the confidence in prediction probability; red ≥ 99%; orange ≥ 95%; yellow ≥ 90%; dark green ≥ 80%; light green ≥ 70%; light blue ≥ 60%; dark blue ≥ 50%; pink < 50%. ‘n.p.’ indicates RNAstructure did not generate a predicted secondary structure. Structures were not repeated for the modified versions of unmodified RNAs. The length of each sequence is indicated, as well as two measures of the UG-richness of the sequence. UG (%) was calculated as the percentage of nucleotides in the sequence that are either U or G. UG (#) was calculated as the number of ‘UG’ dinucleotide repeats within the sequence. The rounded IC_50_ and K_D,app_ values for each RNA with WT TDP-43 are listed, referring to data shown in Fig. 3C-E, Fig. 4C, fig. S9B-E, and fig. S10D-G, J; RNAs for which these values were not determined are indicated as ‘n.d.’ *Both IC_50_ and K_D,app_ values are displayed for reference for both the unmodified and fluorophore-labeled versions of the same RNA sequence, although IC_50_ values were only calculated with the unmodified RNA and K_D,app_ values only with the fluorophore-labeled RNA.

**Table S2.** HXMS data summary. Data concerning HXMS experiments (*99*). Only peptides determined to have good signal were used for HX analysis; only these peptides were used to calculate the values for back-exchange, sequence coverage, average peptide length, redundancy, and repeatability. Back-exchange was determined including all charge states for each peptide. Average peptide length counts all amino acids in the peptide, including the two N-terminal amino acids and proline residues. Average peptide length was determined based on only unique peptide sequences; additional charge states for the same peptide sequence were excluded from the calculation. Redundancy was determined based on the total number of peptides used for HX analysis, including all charge states for each peptide. Redundancy was calculated as the total number of peptides multiplied by the average number of deuterons per peptide (peptide length excluding the two N-terminal amino acids and proline residues), divided by the total number of available amides in the protein (excluding the two N-terminal amino acids and proline residues). Repeatability was determined including all charge states for each peptide and was calculated based on percent deuteration values for each peptide (rather than for the absolute number of deuterons).

| **Table S2. HXMS data summary** |  |  |
| --- | --- | --- |
| **Dataset** | **TDP-43 (free)** | **TDP-43 + Clip34 RNA (bound)** |
| HX reaction details | 150 mM NaCl, 20 mM HEPES-NaOD, 1 mM DTT in D<sub>2</sub>O, pD<sub>read</sub> 6.0 or 7.0, at 25°C; 80% D<sub>2</sub>O | 150 mM NaCl, 20 mM HEPES-NaOD, 1 mM DTT in D<sub>2</sub>O, pD<sub>read</sub> 6.0 or 7.0, at 25°C; 80% D<sub>2</sub>O |
| HX time course | pD<sub>read</sub> 6.0: 1 s, 2 s, 6 s, 18 s, 1 min, 3 min. pD<sub>read</sub> 7.0: 20 s, 1 min, 3 min, 10 min, 30 min, 1.5 hr, 4.5 hr, 14.5 hr | pD<sub>read</sub> 6.0: 1 s, 2 s, 6 s, 18 s, 1 min, 3 min. pD<sub>read</sub> 7.0: 20 s, 1 min, 3 min, 10 min, 30 min, 1.5 hr, 4.5 hr, 14.5 hr |
| HX control samples | 6 Non-deuterated (ND) and 1 fully deuterated (FD; 30°C for 18 hr) | 6 Non-deuterated (ND) and 1 fully deuterated (FD; 30°C for 18 hr) |
| Back-exchange | Mean: 17%; interquartile range: 11% | Mean: 17%; interquartile range: 11% |
| Number of peptides | MS/MS identified 246 unique peptides. 137 peptides were found to have good signal and used for HX analysis (a subset of these peptides were analyzed at multiple charge states). Some peptides were not recovered at all timepoints. | MS/MS identified 246 unique peptides. 135 peptides were found to have good signal and used for HX analysis (a subset of these peptides were analyzed at multiple charge states). Some peptides were not recovered at all timepoints. |
| Sequence coverage | 92.7-97.1% (differences in sequence coverage at each timepoint are displayed in figS4A) | 87.9-97.1% (differences in sequence coverage at each timepoint are displayed in figS4B) |
| Average peptide length/redundancy | 15.8/5.24 | 15.9/5.20 |
| Replicates | 3 (1 s, 18 s, 20 s, 10 min, 30 min, 1.5 hr, 4.5 hr, 14.5 hr); 5 (2 s, 6 s); 6 (3 min; 3 replicates at each pD); or 7 (1 min; 4 replicates at pD 7.0, 3 at pD 6.0) | 3 (1 s, 18 s, 20 s, 10 min, 30 min, 1.5 hr, 4.5 hr, 14.5 hr); 5 (2 s, 6 s); 6 (1 min, 3 min; 3 replicates at each pD) |
| Repeatability | 2.67% (average standard deviation from replicate measurements of percent deuteration of peptides at 10 min timepoint) | 1.34% (average standard deviation from replicate measurements of percent deuteration of peptides at 10 min timepoint) |
| Significant differences in HX | Unpaired t-test with Welch's correction of a difference between free and bound states using p-value < 0.05 | Unpaired t-test with Welch's correction of a difference between free and bound states using p-value < 0.05 |

